# Cross-species biomarkers of cerebellar injury in acute sepsis: integrated metabolomics of mouse cerebellum and plasma identify conserved human plasma biomarkers and therapeutic modulation by human MSC-sEVs

**DOI:** 10.1101/2025.06.26.660174

**Authors:** Dylan W. Crawford, Maria Triantafyllou, Nora Wolff, Marc Carlson, Chad Byrd, David R. Graham, Panagiotis Kratimenos, Vittorio Gallo, Ioannis Koutroulis

## Abstract

Septic encephalopathy (SE) is a severe complication of sepsis, characterized by neuroinflammation and metabolic dysfunction, with the cerebellum among the most vulnerable brain regions. Advances in the field have been constrained by the lack of reliable biomarkers for SE detection, incomplete mapping of cerebellar metabolic alterations in SE, and limited insight into the therapeutic mechanisms of human mesenchymal stem cell (MSC)-derived small extracellular vesicle (sEV) therapy. To overcome these challenges, we used a murine sepsis model and conducted integrated metabolomic analyses of cerebellar tissue and plasma, with and without MSC-sEV administration. Cross-compartment analyses identified six plasma metabolites with strong diagnostic potential in mice, three of which (n-acetylputrescine, aspartic acid, and cystathionine) were also observed in the plasma of human septic patients, supporting their promise as translatable biomarker candidates. Sepsis triggered profound cerebellar metabolic dysfunction, suppression of oxidative energy metabolism, and redox imbalance. MSC-sEVs attenuated these disturbances via their bioactive cargo, restoring cellular energetics and reestablishing antioxidant balance. Collectively, these results highlight cross-species plasma biomarkers for SE diagnosis, delineate key cerebellar metabolic mechanisms in SE, and demonstrate therapeutic modulation by human MSC-sEVs.

## INTRODUCTION

Sepsis is a major global health concern, responsible for nearly 20% of deaths worldwide¹ and disproportionately affecting children and underserved populations². It arises from a dysregulated host response to infection, marked by an imbalance between hyperinflammation and immunoparalysis¹,³. One serious consequence is septic encephalopathy (SE), characterized by neuroinflammation and metabolic dysfunction in the brain,, leading to central nervous system impairments, including cognitive and motor deficits in survivors. The cerebellum is especially vulnerable, with damage driven by immune-mediated neuroinflammation and neuronal death . Multiple studies, including our recent work, have shown that sepsis-associated energy deficits contribute to cerebellar and broader brain dysfunction, a defining feature of SE .

A major barrier to progress has been the lack of accessible, reliable biomarkers for SE. Metabolomics enables quantification of metabolites within biological systems and links them to disease states and treatment responses, . Since sepsis disrupts cellular energetics and alters metabolic pathways, metabolomic analysis offers a powerful approach to identify metabolic fingerprints associated with sepsis-induced adaptations. It can also reveal metabolites involved in recovery following interventions designed to restore energy balance. By capturing a broad spectrum of endogenous metabolites, metabolomics provides insights into gene activity, enzyme function, and physiological status at the time of sampling, yielding a more precise metabolic snapshot than conventional biomarkers,¹ . This makes it a strong platform for identifying biomarkers and therapeutic targets to improve diagnosis, prognosis, and treatment of sepsis-related conditions. However, most metabolomics studies in sepsis have focused on diffuse compartments such as plasma or urine, limiting discovery of organ- or region-specific signatures (e.g., cerebellum). Identifying organ-related metabolic changes that also appear in accessible compartments could provide minimally invasive markers of early disruption, with major diagnostic implications. In addition, few studies have examined cross-species metabolomic conservation, a critical step for validating experimental biomarkers and advancing them toward clinical translation.

Beyond biomarker discovery, metabolomics also helps define the energetic disruptions underlying SE. Sepsis shifts energy production from efficient mitochondrial oxidative phosphorylation (OXPHOS) to less efficient glycolysis¹¹. Reduced ATP production leads to energy deficiency, oxidative stress, and accumulation of reactive oxygen species (ROS), ultimately driving apoptosis and organ failure¹². Emerging evidence suggests that these metabolic deficits actively contribute to immune-mediated neuroinflammation and neuronal death¹³,¹, though the relationship between immunometabolic dysfunction and SE pathophysiology remains incompletely understood.

Therapies addressing both immune dysfunction and energy deficits remain limited. One promising approach involves adipose-derived mesenchymal stem cell (MSC)-derived small extracellular vesicles (sEVs), which act on both immune and metabolic pathways. In murine sepsis models and other related conditions such as acute respiratory distress syndrome and bacterial pneumonia¹, MSC-sEVs have shown beneficial effects on immunometabolism. These findings emphasize the dual importance of immune and metabolic regulation in sepsis outcomes and suggest therapeutic potential. In our prior work, MSC-sEVs reduced SE severity, partially restored aerobic metabolism, and lowered pro-inflammatory cytokines in the cerebellum . Yet critical gaps remain regarding mechanisms of injury, the biological pathways involved, and the identity of clinically relevant biomarkers.

MSC-sEVs exert their effects primarily through bioactive cargo, including microRNAs (miRNAs)— such as miR-21, miR-146a-5p, and miR-199a, all linked to improved sepsis outcomes¹,¹ —and proteins including Annexin A1, ATP6V1A, and valosin-containing protein (VCP). Other sEV miRNAs, such as miR-30, Let-7, miR-15, and miR-338, regulate mitochondrial dynamics and OXPHOS, while miR-30 also maintains lipid metabolism homeostasis¹,¹ –²¹. These factors, found abundantly in MSC-sEVs and activated in cerebellar tissue of treated mice, represent strong candidates for driving their therapeutic mechanisms.

Despite these advances, three major knowledge gaps persist: (1) the absence of plasma-detectable biomarkers of cerebellar injury, limiting early clinical intervention; (2) incomplete characterization of metabolic disruption in SE; and (3) unclear mechanisms by which human MSC-sEVs restore metabolic function. To address these gaps, we applied targeted and untargeted metabolomics to cerebellar tissue and plasma from a polymicrobial cecal slurry mouse model of sepsis. First, we compared metabolomic responses across compartments to identify plasma metabolites with diagnostic potential for early SE detection, and then assessed their cross-species consistency in human sepsis plasma samples. Second, we defined metabolomic signatures of acute sepsis in cerebellar tissue. Finally, we evaluated how human MSC-sEV treatment modifies these profiles and integrated our findings with existing knowledge of MSC-sEV cargo to propose mechanistic pathways.

By combining advanced metabolomics with bioinformatics across tissues, plasma, and species, this study identifies novel biomarkers that are both mechanistically informative and clinically relevant for SE diagnosis, monitoring, and patient stratification. In parallel, it defines key biochemical pathways underlying disease progression and therapeutic response.

## MATERIALS AND METHODS

### Subjects

A total of 62 wildtype C57BL/6 male mice (Charles River Laboratories, strain code: 027 for cerebellum and Jackson Laboratories strain code: 000664 for plasma) aged 8-12 weeks served as the experimental subjects. Age-matched wildtype C57BL/6 male mice served as donor animals for cecal slurry preparation. All mice were maintained in the animal facility of the Children’s National Hospital and handled in accordance with the Institutional Animal Care and Use Committee (IACUC #00030714) of Children’s National Hospital, who approved all the protocols and the Guide for the Care and Use of Laboratory Animals (National Institute of Health). All mice were maintained in the animal facility under a 12-hour dark-light cycle, consistent temperature (20-26 °C) and humidity (40-60%), and *ad libitum* access to food and water.

### Sepsis induction via intraperitoneal cecal slurry intraperitoneal injection and MSC-sEV treatment administration via intravenous injection

Polymicrobial sepsis was induced using a well-established and extensively studied protocol^22^ commonly used in our laboratories^5^. This model was chosen for its high level of reproducibility and ability to closely model the pathophysiology observed in human sepsis cases while avoiding additional undue surgical trauma or tissue damage concerns stemming from alternative methods such as traditional cecal ligation and puncture models^23^.

Donor mice were euthanized and cecal contents were extracted and diluted with 5% dextrose to produce a cecal slurry with a concentration of 80 mg cecal contents/mL. Experimental mice then received an intraperitoneal (IP) injection of cecal slurry at a dose of 0.9 mg/g body weight, while controls received equivalent volume of phosphate-buffered saline (PBS).

Following a six-hour cecal slurry incubation period, a single dose of human adipose tissue MSC-derived sEVs or sEV-depleted media (used for MSC culture) was administered intravenously via lateral tail vein injection^5^. MSC-sEVs were prepared at a concentration of 10E+08 to 10E+09 particles in 250 µL by Zen-Bio (Durham, NC).

### Experimental design

Experimental mice were randomly assigned to one of four groups, as previously described^5^ (see **Figure 1a**):

1. **Control (-)** mice received IP injection of PBS and tail vein injection of sEV-depleted media.
2. **Control (+)** mice received IP injection of PBS and tail vein injection of MSC-derived sEVs.
3. **Sepsis (-)** mice received IP injection of cecal slurry and tail vein injection of sEV-depleted media.
4. **Sepsis (+)** mice received IP injection of cecal slurry and tail vein injection of MSC-derived sEVs.

**Figure 1.**
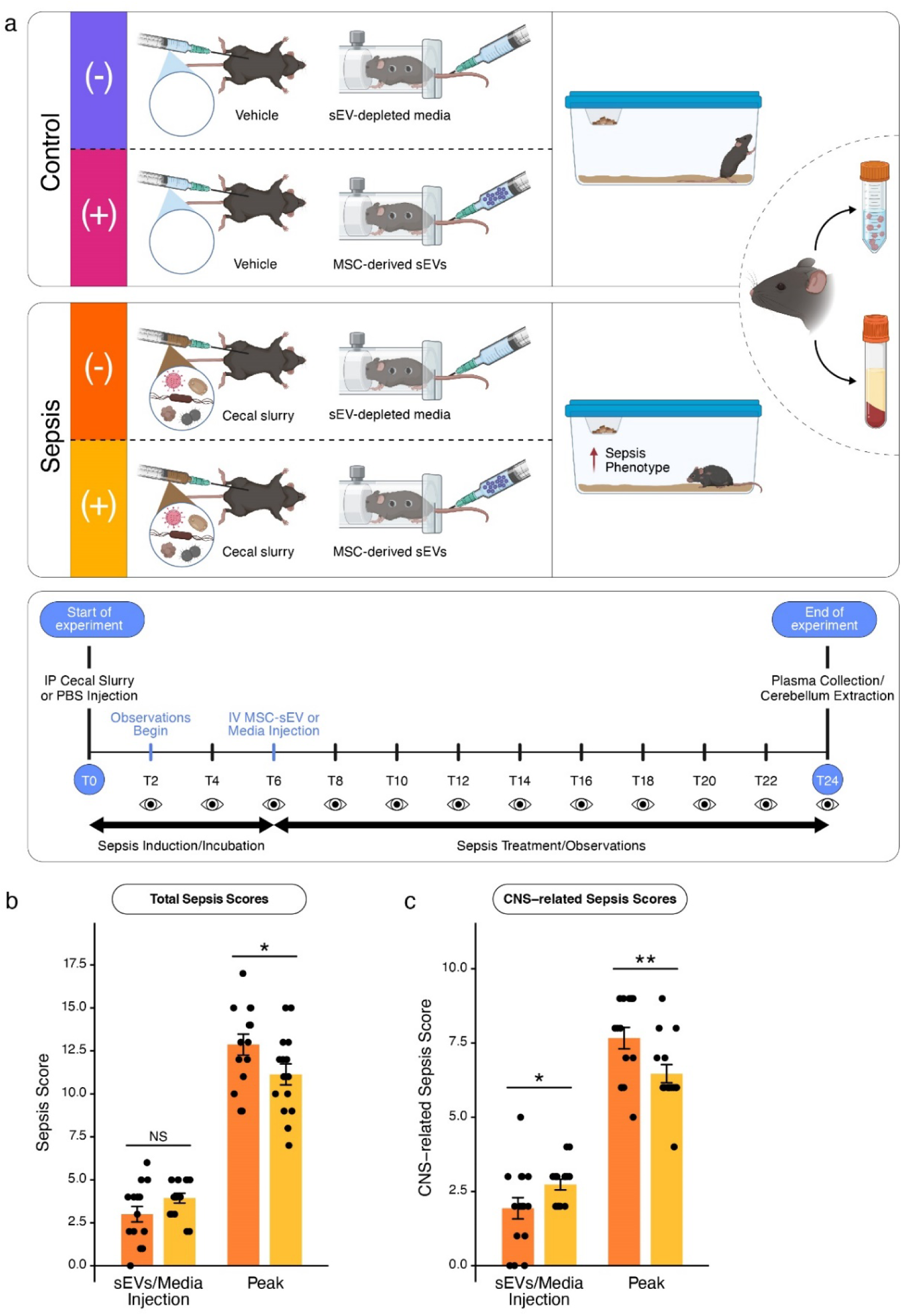
Overview of experimental design and clinical scoring. (**a**) Schematic and experimental timeline of the cecal slurry model of sepsis. This panel outlines the nature of the four experimental groups, their treatments, and the timeline on which sepsis is inducted and observed followed by sample collection. Schematic created in BioRender. Crawford, D. (2025) https://BioRender.com/y63q355. (**b**) Group means and variability in average total clinical sepsis scores for untreated, Sepsis (-), and treated, Sepsis (+), animals at the time of treatment administration and at peak observed illness. (**c**) Group average scores based on a subset of total clinical sepsis scores which reflects a CNS-specific phenotypes. Control groups are not shown in the plots, as scores remained negligible (≤ 0.5) across both measures at both timepoints. Data represented as mean ± SEM, Independent samples t-test. NS: non-significant, * denotes p ≤ 0.05, ** denotes p-value ≤ 0.01.

**Figure 2.**
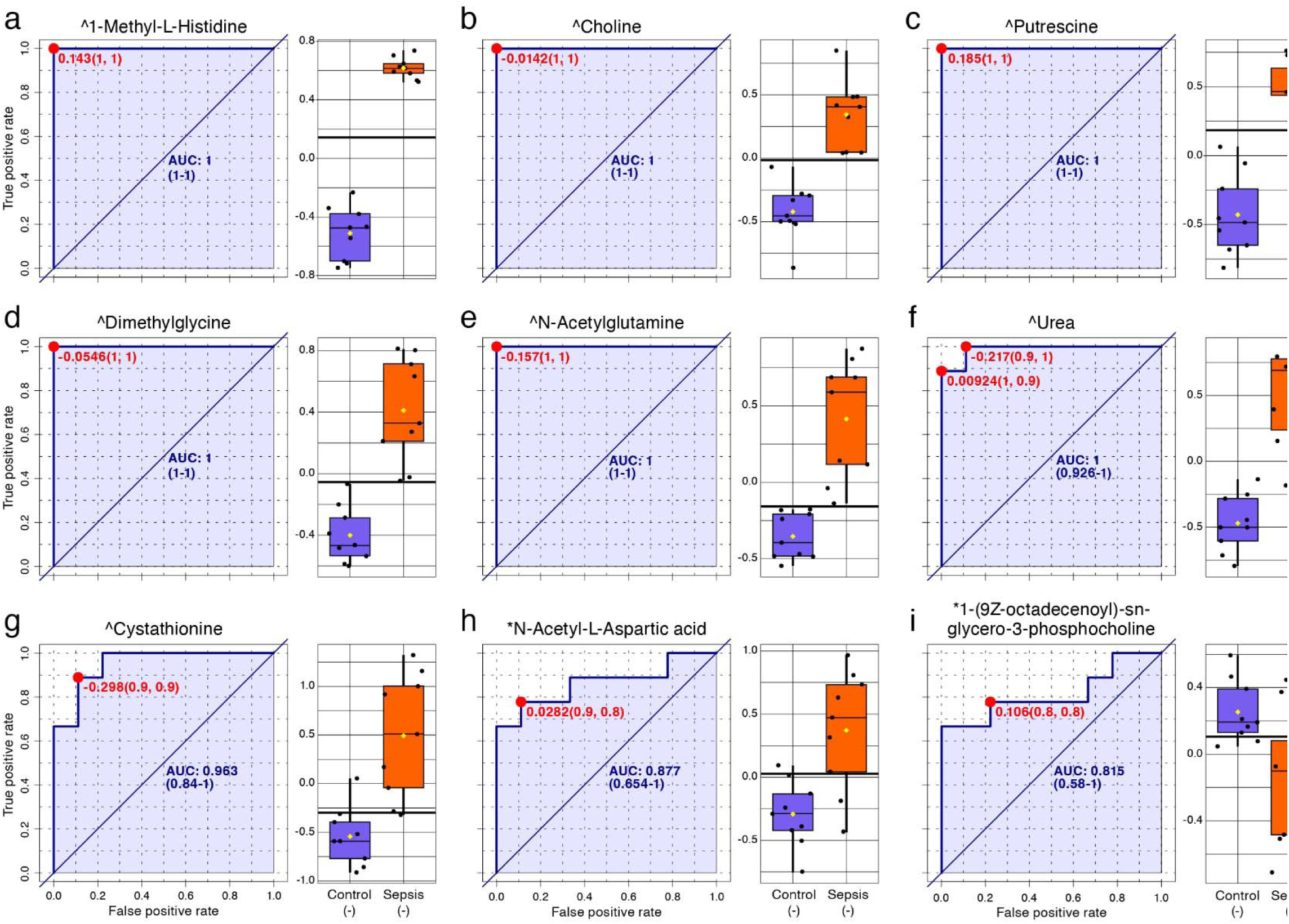
Overlapping biomarker candidates with high diagnostic performance in both cerebellum and plasma. This figure displays all nine metabolite features with strong diagnostic potential (AUC ≥ 0.8) that were altered in both cerebellar and plasma samples considering the disease state comparison, Sepsis (-) v. Control (-). Panels **a-i** display receiver operating characteristic (ROC) curves for each overlapping metabolite candidate, with AUC values shown to reflect classification performance. Box plots are displayed to the right of each metabolite’s ROC curve, illustrating the relative average concentrations of each metabolite by condition. Where known, the best diagnostic threshold is displayed by a black horizontal line on the box plots. Caret symbols (^) denote a Level 1-2 confidence identification from core facility panel confirmed by reference standard; Asterisks (*) denote a putatively annotated feature from LC-MS/MS fragmentation matches using the NIST MS Search software, earning Level 3+ identification confidence^25^.

Following IP injection in-line with group assignment, all mice were subject to evaluations every two hours for the subsequent 24-hours. Evaluations included the tracking of total and CNS-specific sepsis severity progression using a validated murine sepsis severity rubric developed by Shrum et al^24^. Mice were sacrificed 24-hours post-IP injection. For ethical reasons, mice reaching a total sepsis score ≥ 15 (on a scale from 0 to 28) before the 24-hour timepoint were sacrificed immediately and, thus, excluded from the study.

### Tissue and Plasma Extraction

Cerebellar tissue was harvested from 32 mice following perfusion with PBS at the time of euthanasia. Harvested tissue was stored at −80°C until submission for mass spectrometry analysis.

Whole blood was collected intracardially from 29 mice during perfusion while anesthetized with isoflurane. Blood was drawn into 200 µL microvette tubes containing potassium EDTA (Kent Scientific, Torrington, CT). Samples were centrifuged at 2000 g for 15 minutes to obtain platelet-poor plasma, which was then stored at −80°C until submission for mass spectrometry.

Briefly, tissue or plasma metabolites were extracted on ice using a biphasic solvent mix (H_2_0/MeOH/IPA) that included two internal standards to ensure extraction and system reproducibility. Acetonitrile was added for protein precipitation and samples were then configured and clarified supernatants were stored for subsequent analysis by LC-MS or LC-MS/MS analysis.

### Targeted Metabolomics (Panel)

The targeted method includes approximately 270 endogenous molecules with selective reaction monitoring transitions generated using authentic chemical standards and retention times using QTRAP 5500 LC-MS/MS System (Sciex) operating in both positive and negative ion modes. Five microliters of the extracted supernatants from samples were injected chromatographically resolved using a Kinetex 2.6 μm (Phenomenex) column at 0.2 mL/min flow rate with a binary gradient of water and acetonitrile (both containing 0.1% formic acid).

Intensities were normalized to respective internal standard area and processed using MultiQuant 3.0.3. The quality and reproducibility of LC-MS data were ensured using pooled quality control (QC) samples, National Institute of Standards and Technology (NIST) plasma samples, and blank solvent runs between sets of samples to minimize carry-over effects.

### Untargeted Metabolomics

Untargeted metabolomics analysis was comprised of two complementary mass spectrometry methods, LC-MS (a single mass spectrometry analysis following physical sample separation using liquid chromatography) and LC-MS/MS (liquid chromatography-tandem mass spectrometry, with two mass spectrometry runs to select specific ions and subsequently fragment those ions).

*LC-MS.* The column eluent was introduced directly into a Xevo-G2-S quadrupole-time-of-flight (QTOF) mass spectrometer (Waters Corp.) by electrospray. Mass spectrometry was performed on a QTOF mass spectrometer operating in either negative or positive electrospray ionization mode. Data was acquired in Centroid mode with a 50.0 to 1200.0 m/z mass range for TOF MS scanning. Data acquisition quality was performed using pooled QC samples (see **Supplemental Figures 1and 3**). Metabolite annotations from this method are understood to be of Level 1-2 confidence, through identification from core facility panel confirmed by reference standard (See Schymanski et al^25^ for metabolomics annotation confidence guidelines).

**Figure 3.**
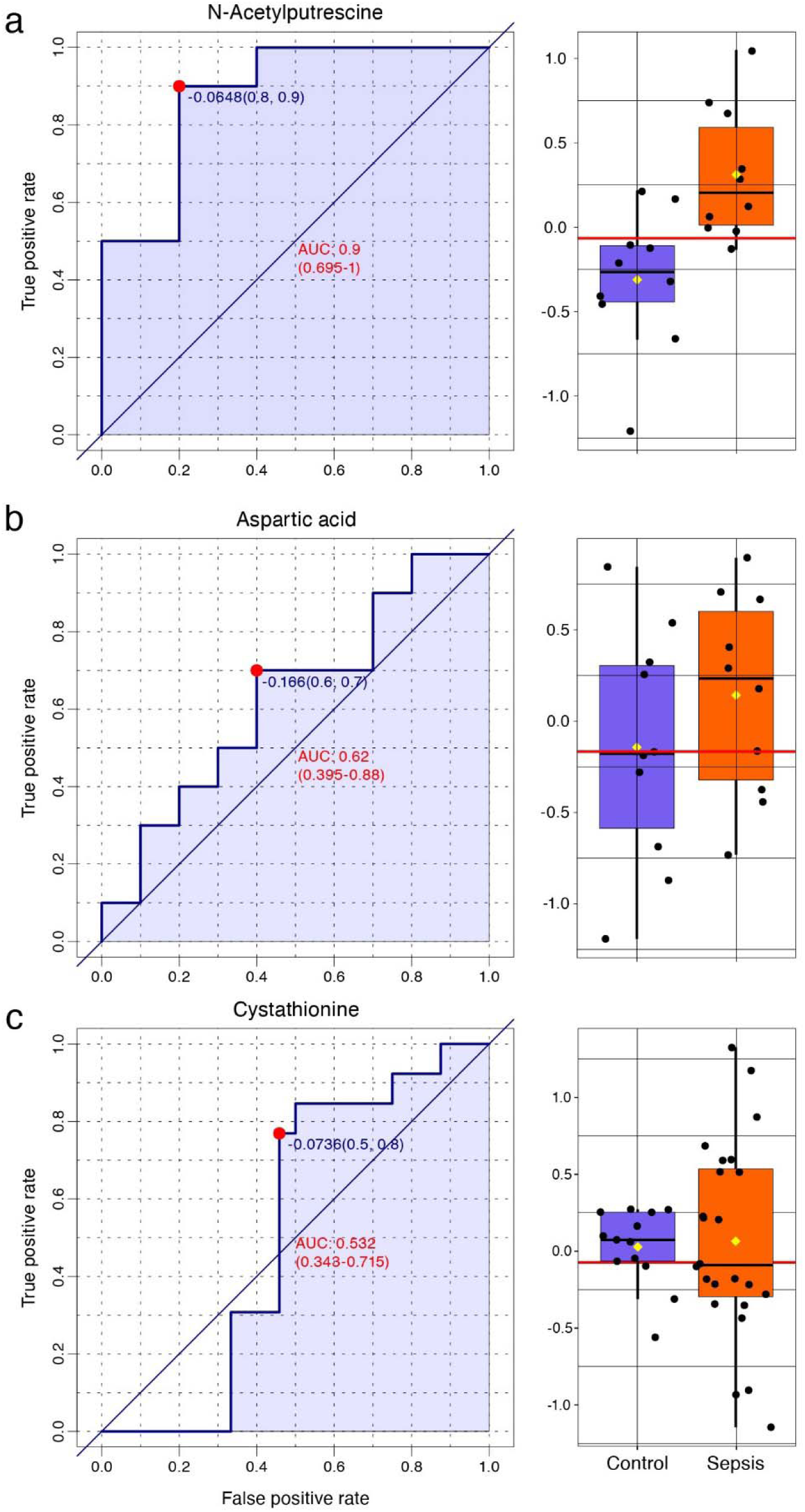
Human metabolite biomarker matches/analogs of identified mouse metabolites. This figure displays the three metabolite features in human blood plasma that were consistently altered in both cerebellar and plasma compartments in mice. Samples considering septic and non-septic (control) patients. Panels **a-c** display receiver operating characteristic (ROC) curves for each overlapping metabolite candidate, with AUC values shown to reflect classification performance. Box plots are displayed to the right of each metabolite’s ROC curve, illustrating the relative average concentrations of each metabolite by condition. The best diagnostic threshold is displayed by a red horizontal line on the box plots.

*LC-MS feature annotation.* Features that were found to differ significantly between groups following detection via LC-MS analysis were subsequently subjected to LC-MS/MS analysis to obtain spectral fragmentation data to enhance the annotation of those features. Samples were again introduced into the Xevo-G2-S QTOF (Waters Corp). Mass spectrometry was performed on a QTOF mass spectrometer operating in either negative or positive electrospray ionization, this time targeting specific m/z determined to differ significantly between groups. These features were selected in the first quadrupole (Q1), fragmented in a traveling wave collision cell for collision-induced dissociation, and detected using high-resolution time-of-flight (TOF) mass spectrometry. Fragmentation spectral data were then used to prepare QC analysis comparisons (See **Supplemental Figures 2and 4**) and to annotate the features previously determined to differ between groups. We utilized the NIST MS Search software (version 2.4) to determine the highest probability spectral match for each targeted feature. These putative annotations are probable but not confirmed by reference samples, earning a Level 3+ degree of confidence^25^.

**Figure 4.**
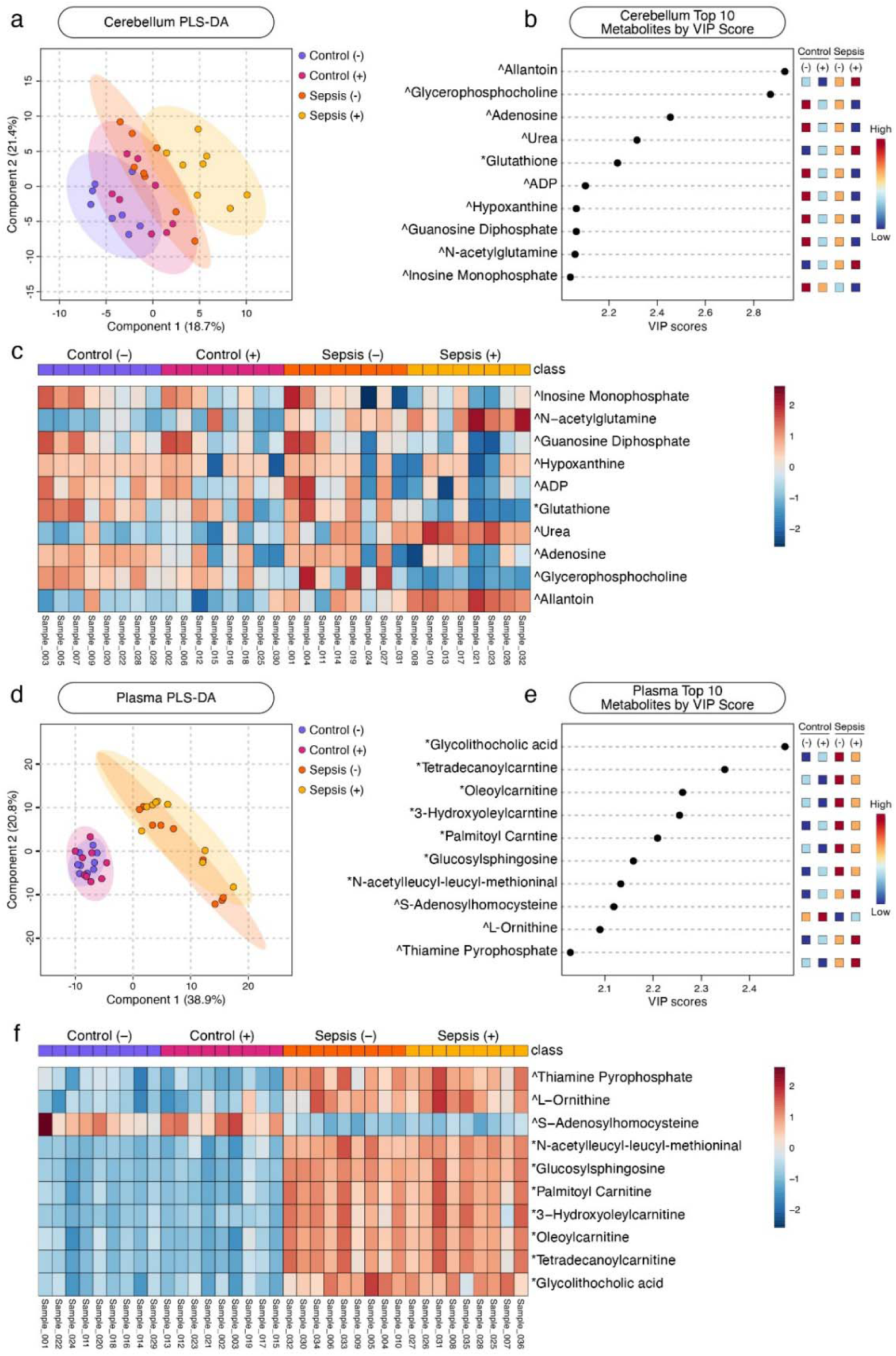
**Multivariate analysis identifies discriminatory metabolite profiles in cerebellar tissue and plasma**. PLS-DA plots of cerebellum (**a**) and plasma (**d**) samples where the horizontal and vertical axes represent Components 1 and 2, respectively. These components capture directions in the data that maximize covariance between metabolite profiles and group labels where Component 1 explains the strongest class-related pattern and Component 2 the next most discriminative, orthogonal pattern. Ellipses represent the 95% confidence interval. **b**) Variable Importance in Projection (VIP) score graphs for cerebellum (**b**) and plasma (**e**) samples display the top 10 metabolites contributing to group separation along Component 1, with VIP scores reflecting each metabolite’s contribution to the model’s power to discriminate samples by group. Heatmaps of the top 10 metabolites by VIP score from cerebellum (**c**) and plasma (**f**) PLS-DAs showing relative peak intensities across each sample and grouped by experimental condition. Caret symbols (^ denote a Level 1-2 confidence identification from core facility panel confirmed by reference standard; Asterisks (*) denote a putatively annotated feature from LC-MS/MS fragmentation matches using the NIST MS Search software, earning Level 3+ identification confidence^25^.

### Biomarker candidate selection and evaluation

Metabolites were evaluated on their capacity for discriminating animals between Sepsis (-) and Control (-) conditions based on concentration levels in both cerebellar tissue and plasma. Receiver operating characteristic (ROC) curves were generated using MetaboAnalyst 6.0^26^ and MetaboAnalystR^27^ to evaluate and visualize sensitivity-specificity trade-offs for individual metabolites. Biomarker viability was evaluated using models’ area under the ROC curve (AUC) as a measure of the metabolite’s ability to distinguish between Sepsis (-) and Control (-) groups. Metabolites with AUC values ≥ 0.8 and < 0.9 have considerable diagnostic performance and those with AUC values ≥ 0.9 have excellent performance^28^. In line with these criteria interpretations, endogenous metabolites with AUC values ≥ 0.8 were considered as biomarker candidates. Human metabolites matching those discovered in mice were subsequently evaluated using ROC curves to determine discriminatory value is determining septic patients from non-septic patients.

### Comprehensive metabolomic profiling validity

Although untargeted metabolomics offers broad profiling capabilities, its use in CNS-focused models is often constrained by annotation uncertainty and biological ambiguity. To address this, we employed a hybrid approach, integrating untargeted discovery with targeted reanalysis to ensure biological relevance, reproducibility, and quantitative reliability. For this reanalysis we repeated our core metabolic profiling analyses using only metabolites from the high confidence targeted metabolomics panel. This included partial least squares discriminant analysis (PLS-DA), pathway analyses, upstream regulator analyses, and biomarker screening, as described below.

### Multivariate modeling

For multivariate modeling, we used unsupervised principal component analysis (PCA). For clarity of presentation, we used PLS-DA, a supervised dimensionality reduction technique that incorporates known group labels to identify latent variables that best discriminate samples between experimental conditions. While this method is known to overfit data for small sample sizes, we chose this method for narrative clarity to characterize group-level differences in metabolic profiles and identify the metabolites which have the strongest discriminatory power. Analyses were completed using the MetaboAnalyst 6.0 web interface^26^ and the MetaboloAnalystR 4.0 R package^27^. Features that were putatively annotated as metabolites known to be exogeneous to mice were removed from consideration.

### Pathway Analyses and Upstream Regulators

To determine pathways that were most significantly affected by differences in the screened differential metabolites from previous analyses, we utilized Qiagen Ingenuity Pathway Analysis (IPA; Qiagen Inc., https://digitalinsights.qiagen.com/IPA)^29^, which references its internal catalog of primary literature to determine top canonical pathways that are expected to be up- or down-regulated based on uploaded concentrations of metabolites from our samples (see Kramer et al.^29^ for a full description of IPA prediction algorithms).

In considering which top canonical pathways to consider for further analysis, we determined three pathway characteristics of interest: statistical significance, pathway coverage (number of metabolites from pathway for which we had measurements), and whether directionality of pathway regulation could be determined. Accordingly, we established a metric that we called “Pathway Impact” that incorporates these three characteristics, along with weights for each characteristic, as follows:

Pathway Impact = w^l^ · Normalized Significance + w^2^ · Pathway Coverage + w^3^ · Directionality Score. “Normalized Significance” was normalized to a range of [0,1] by dividing each pathway’s −log(p-value) by the maximum observed −log(p-value) across all values, “Pathway Coverage” was reported as a ratio between [0,1] (where “zero” reflects no available measurements for metabolites from the pathway and “one” reflects all known metabolites from the pathway measured), and “Directionality Score” was a binary score based on the availability of directionality information (up- or down-regulation; 0 = no direction information, 1 = known directionality). Finally, we applied labels to impact scores, “+” or “-“, which represent predicted up- or down-regulation, respectively. For this study, weights of 0.5, 0.3, and 0.2 were used for “Normalized Significance”, “Pathway Coverage”, and “Directionality Score”, respectively.

Comparison of the Sepsis (-) v. Control (-) groups served to assess the disease state condition, Sepsis (+) v. Sepsis (-) as the treatment state condition, and the four remaining comparisons (Control (+) v. Control (-), Sepsis (-) v. Control (+), Sepsis (+) v. Control (-), Sepsis (+) v. Control (+)) served as subordinate supplemental comparisons. We also considered the top 10 identified differences between conditions in upstream regulators as determined by absolute value of their activation z-score (i.e., those regulators that were projected to be influenced the most by condition) using Qiagen IPA’s Upstream Regulator Analysis.

### Human subjects

Human participants aged 3 months-17 years were recruited from the emergency room (ER) at Children’s National Hospital in Washington, D.C. between January 2020 and May 2022 in two separate cohorts. A total of 35 septic children were enrolled to the study. Those patients either received a sepsis diagnosis from an clinician based on current diagnostic criteria, including ICD-10 codes for sepsis or were treated for sepsis with 2 intravenous fluid boluses and antibiotics (sepsis treatment per standard of care). A group of 23 non-septic children made up the control group, consisting of children with no evidence of infection as determined by clinical evaluation. Blood samples were collected from all participants within 6 hours of admission to the ER. Blood was collected in ethylenediaminetetraacetic acid (EDTA)-containing tubes and plasma was separated following centrifugation. Samples were then immediately stored at −80°C following standardized biobank protocols.

In Cohort 1, the sepsis group (n = 25) had a mean age of seven years and six months (range 11 months – 17 years and 11 months), while controls (n = 13) had a mean age of 14 years and 7 months (range eight years and nine months – 18 years). Seventeen septic participants (68%) and five controls (38%) were male. In Cohort 2, the sepsis group (n = 10) had a mean age of seven years and 10 months (range two years –13 years and seven months), compared to 13 years and seven months (range eight years –17 years) in controls (n = 10). Six septic participants (60%) and five controls (50%) were male.

This study was approved by the Institutional Review Board of Children’s National Hospital, and parental or guardian consent was obtained prior to participation in accordance with institutional policy. No patient-identifying information in the public data or this manuscript, in accordance with applicable guidelines and regulations. All data were also anonymized before analysis. All procedures followed the Declaration of Helsinki (2020/340).

## RESULTS

### Cecal slurry injection induced acute sepsis and SE with clinical recovery following human MSC-sEV treatment

A total of 61 animals (n_Control (-)_ = 16, n_Control (+)_ = 15, n_Sepsis (-)_ = 15, n_Sepsis (+)_ = 15) were evaluated using the aforementioned validated murine sepsis severity rubric^24^ (**Figure 1b**). Following a one-way ANOVA comparing average total sepsis scores at the time of treatment injection (F_(3,57)_ = 61.18, p < 0.001), Tukey’s HSD post-hoc analysis revealed no significant difference between Sepsis (+) and Sepsis (-) groups (M_Sepsis (+)_ = 3.93, M_Sepsis (-)_ = 3.00, p = 0.07). In considering average peak total sepsis scores during observations (i.e., highest sepsis score throughout the 24-hour observation period), we found a significant main effect of condition (F_(3,57)_ = 253.05, p < 0.001), characterized by a significant decrease in septic mice treated with MSC-sEVs relative to untreated mice (M_Sepsis (+)_ = 11.13, M_Sepsis (-)_ = 12.87, p = 0.03) according to post-hoc analyses. Despite a significant main effect in the average total sepsis scores between sepsis groups at the time of sacrifice as determined by one-way ANOVA (F_(3,57)_ = 125.40, p < 0.001), Tukey’s HSD post-hoc analysis found no significant difference between sepsis groups (M_Sepsis (+)_ = 10.13, M_Sepsis (-)_ = 11.47, p = 0.35). At each of the three points of measurement described above, both sepsis groups scored significantly higher relative to both control groups (p < 0.001 for each comparison), while there was no significant difference between Control (-) and Control (+) groups (p_Injection_ = 1.0, p_Peak_= 0.99, p_Sacrifice_ = 1.0).

We also considered a subset of the total sepsis scores previously reported^5^ to assess CNS-related function (i.e., level of consciousness, general activity, and response to stimuli; **Figure 1c**). In addition to a significant main effect of condition as determined by one-way ANOVA (F_(3, 57)_ = 49.26, p < 0.001), Tukey’s HSD post-hoc analysis comparing average CNS-related sepsis scores at the time of treatment injection revealed that mice in the Sepsis (+) group had a significantly higher average CNS-related sepsis score at the time of treatment injection relative to mice in the Sepsis (-) group (M_Sepsis (+)_ = 2.73, M_Sepsis (-)_ = 1.93, p = 0.03).

Following a significant one-way ANOVA considering average CNS-related peak scores (F_(3,57)_ = 283.57, p < 0.001), Tukey’s HSD post-hoc analysis determined that there was a significant decrease of CNS-related peak scores in the group treated with MSC-sEVs relative to non-treated animals (M_Sepsis (+)_ = 6.47, M_Sepsis (-)_ = 7.67, p = 0.005). While there was an overall main effect of condition on average CNS-related sepsis scores at the time of sacrifice between groups (F_(3,57)_ = 161.08, p < 0.001), post-hoc analysis revealed no significant difference between sepsis groups at the time of sacrifice (M_Sepsis (+)_ = 5.87, M_Sepsis (-)_ = 6.93, p = 0.06). At each of the three points of measurement described above, both sepsis groups had significantly higher scores relative to control groups (p < 0.001 for each comparison), while there was no significant difference between Control (+) and Control (-) groups (p_Injection_ = 1.0, p_Peak_= 0.98, p_Sacrifice_ = 1.0).

These clinical observations were consistent with our previously published findings, confirming the successful induction of acute sepsis and mitigation of both systemic and CNS-specific severity following MSC-sEV treatment under the same experimental conditions^5^. Building on our prior evidence of cerebellar vulnerability in this model, the current findings support a cerebellum-focused metabolomic analyses to investigate the underlying mechanisms of injury and MSC-sEV-mediated recovery.

### Metabolic profiling revealed distinct cerebellar and plasma signatures in sepsis and treatment

The feature detection process was applied to both mouse cerebellar tissue and plasma samples to identify and annotate as many features/metabolites as possible (see **Supplemental Table 1** for a summary). Using untargeted LC-MS, we detected 9550 features in cerebellar tissue, including 4024 in NEG ionization mode and 5526 in POS ionization mode. In plasma samples, 10,366 features were detected (5187 NEG and 5179 POS). Of these detected features, 463 in cerebellar tissue and 671 in plasma samples differed significantly in concentration between groups. Importantly, these represent raw spectral features, many of which are not directly interpretable as known metabolites.

Samples were re-injected and tandem MS was performed on features previously identified as significant, using targeted retention time and m/z windows for collision-induced dissociation (CID). Following LC-MS/MS analysis, fragmentation spectra were compared with spectral libraries using the NIST MS Search 2.4 tool which allowed for the putative annotation of 174 metabolites from mouse cerebellar tissue and 373 metabolites from mouse plasma samples. Combined with the 324 known metabolites from the metabolite panel (previously annotated), a total of 498 metabolites in cerebellar tissue and 697 in plasma samples were annotated and quantified for use in all subsequent downstream analyses.

### Cross-compartment metabolite screening identified candidate plasma biomarkers of sepsis-associated cerebellar injury

To evaluate the potential for peripheral biomarkers of sepsis-associated cerebellar injury, we next performed cross-compartment analyses of metabolite candidates from mouse cerebellar tissue and plasma. ROC analyses were used to assess diagnostic accuracy and efficiency of candidate metabolites in distinguishing septic from control animals. We identified metabolites with considerable (AUC values ≥ 0.8) and excellent (AUC values ≥ 0.9) diagnostic criteria^28^ between septic and control animals in cerebellar tissue and plasma. Among the cerebellar metabolite candidates, we aimed to identify overlapping plasma metabolites that could serve as peripheral indicators of sepsis-associated cerebellar injury.

We found 24 metabolites with AUC values ≥ 0.8 that were considered as viable biomarker candidates from cerebellar tissue (**Supplemental Figure 14**). Of these, 16 were up-regulated in septic animals relative to controls: ^N-acetylglutamine (AUC: 0.977), *N-oleoylglycine (AUC: 0.914), ^Betaine (AUC: 0.875), ^Cystathionine (AUC: 0.875), *1-(9Z-octadecenoyl)-sn-glycero-3-phosphocholine (AUC: 0.859), *17-Oxo-4(Z),7(Z),10(Z),13(Z),15(E),19(Z)-docosahexaenoic acid (AUC: 0.844), ^L-acetylcarnitine (AUC: 0.844), *Ptdlns-(3,4,5)-P3 (1-stearoyl, 2-arachidonoyl) (AUC: 0.844), *Substance P (3-11) (AUC: 0.844), *[Ser2]-Neuromedin C (AUC: 0.836), ^1-Methyl-L-Histidine (AUC: 0.828), *Arachidonamide (AUC: 0.828), *ADP-D-Ribose (AUC: 0.828), ^Urea (AUC: 0.828), ^Putrescine (AUC: 0.812), and *N-acetyl-L-aspartic acid (AUC: 0.812). The remaining eight were decreased in septic animals relative to controls: *Amyloid beta-protein (25-35) amide (AUC: 0.953), ^Dopamine (AUC: 0.906), ^Betaine aldehyde (AUC: 0.883), *1,2-dipalmitoyl-sn-glycero-3-phosphoethanolamine (AUC: 0.875), ^Choline (AUC: 0.875), ^Dimethylglycine (AUC: 0.875), *1,2-dilinoleoylglycerol (AUC: 0.844), and Atrial natriuretic peptide (ANP, 1-28; AUC: 0.828).

In mouse plasma, we found a total of 396 metabolites with AUC values ≥ 0.80, qualifying for biomarker consideration. Of those, 187 metabolites were decreased in septic animals relative to controls and 210 were increased in septic samples. Of the 24 biomarker candidates discovered in cerebellum, we found nine overlapping metabolites with sufficient diagnostic power (i.e., AUC ≥ 0.8) to identify septic vs. control animals present in plasma: ^1-Methyl-L-Histidine (AUC: 1), ^Choline (AUC: 1), ^Putrescine (AUC: 1), ^Dimethylglycine (AUC: 1), ^N-Acetylglutamine (AUC: 1), ^Urea (AUC: 1), ^Cystathionine (AUC: 0.963), *N-Acetyl-L-Aspartic acid (AUC: 0.877), *1-(9Z-octadecenoyl)-sn-glycero-3-phosphocholine (AUC: 0.815; **Figure 2a-i**). While all nine metabolites were retained and displayed in **Figure 2** for meeting the specified AUC criteria, six of the nine overlapping metabolites showed consistent directionality in concentrations differences between groups (e.g., Sepsis (-) showed an increase in 1-Methyl-L-Histidine concentration relative to Control (-) in both cerebellum and plasma). The remaining three metabolites, ^Choline, ^Dimethylglycine, and *1-(9Z-octadecenoyl)-sn-glycero-3-phosphocholine, had concentration levels were inconsistent between cerebellum and plasma and were not considered biologically congruent between compartments.

Of the six overlapping biomarker candidates consistent between both cerebellum and blood plasma of mice (^1-Methyl-L-Histidine, ^Putrescine, ^N-Acetylglutamine, ^Urea, ^Cystathionine, and *N-Acetyl-L-Aspartic acid), we found either a direct match, in the cases of 1-methyl-L-histidine and cystathionine, or metabolite analog (N-acetylputrescine, glutamine, and aspartic acid) for each in human blood plasma of septic patients with the exception of ^Urea, for which we found intermediate markers known from the Urea cycle (citrulline, ornithine, and arginine) to evaluate. Three of the matches/analogs found in human plasma were consistent directionally consistent in concentration between septic and non-septic patients relative to findings in mice. Specifically, N-acetylputrescine (AUC: 0.9), Aspartic acid (AUC: 0.62), and Cystathionine (AUC: 0.53) were all found in increased concentration in septic patients relative to non-septic patients (**Figure 3**). Metabolomics results for cystathionine were obtained from Cohort 1, while results for N-acetlyputrescine and aspartic acid were obtained from Cohort 2.

Through these analyses we identified a subset of cerebellar metabolites that not only distinguished septic animals from controls with high diagnostic accuracy, but were also detectable in the plasma of septic patients with comparable diagnostic value. These findings suggest that such metabolites could serve as peripheral biomarkers of cerebellar injury in sepsis. Moreover, overlapping metabolites with consistent directional changes across compartments further support their potential as clinically accessible indicators of CNS-specific metabolic disruption.

### Cerebellar metabolomic signatures reflected both disease and treatment effects, while plasma profiles were driven by disease status alone

Maps of the first two components identified by PLS-DA modeling of cerebellar tissue results are shown in **Figure 4a**, where within-group samples are generally shown to cluster together (shaded ovals represent a 95% confidence interval). Component 1 explained 18.7% of the variance in the metabolite data matrix (X), which consists of the relative abundances of metabolites across all samples and captured the strongest direction of group separation. Component 2 explained 21.4% of the X variance and represented a secondary, orthogonal pattern that also contributed to group separation. Although Component 2 captured more of the total variance in metabolite concentrations across all samples, Component 1 was assigned as such because it more closely aligned with group labels, consistent with the goal of PLS-DA in maximizing the covariance between predictors (X) and group membership (Y). An analogous PCA plot showing the unsupervised separation between groups can be found in **Supplemental Figure 5a.**

Using 5-fold cross validation, we found that the model performed best with four group comparisons, yielding an average classification accuracy of 0.608, R^2^ of 0.790, and Q^2^ of 0.777, indicating strong performance with high explained variance and predictive capability. Notably, each of the four experimental groups forms a partially distinct cluster, suggesting the cerebellar metabolic profile is sensitive to both disease and treatment effects. We also obtained variable impact importance in projection (VIP) scores from Component 1, which quantify each metabolite’s weighted contribution to the model’s discriminative power (the top 10 metabolites by VIP score and their relative concentrations in each group are displayed in **Figure 4b**). Relative concentrations of these top 10 metabolites for individual samples are displayed in the heatmap found in **Figure 4c**.

In contrast to cerebellum, a PLS-DA model of plasma samples revealed clear separation between control and sepsis groups with limited separation based on treatment (i.e., Control (-) and Control (+) groups clustered together, as did Sepsis (-) and Sepsis (+) groups; see **Figure 4d**). This lack of treatment-associated separation in plasma, in stark contrast to the distinct treatment effects observed in cerebellar tissue, suggests that MSC-sEV treatment impact was localized rather than systemic. Component 1 explained 38.9% of the variance in the metabolite data matrix while Component 2 explained 20.8%. The PLS-DA was optimized with two group comparisons, yielding an average classification accuracy of 0.412, R^2^ of 0.613, and Q^2^ of 0.538, indicating moderate separation with acceptable predictive capacity. Top 10 metabolites by VIP score and their relative concentrations are shown in **Figure 4e**. Relative concentrations of these top 10 metabolites for individual samples are displayed in the heatmap found in **Figure 4f**. See **Supplemental Figure 5b** for a PCA plot of these groups.

Cumulatively, these results illustrated that cerebellar metabolic profiles were influenced by both disease state and treatment conditions, while plasma profiles were primarily influenced by disease state condition alone. Having established global metabolomic patterns across cerebellar tissue and plasma, we next focused exclusively on cerebellar tissue to isolate sepsis- and treatment-induced metabolic changes specifically in this brain region.

### Cerebellar metabolomic profiling revealed metabolic dysfunction, suppression of oxidative energy metabolism, and redox imbalance in SE

To characterize metabolic alterations associated with SE, we applied an integrated metabolic analysis combining PLS-DA modeling, pathway enrichment, and upstream regulator prediction using annotated and quantified metabolites from cerebellar tissue. The following results identify key metabolites, canonical pathways, and upstream regulators driving changes across septic and control conditions.

Isolating disease state, we identified a PLS-DA model with strong separation between groups (**Figure 5a**) where Component 1 explains 13.3% of the variance in the metabolite data matrix and Component 2 explains 28.6%. This model was optimized for two groups, resulting in an average classification accuracy of 0.817, R^2^ of 0.886, and Q^2^ of 0.474. We then identified the top 10 metabolites driving differences between groups using metabolite VIP scores (ranging from 2.15 – 4.22 for these top 10 features; see **Figure 5b**). Identified metabolites include *N-oleoglycine, ^Spermine, ^Spermidine, ^Dopamine, ^Hypoxanthine,^Putrescine, ^1-methyl-L-histidine, ^3-hydroxyl-3-methylglutaryl-COA, ^Glucosamine, and ^Guanosine. All metabolites with the exceptions of ^Dopamine and ^Hypoxanthine were increased in the septic group relative to controls (see **Figure 5c-l**; ^ Denotes a Level 1-2 confidence identification from core facility panel confirmed by reference standard; * Denotes a putatively annotated feature from LC-MS/MS fragmentation matches using the NIST MS Search software, earning Level 3+ identification confidence^25^). A PCA plot of this comparison can be found in **Supplemental Figure 7a.**

**Figure 5.**
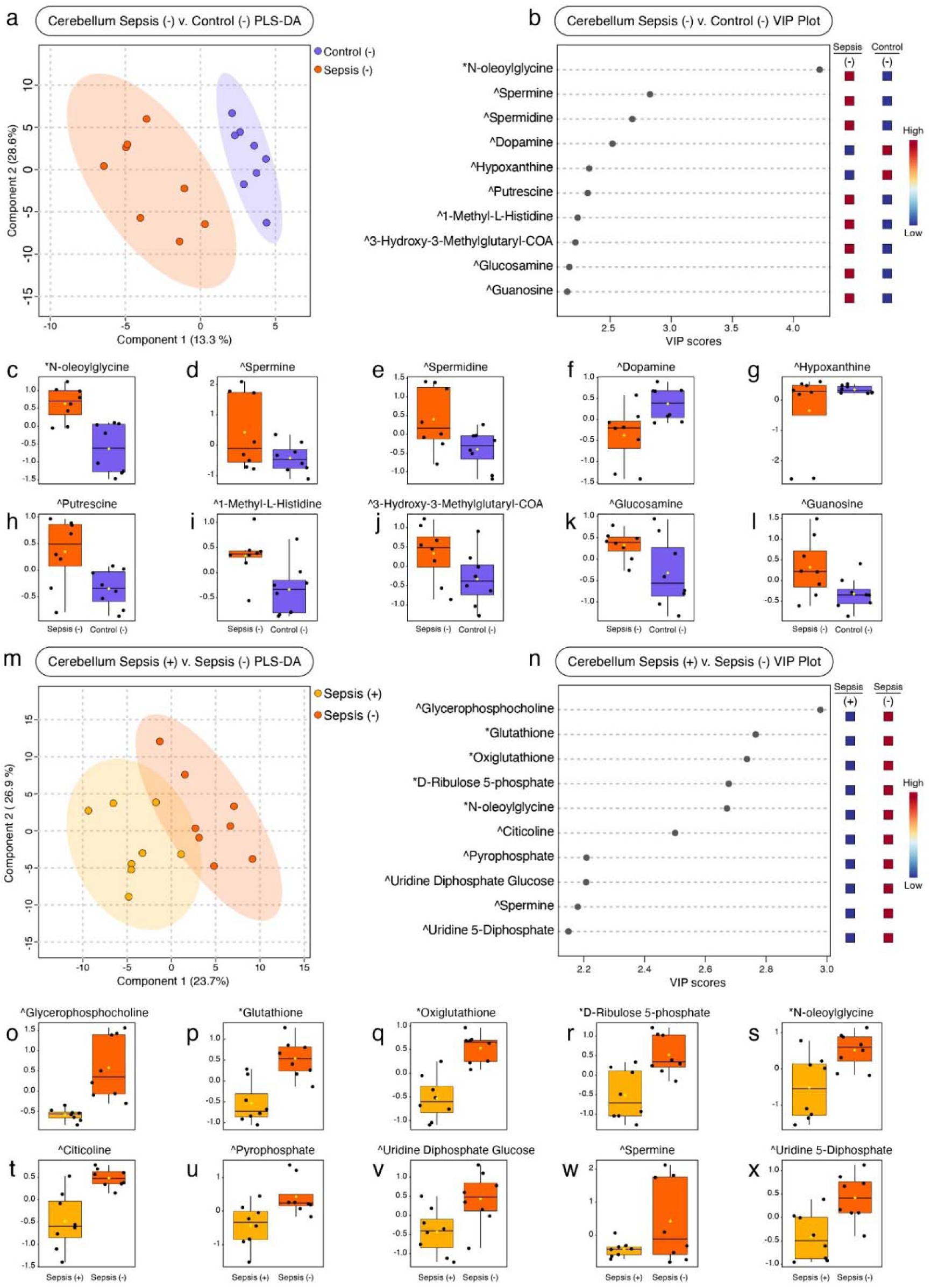
**Variable isolating group comparisons in cerebellar metabolite profiles reveal effects of disease state and treatment**. This figure presents PLS-DA-derived discriminant features from cerebellar samples, restricted to two pairwise group comparisons. Panels **a-l** illustrate the comparison between untreated septic animals and healthy control, Sepsis (-) v. Control (-), isolating the metabolic profiles associated with disease state. Panels **m-x** display data from the Sepsis (+) v. Sepsis (-) comparison, capturing effects related to the effect of MSC-sEV treatment under septic conditions. (**a, m**) PLS-DA model illustration showing sample clustering along the first two components with 95% confidence ellipses for each group. (**b, n**) VIP score plots showing the top 10 metabolites contributing to Component 1 and driving separation by experimental condition. (**c-l, o-x**) Box plots for each of the top 10 metabolites by VIP score in their respective comparisons, depicting average relative peak intensity by group. Caret symbols(^) denote a Level 1-2 confidence identification from core facility panel confirmed by reference standard; Asterisks (*) denote a putatively annotated feature from LC-MS/MS fragmentation matches using the NIST MS Search software, earning Level 3+ identification confidence^25^.

Pathway analyses resulted in a total of 904 canonical pathways predicted to be altered between septic and non-septic conditions. **Figures 4a** displays all pathways predicted to be altered between disease states. Each bubble represents an altered pathway and is grouped based on biological function (y-axis). We further explored the nature of the top 20 pathways (illustrated in **Figure 6b**) as determined by Pathway Impact score (see **Supplemental Figures 7-10** and **Supplemental Table 2** for data from additional comparisons).

**Figure 6.**
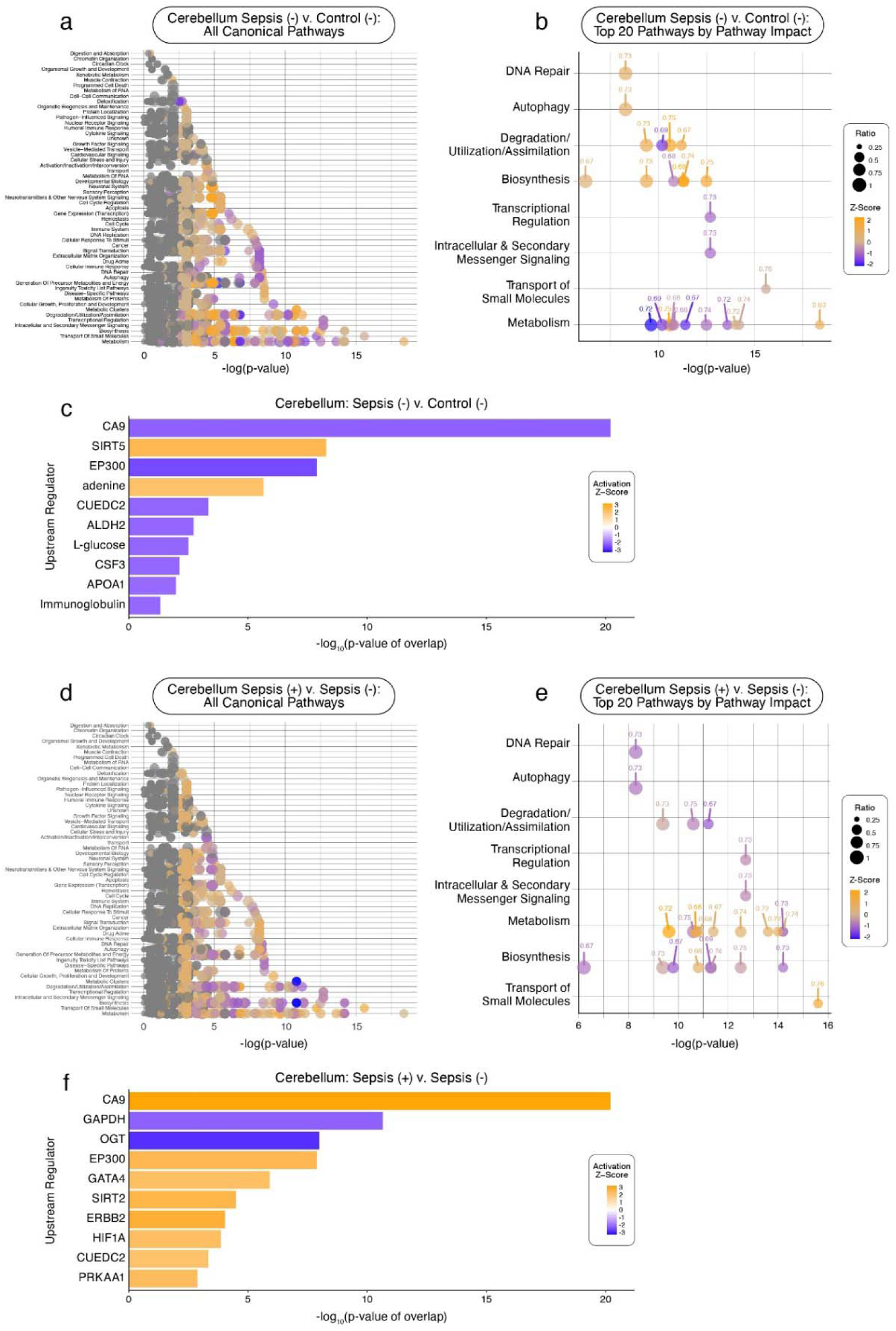
**Canonical pathway activity and upstream regulator predictions in cerebellar tissue across disease and treatment comparisons**. Panels **a-c** focus on the disease state comparison, Sepsis (-) v. Control (-), while panels **d-f** focus on the treatment comparison, Sepsis (+) v. Sepsis (-). (**a,d**) Canonical pathway activity showing all pathways predicted to be affected as determined by cerebellar metabolite intensities. Pathways are grouped by their biological function (y-axis) and plotted against their alteration significance (-log_10_[p-value]; x-axis). Bubble size reflects the proportion of the measured metabolites relative to total known pathway metabolite profile, while bubble color represents the predicted activation direction and intensity (orange = predicted up-regulation, blue = predicted down-regulation). (**b,e**) Top 20 canonical pathways as ranked by Pathway Impact score. (**c,f**) Top 10 predicted upstream regulators influenced by each condition based on metabolite representation. Each regulator (y-axis) is displayed as a bar with a length representing its −log_10_(p-value of overlap; x-axis) and is colored according to its predicted activation direction and intensity.

We identified the top 20 affected pathways and grouped them into eight categories based on their overall biological functions, listed in order of descending impact: metabolism, transport of small molecules, intracellular and secondary messenger signaling, transcription regulation, biosynthesis, degradation/utilization/assimilation, autophagy, and DNA repair (**Figure 6b**, see **Table 1** for a complete list of top 20 pathways, their biological function category, and Pathway Impact score). The five most significantly impacted pathways included nucleotide catabolism, transport of bile salts and organic acids, metal ions and amine compounds, superpathway of citrulline metabolism, urea cycle, and arginine biosynthesis IV.

**Table 1.** Top 20 canonical pathways altered in cerebellar tissue by disease and treatment conditions. Listed in this table are the top 20 canonical pathways altered from between disease state, Sepsis (-) v. Control (-), and treatment, Sepsis (+) v. Sepsis (-), comparisons as determined by Pathway Impact score. This table identifies the individual pathway names, biological function category to which each pathway belongs, and the Pathway Impact score for each pathway. Direction of regulation is denoted by sign on impact score: positive (+) for predicted up-regulation, negative (-) for predicted down-regulation.

| Sepsis (-) v. Control (-) |  |  |  | Sepsis (+) v. Sepsis (-) |  |  |
| --- | --- | --- | --- | --- | --- | --- |
| Rank | Pathway Name | Biological Function Category | Impact Score | Pathway Name | Biological Function Category | Impact Score |
| 1 | Nucleotide catabolism | Metabolism | +0.83 | Transport of bile salts & organic acids, metal ions & amine compounds | Transport of Small Molecules | +0.76 |
| 2 | Transport of bile salts & organic acids, metal ions & amine compounds | Transport of Small Molecules | -0.76 | Superpathway of Citrulline Metabolism | Biosynthesis | -0.75 |
| 3 | Superpathway of Citrulline Metabolism | Biosynthesis | +0.75 | Urea Cycle | Metabolism, Degradation/Utilization/Assimilation | -0.75 |
| 4 | Urea Cycle | Metabolism, Degradation/Utilization/Assimilation | +0.75 | Arginine Biosynthesis IV | Biosynthesis | -0.74 |
| 5 | Arginine Biosynthesis IV | Metabolism | +0.74 | Citric acid cycle (TCA cycle) | Metabolism | +0.74 |
| 6 | Citric acid cycle (TCA cycle) | Metabolism | -0.74 | Interconversion of nucleotide di- and triphosphates | Metabolism | +0.74 |
| 7 | Interconversion of nucleotide di- and triphosphates | Metabolism | -0.74 | Nucleotide salvage | Metabolism | -0.73 |
| 8 | Citrulline-Nitric Oxide Cycle | Biosynthesis, Degradation/Utilization/Assimilation | +0.73 | Citrulline-Nitric Oxide Cycle | Biosynthesis, Degradation/Utilization/Assimilation | -0.73 |
| 9 | Sirtuin Signaling Pathway | Intracellular & Secondary Messenger Signaling, Transcriptional Regulation | -0.73 | Sirtuin Signaling Pathway | Intracellular & Secondary Messenger Signaling, Transcriptional Regulation | -0.73 |
| 10 | Mismatch Repair | Autophagy, DNA Repair | +0.73 | Mismatch Repair | Autophagy, DNA Repair | -0.73 |
| 11 | Sulfur amino acid metabolism | Metabolism | -0.72 | Sulfur amino acid metabolism | Metabolism | +0.72 |
| 12 | Metabolism of water-soluble vitamins and cofactors | Metabolism | +0.72 | Metabolism of water-soluble vitamins and cofactors | Metabolism | +0.72 |
| 13 | Choline catabolism | Metabolism | -0.71 | Choline catabolism | Metabolism | +0.72 |
| 14 | Purine Nucleotides De Novo Biosynthesis II | Biosynthesis | +0.69 | Purine Nucleotides De Novo Biosynthesis II | Biosynthesis | -0.69 |
| 15 | Glutamate and glutamine metabolism | Metabolism | -0.68 | Glutamate and glutamine metabolism | Metabolism | +0.68 |
| 16 | Nucleotide biosynthesis | Metabolism | -0.68 | Nucleotide biosynthesis | Metabolism | +0.68 |
| 17 | Pyruvate metabolism | Metabolism | -0.67 | Pyruvate metabolism | Metabolism | +0.67 |
| 18 | Superpathway of Methionine Degradation | Degradation/Utilization/Assimilation | +0.67 | Superpathway of Methionine Degradation | Degradation/Utilization/Assimilation | -0.67 |
| 19 | Flavin Biosynthesis IV (Mammalian) | Biosynthesis | +0.67 | Flavin Biosynthesis IV (Mammalian) | Biosynthesis | -0.67 |
| 20 | Cytosolic sensors of pathogen-associated DNA | Immune System | +0.67 | Glucogenesis I | Biosynthesis | -0.67 |

We further found that the upstream regulators CA9, SIRT5, EP300, adenine, CUEDC2, ALDH2, L-glucose, CSF3, APOA1, and immunoglobulin were the most influenced regulators. All regulators were down-regulated in septic animal relative to controls, with the exceptions of SIRT5 and adenine which were up-regulated (see **Figure 6c**).

Changes at the metabolites, pathways, and regulator networks following sepsis induction revealed widespread metabolic dysfunction in the cerebellum. Findings suggested disruptions in nucleotide turnover, oxidative energy metabolism, and redox balance. The culmination of these disruptions established a baseline against which the metabolic mechanisms of MSC-sEV treatment cerebellar injury mitigation were assessed.

### Human MSC-sEV treatment partially reversed sepsis-induced cerebellar metabolic disruptions

We next applied the same integrated metabolic profiling procedure using PLS-DA, pathway analysis, and upstream regulator prediction to cerebellar tissue of human MSC-sEV-treated septic mice compared to untreated. The following results identify treatment-associated changes in metabolite levels, canonical pathways, and predicted upstream regulators between MSC-sEV-treated and untreated septic animals.

We identified an effective prediction model constructed by way of PLS-DA, where Component 1 explains 23.7% of the variance in the metabolite data matrix and Component 2 explains 26.9% (**Figure 5m**). This model was optimized with 2 components, resulting in an accuracy measure of 0.75, an R^2^ value of 0.779, and a Q^2^ value of 0.399. The top 10 metabolites driving differences between groups were then identified (VIP scores ranging from 2.15 – 2.98; see **Figure 5n**) as ^Glycerophosphocholine, *Glutathione, *Oxiglutathione, *D-Ribulose 5-phosphate, *N-oleoglycine, ^Citicoline, ^Pyrophosphate, ^Uridine Diphosphate Glucose, ^Spermine, and ^Uridine 5-diphosphate. Each of these metabolites were lower in concentration in MSC-sEV treated animals relative to untreated animals (see **Figure 5o-x**). A PCA plot from this comparison can be found in **Supplemental Figure 7b**.

Pathway analysis identified a total of 904 impacted pathways (**Figure 6d**). Notably, the top 20 pathways fell into the same eight biological function categories previously found to be influenced by sepsis: transport of small molecules, biosynthesis, metabolism, intracellular and secondary messenger signaling, transcriptional regulation, degradation/utilization/assimilation, autophagy, and DNA repair (**Figure 6e**, **Table 1**). The top five most influenced pathways as determined by Pathway Impact score were determined to be transport of bile salts and organic acids, metal ions and amine compounds, superpathway of citrulline metabolism, urea cycle, arginine biosynthesis IV, and citric acid cycle (tricarboxylic acid [TCA] cycle).

Analysis of upstream regulators in cerebellar tissue identified the top 10 most influential regulators, which included CA9, GAPDH, OGT, EP300, GATA4, SIRT2, ERBB2, HIF-1α, CUEDC2, and PRKAA1. Each of the listed regulators were predicted to be up-regulated in MSC-sEV treated septic mice relative to non-treated, with the exceptions of GAPDH and OGT, which were predicted to be down-regulated (see **Figure 6f**).

In summary, alterations in the metabolic profiles of human MSC-sEV treated animals revealed partially reversed sepsis-induced metabolic dysfunction in the cerebellum. Data from metabolite, pathway, and upstream regulator analyses indicated reductions in oxidative stress markers, normalization of amino acid and nucleotide metabolism, and activation of antioxidant and repair-associated regulators. These findings support a treatment model focused on restoring redox homeostasis and cellular energetics. These results also provided a foundation for evaluating the impact that specific MSC-sEV cargo may exert in driving the recovery observed in treated animals. Figures and tables generated from plasma data which are analogous to those described above from cerebellar tissue can be found in **Supplemental Figures 11 – 13** and **Supplemental Tables 3** and **4**.

### Confirmatory reanalysis of metabolic profiling using targeted metabolomics alone

Beginning with PLS-DA, we found consistent group separation patterns between the four experimental groups as determined by the models generated from both metabolite datasets (**Figures 2a, 2d**; **Supplemental Figures 15a, 15d;** PCA plots reported in **Supplemental Figure 16**). In cerebellum, eight of the top 10 metabolites by VIP score were consistent between both sets (**Figure 4b**; **Supplemental Figure 15b**), while three of the 10 were consistent in plasma (**Figure 4e**; **Supplemental Figure 15e**). Considering disease state- and treatment-specific comparisons in cerebellum, we again found consistent group separation in both cases (**Figures 3a, 3m**; **Supplemental Figure 17a, 17m;** PCA plots reported in **Supplemental Figure 18**). Nine of the top 10 metabolites were consistent in the disease state comparison (**Figure 5b**; **Supplemental Figure 17b**), while six out of 10 were consistent in the treatment comparison (**Figure 5n**; **Supplemental Figure 17n**). See **Supplemental Figures 19**, **20** for comparable results in plasma.

Cerebellar pathways analyses between comprehensive and targeted-only metabolomic datasets resulted in consistent pathway identification and alteration directionality in 15 out of the top 20 pathways for both disease state- and treatment-specific comparisons (**Table 1**; **Supplemental Table 5**). Cerebellar upstream regulators showed strong consistency, with seven of the top 10 overlapping in the disease state comparison (**Figure 6c**; **Supplemental Figure 41c**) and eight out 10 in the treatment comparison (**Figure 6f**; **Supplemental Figure 41f**). See **Supplemental Figure 42** for comparable results in plasma.

Finally, reanalysis of cerebellar biomarker candidates using only targeted metabolomics data identified 22 metabolites satisfying our threshold criteria with AUC scores ≥ 0.8 (eight of which were consistent with the comprehensive metabolomics dataset; **Supplemental Figure 43**). Among metabolites overlapping with plasma, we identified 13 biomarker candidates, seven of which were consistent with findings from the comprehensive metabolomics dataset (**Supplemental Figure 44**).

Overall, the reanalysis showed strong agreement between the comprehensive and targeted-only datasets across multivariate modeling, pathway analysis, upstream regulator predictions, and biomarker identification.

This consistency, especially in cerebellar tissue, reinforces the robustness of our core findings and supports the reliability of the identified metabolic signatures. At the same time, differences between datasets helped clarify the unique contributions of each approach to our broader interpretation of the influences of disease state and treatment.

## DISCUSSION

Despite increasing recognition of the CNS’s vulnerability in sepsis, the metabolic underpinnings of sepsis-associated brain injury remain poorly defined. While prior studies have highlighted inflammation and blood– brain barrier (BBB) dysfunction as contributors to SE, there is a significant knowledge gap regarding the specific metabolic alterations occurring within affected brain regions, particularly the cerebellum. Further, clinical opportunities for early intervention are limited by a lack of tissue-specific biomarkers of SE. Moreover, few studies have explored how systemic metabolic changes are reflected in those CNS regions or assessed how therapeutic interventions may modify these metabolic profiles. MSC-sEVs have emerged as a promising therapeutic platform due to their immunomodulatory and tissue-protective properties, yet the mechanisms by which they exert neuroprotective effects, especially at the metabolic level, remain largely undefined. To address these gaps, our study investigates (1) novel metabolomic biomarkers of sepsis-associated cerebellar injury that are detectable in human plasma, (2) the cerebellar tissue and plasma metabolic profiles in acute murine polymicrobial sepsis, and (3) cerebellar tissue metabolomic alterations observed following human MSC-derived sEV treatment.

We identified six metabolites (1-Methyl-L-Histidine, Putrescine, N-Acetylglutamine, Urea, Cystathionine, and N-Acetyl-L-Aspartic acid) that were elevated in both cerebellar tissue and plasma of septic animals compared to controls. Coupled with high diagnostic performance demonstrated by results from ROC curve analyses across both components, the mechanistic relevance to sepsis pathophysiology makes these metabolites strong candidates for consideration in the search for diagnostic biomarkers of sepsis-associated cerebellar damage. Of these six metabolites, we found either direct matches, analogs, or pathway-associated intermediates for each of these metabolites in metabolomics data from human blood plasma for further evaluation of biomarker validity.

Three of these six metabolites found in the human data (N-acetylputrescine, Aspartic acid, and Cystathionine) were found in higher concentration in septic patients relative to non-septic controls, consistent with the findings of both cerebellar and plasma from mice. Our first finding, putrescine, is a polyamine derived from the amino acid ornithine that has been associated with increased BBB permeability and neuronal integrity during times of neuroimmune stress^64^. Its derivative, N-acetylputrescine, has previously been reported as a potential biomarker to distinguish patients with systemic inflammatory response syndrome (SIRS) or sepsis from heathy controls^65^, which has been validated in our findings. In the present study, we found that n-acetylputrescine was elevated in septic patients and showed excellent diagnostic performance between septic and control human patients with an AUC = 0.9, making it the strongest biomarker candidate identified in our data. Our second metabolite finding, N-acetyl-L-aspartic acid, is a predominant neuronal metabolite involved in myelin lipid production and mitochondrial energy production. While there is no prior direct evidence of its effects in sepsis, its derivative N-acetyl-L-aspartyl-L-glutamate (NAAG) has been shown to increase BBB permeability, a common occurrence under septic conditions^68^. In our results, we discovered that aspartic acid was elevated in septic patients compared to controls and showed moderate discriminative performance with an AUC = 0.62. Our final biomarker candidate, cystathionine, is an important intermediate in the transsulfuration pathway, which links methionine metabolism to cysteine and glutathione synthesis and makes it an important metabolite in the maintenance of redox homeostasis. Increased concentration of cystathionine in septic animals may demonstrate an increased demand for glutathione to counteract oxidative stress. In concert, cystathionine was elevated in septic patients relative to controls, though its diagnostic performance was modest in our human plasma cohort with an AUC = 0.53. Although further validation is required, our novel cross-compartment and cross-species approach has yielded a list of strong candidates of clinically-relevant, mechanistically supported biomarker candidates and a framework for discovery that will accelerate early diagnostic efforts in SE, especially n-acetylputrescine and aspartic acid.

The three remaining mouse metabolite biomarkers were also represented in our human data in either analog or pathway intermediate form. 1-Methyl-L-Histidine is a methylated derivative of amino acid histidine, derivatives of which have been associated with protein degradation under septic conditions and identified as potential sepsis biomarkers from serum and urine analyses in critically-ill septic patients^63^. While increased in mouse profiles, findings from human data included an increased concentration of 3-methylhistidine, but is complicated by concurrent findings of decreased concentrations of 1-methylhistidine and histidine itself. These results leave the role of 1-methylhistidine as inconclusive, but does strengthen the assertion that histidine and its metabolism is tied to sepsis pathology and warrants further investigation. N-acetylglutamine is an acetylated form of the amino acid L-glutamine, a precursor of excitatory neurotransmitter glutamate. Impairments in the urea cycle during sepsis may lead to accumulations of upstream intermediates such as N-acetylglutamine, which has previously been identified as a biomarker for sepsis in a pig model^66^. In humans, we found glutamine in lower concentrations in septic patients relative to non-septic controls. Urea, a byproduct of protein metabolism, is a critical component of blood urea nitrogen SIRS levels which are commonly higher in septic patients and indicative of renal impairment^67^. Although we did not have concentrations of urea in our human data to directly compare, we did find that urea cycle intermediates such as citrulline, ornithine, and arginine were decreased in concentration in septic patients relative to controls, suggesting an increase in urea and aligning with the cross-compartment increase in urea observed in mice. Collectively, these results confirm that these remaining metabolites are consistently present in sepsis pathophysiology between compartments and species, warranting further investigation to understand individual metabolite roles and potential as diagnostic biomarkers.

Comprehensive profiling cerebellar metabolic signatures in septic and treated animals revealed extensive changes across a wide array of metabolites, offering insight into underlying cellular processes. These findings reveal significant disruptions in metabolic pathways within cerebellar tissue of septic mice compared to non-septic controls, indicating substantial neuro-metabolic involvement during sepsis. Importantly, treatment with MSC-sEVs notably reshaped these metabolic alterations, pointing to specific mechanistic pathways of action described below (see **Figure 7**). We propose potential mechanisms of action of human MSC-sEVs through their specific miRNA and protein cargo previously identified by our group (such as members of Let-7, miR-21, and miR-30 families^5^) and how these bioactive molecules may regulate metabolic and inflammatory pathways to mitigate sepsis-associated deficits as suggested by our metabolomic findings. Additionally, our comparative analysis between cerebellum and plasma provided insights into localized versus systemic metabolite profiles during sepsis. This comparative approach enabled the identification of cerebellum-specific metabolites with strong diagnostic potential in plasma, laying the groundwork for early, non-invasive detection of sepsis-associated brain injury.

**Figure 7.**
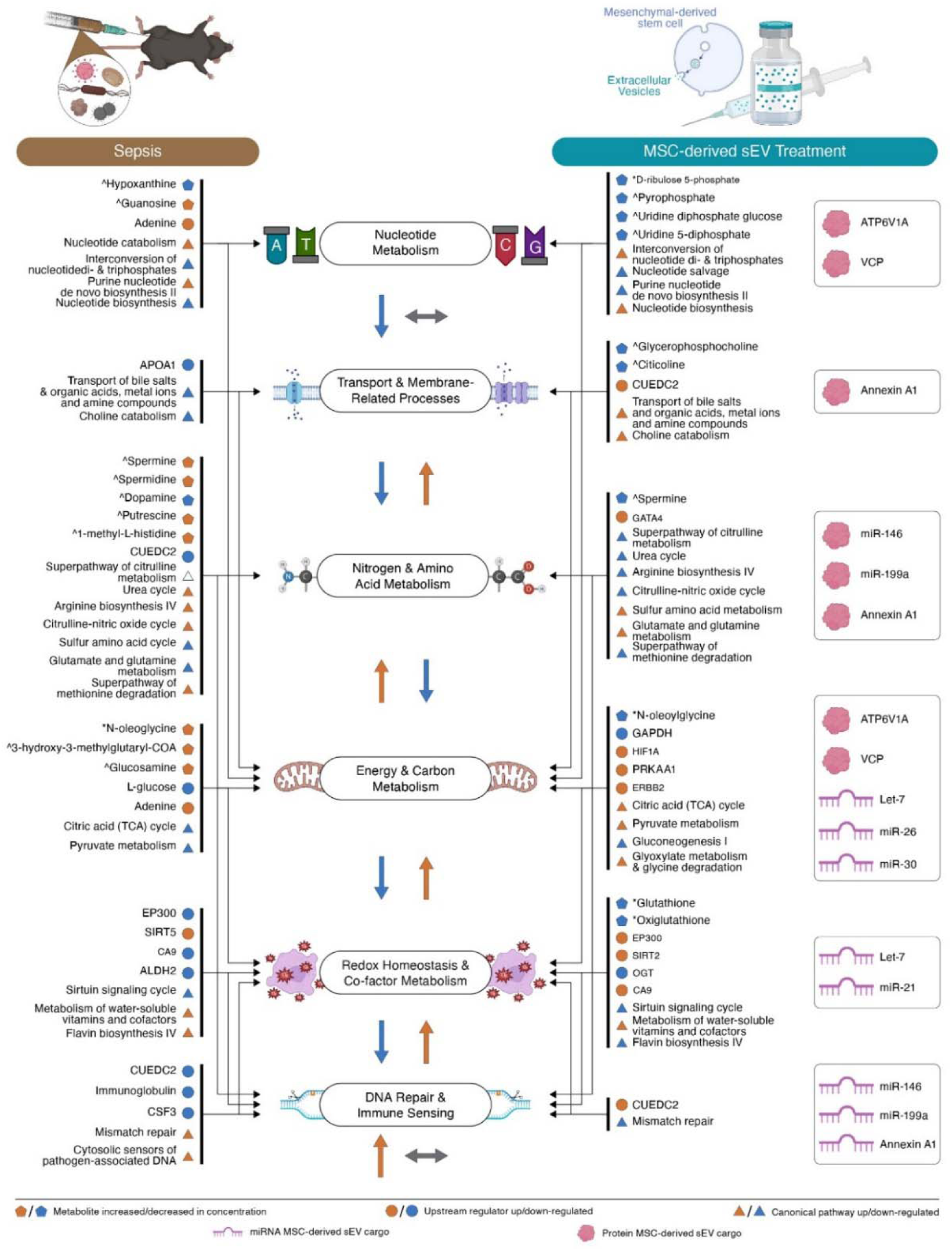
Integrated summary of comprehensive metabolic profiles (including metabolite alterations, predicted canonical pathway and upstream regulator activity) across disease and treatment conditions, alongside potential mechanistic influences of MSC-sEV cargo. This figure summarizes the affected metabolites, regulatory networks, and pathway-level alterations, illustrated as influences to inferred net biological outcomes. Arrows demonstrate which groups of metabolic profile components are likely to influence which biological outcomes. The **left column** presents changes observed or predicted in the disease state comparison, Sepsis (-) v. Control (-). The **right column** depicts changes associated with the treatment comparison, Sepsis (+) v. Sepsis (-), in addition to the potential MSC-sEV cargo influences the observed changes. Component shape denotes type (pentagon = metabolite, circle = upstream regulator, and triangle = canonical pathway) and color indicates directionality (orange = increase concentration or predicted up-regulation, blue = decreased concentration or predicted down-regulation). Large arrows under each biological outcome indicates the predicted net effect on that function. All metabolite concentrations changes were directly observed, while pathway and upstream regulator activation state are predicted based on metabolite data. Caret symbols (^) denote a Level 1-2 confidence identification from core facility panel confirmed by reference standard; Asterisks (*) denote a putatively annotated feature from LC-MS/MS fragmentation matches using the NIST MS Search software, earning Level 3+ identification confidence^25^.

Sepsis was associated with dysregulated nucleotide metabolism, including elevated levels of guanosine and adenine. This coincided with increased activation of the purine nucleotide de novo biosynthesis pathway, suggesting that cerebellar cells may be attempting to augment purine nucleotide production^30^ in response to stress or increased turnover. However, concurrent downregulation of the interconversion of nucleotide di- and triphosphates and overall nucleotide biosynthesis pathways suggest a bottleneck in converting intermediates into fully functional high-energy nucleotides such as ATP and GTP. This likely impairs both cellular energy homeostasis and nucleic acid synthesis. MSC-sEV treatment reversed these effects by up-regulating NDP ⇄ NTP interconversion and basal nucleotide biosynthesis, while suppressing nucleotide salvage and purine nucleotides de novo biosynthesis II pathways. These changes suggest a shift away from compensatory production routes toward restored homeostatic balance. Reductions in intermediate metabolites such as D-ribulose-5-phosphate and pyrophosphate further imply decreased flux through pathways typically activated under cellular stress conditions. These restorative effects may be partially mediated by MSC-sEV cargo proteins ATP6V1A^31^ and VCP^31^, which have been shown to regulate ATP-generating metabolic pathways and mitochondrial function^32^, thereby supporting improved nucleotide metabolism and energy homeostasis.

Sepsis impaired membrane transport and remodeling, as evidenced by down-regulations of APOA1, along with suppression of the transport of bile salts and organic acids, metal ions, and amine compounds pathway, and the choline catabolism pathway. These alterations are hallmarks of defective lipid clearance, detoxification, and phospholipid turnover that likely compromise membrane integrity and synaptic signaling. Following MSC-sEV treatment, we observed an upregulation of CUEDC2 expression which may suggest enhanced regulatory control over membrane-associated inflammatory responses. Additionally, the reactivation of transport pathways indicates an improved capacity for transmembrane movement of key metabolites and ions. Upregulation of the choline catabolism pathway following MSC-sEV treatment may reflect increased choline turnover to support membrane phospholipid remodeling. Notably, MSC-sEV cargo protein Annexin A1 may contribute to the modulation of membrane-associated inflammation^33, 34^, potentially acting in concert with CUEDC2 to regulate inflammatory responses. Concurrent decreases in glycerophosphocholine and citicoline^35^ may indicate reduced demand for acute membrane repair mechanisms.

With regard to nitrogen and amino metabolism, a pronounced shift towards hypercatabolism was evident in septic metabolic profiles. We found that concentrations of polyamines and 1-methyl-L-histidine were increased, alongside up-regulations of urea cycle, arginine biosynthesis IV, and citrulline-nitric oxide (NO) cycle pathways. These results suggest increased synthesis of NO, a key mediator of inflammatory and vasodilatory responses commonly implicated in sepsis pathogenesis. Up-regulation of the super pathway of methionine degradation indicates an increased breakdown of methionine^36^, contributing further to the catabolic state of septic cerebellar tissue. In parallel, a decrease in dopamine accompanied prediction down-regulation of CUEDC2^37^, sulfur amino acid metabolism, and glutamate/glutamine metabolism pathways, reflecting a potentially compromised antioxidant defense, while simultaneously reducing the substrates needed for anabolic processes or efficient energy transfer. MSC-sEV treatment appears to have rebalanced these processes via GATA4-driven transcriptional activation^38^ and up-regulating sulfur amino acid and glutamate/glutamine metabolism pathways to preserve glutathione, optimize nitrogen handling, and limit polyamine accumulation, urea cycle flux, arginine/NO production, and methionine degradation. The balanced inflammatory response associated with MSC-sEV treatment could be attributable to the actions of cargo such as miR-146 and miR-199a, known regulators of TNF-α, IL-6, and NF-κB signaling^17, 39, 40^, as well as protein Annexin A1^33, 34^.

Core energy and carbon metabolism pathways were severely impaired during sepsis. Marked downregulation of the TCA cycle and pyruvate metabolism pathways, accompanied by reduced availability of L-glucose, provides strong evidence that central pathways for ATP production and carbon utilization were compromised. Concurrent increases in intermediate metabolites such as N-oleoylglycine, 3-hydroxy-3-methylglutaryl-CoA, glucosamine, and adenine could suggest compensatory efforts via alternative substrates to sustain energy production and cell viability. MSC-sEVs reversed these deficits by up-regulating HIF-1α^41^ to defend the cell against oxidative damage by decreasing the OXPHOS and mitochondrial ROS production^42^.

AMP-activated protein kinase (AMPK; evidenced through subunit PRKAA1) was also up-regulated, driving mitochondrial biogenesis^43^ and mitophagy^42^ and resulting in a net increase in healthy cellular mitochondria. Here, another subset of MSC-sEV cargo, miR-30, Let-7, and miR-26, are likely to be responsible for these ameliorations on mitochondrial energetics and metabolic pathways, as they have all been shown to regulate mitochondrial dynamics and OXPHOS (with miR-30 also altering metabolic pathways to maintain lipid metabolism homeostasis)^15, 18–21, 44^. Cargo proteins ATP6V1A and VCP have also been associated with the regulation of ATP-generating pathways^31, 32^, with VCP also shown to support mitochondrial respiration^32^.

Predicted up-regulation of receptor kinase ERBB2 can drive activation of the PI3K/AKT signaling pathway, which serves to further up-regulate H1F-1α transcription^45–47^. These changes restored TCA cycle, pyruvate metabolism, and glyoxylate metabolism/glycine degradation. Down-regulation of N-oleoylglycine and GAPDH^48^ thereby attenuates TNF-α production and suppresses the gluconeogenesis I pathway, indicating that cells are no longer relying on emergency glucose production following the administration of the MSC-sEV treatment. Given that hyperglycemia is a frequent and clinically-concerning metabolic derangement observed in septic patients^49, 50^, this down-regulation may be indicative of a return to a basal gluconeogenesis state.

Redox homeostasis and co-factor metabolism were also impaired in sepsis. Key protective and regulatory elements, including EP300, CA9, ALDH2, and the sirtuin signaling cycle, were all downregulated in septic animals relative to controls. These components are critical for mounting an effective antioxidant response and maintaining redox balance^51, 52^. We also found a predicted up-regulation of the metabolism of water-soluble vitamins and cofactors and flavin biosynthesis IV pathways. Water-soluble vitamins, such as vitamins B and C, are essential for enzymatic reactions involved in energy metabolism, redox balance, and antioxidant defense which have shown promising potential as therapeutic targets for sepsis treatment in recent years^53^. Flavins, synthesized through the flavin biosynthesis IV pathway, function as critical cofactors (FAD and FMN) for various redox reactions within the mitochondria and cytoplasm, significantly influencing cellular energy production and antioxidant defense mechanisms^54^. Up-regulation of these pathways could reflect a compensatory response to severe oxidative stress and cellular damage. Notably, the SIRT5 upstream regulator was up-regulated in septic animals, in contrast to the above expected biological output and a recently published report^55^. This paradoxical increase may reflect a compensatory cellular response aimed at counteracting severe oxidative and metabolic stress through alternative regulatory pathways; however, it remains unclear whether this up-regulation is functionally protective or simply indicative of stress-induced dysregulation. Following MSC-sEV treatment, predicted activation of EP300^56, 57^ and SIRT2^58, 59^ may drive transcription of antioxidant genes and deacetylation of metabolic enzymes critical for ROS detoxification. We propose a support mechanism by way of MSC-sEV cargo such as miR-21 - which is overexpressed in neutrophils and macrophages in sepsis^16^ - and Let-7, providing additional reinforcement to these antioxidant defenses by attenuating oxidative stress and inflammatory signaling. Concurrent up-regulation of CA9 supports NADPH/NADP^+^ homeostasis and glutathione reductase activity^57^, while continued up-regulation of water-soluble vitamin/cofactor metabolic pathway suggests improved supply of essential vitamins and cofactors needed for redox enzymes.

Glutathione^60^, as well as its oxidized form oxiglutathione, was present in lower concentration in the tissue of treated mice relative to untreated, potentially indicative of alleviated oxidative burden, while down-regulation of OGT^61, 62^, the sirtuin signaling cycle pathway, and the flavin biosynthesis IV pathway could reflect similar diminished requirements for emergency or compensatory antioxidant responses.

Finally, sepsis triggered activation of core DNA repair mechanisms (e.g., mismatch repair) and innate immune detection (cytosolic sensors of pathogen-associated DNA), suggesting that cells are actively responding to DNA damage and heightened oxidative stress. The septic metabolic profile also describes the downregulation of several negative or modulatory components of immune regulation (including CUEDC2, immunoglobulin, and CSF3), suggesting impaired capacity to regulate inflammatory and stress responses. This imbalance may result in prolonged inflammatory signaling, exacerbating cellular stress and potentially contributing to tissue damage or dysfunction characteristic of sepsis. MSC-sEV treatment re-established homeostasis by up-regulating CUEDC2 to initiate the cessation of inflammation. Simultaneously, down-regulation of the mismatch repair pathway suggests a reduced need for DNA repair, consistent with reduced oxidative and inflammatory stress following treatment. Overall, these results suggest that MSC-sEV treatment is alleviating the DNA damage and inflammatory burden placed on cells by sepsis. These reductions in DNA damage responses and inflammatory signaling pathways may also reflect a specific mechanism of miRNA intervention, such as miR-146 and miR-199a, in addition to Annexin A1. Collectively, this cargo may be modulating critical inflammatory signaling cascades^17, 33, 34, 39^ and reducing oxidative stress-induced DNA damage, supporting an overall decrease in cellular stress and injury.

In summary, our proposed mechanistic models (see **Table 2**) highlight how MSC-derived sEV cargo, including specific miRNAs (miR-146a, miR-199a, miR-30, Let-7, miR-26, and miR21) and proteins (Annexin A1, ATP6V1A, VCP), may regulate key metabolic, inflammatory, and stress-response pathways during sepsis. These integrated pathways align with the treatment-associated improvements observed in the cerebellum, including reductions in oxidative and inflammatory stress, restoration of mitochondrial and energy metabolism, and decreased evidence of DNA damage, as revealed by our comprehensive metabolomics analyses. While the links reported between MSC-sEV cargo and observed metabolic profile are limited to correlation in this study, these results provide preliminary support for future studies to functionally validate causal relationships.

**Table 2.**
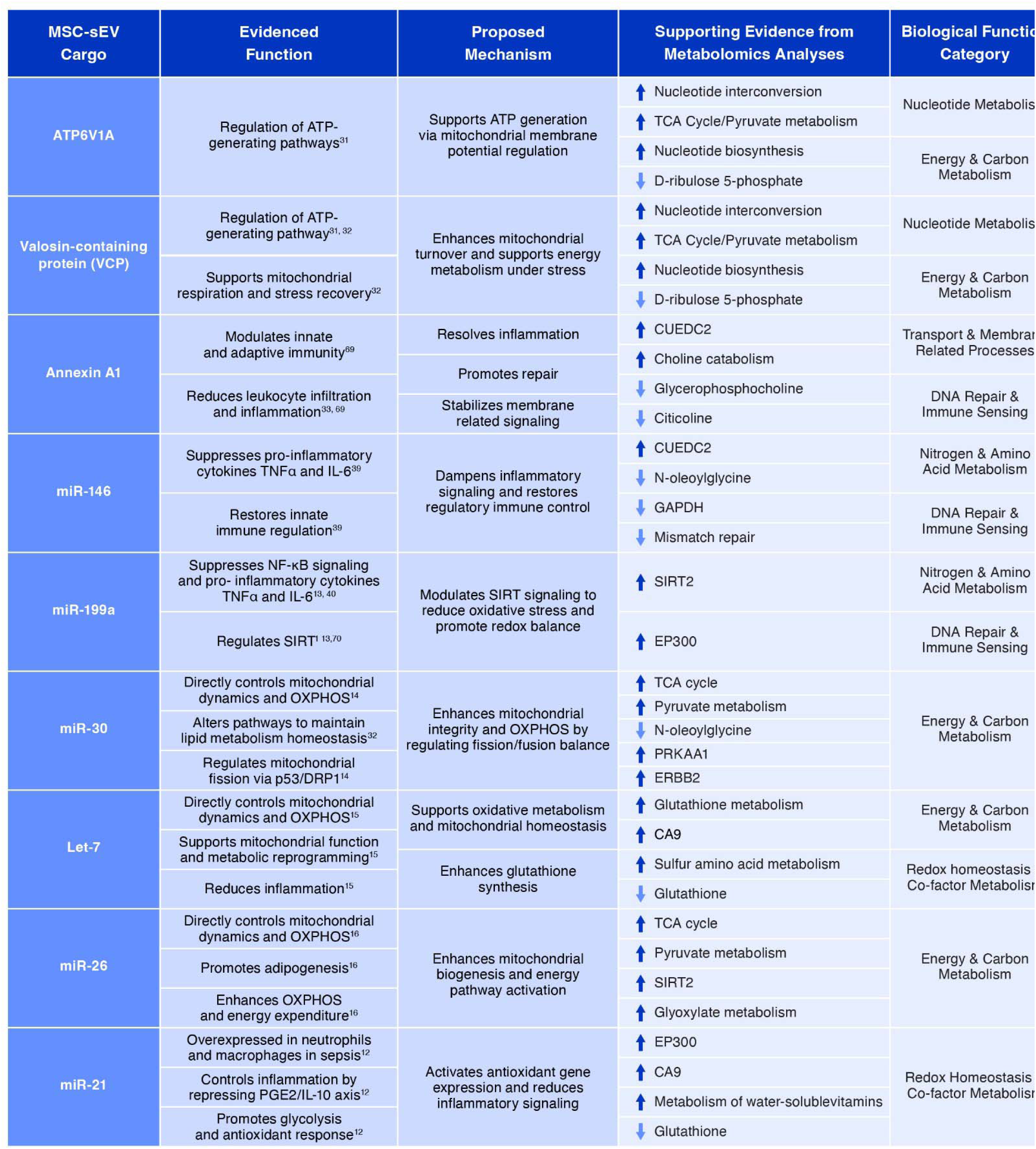
Summary of established MSC-sEV miRNA and protein bioactive cargo, with proposed mechanisms of influence in treated septic animals. For each cargo, known biological functions are listed alongside a proposed mechanism by which it may affect physiological processes in septic animals given MSC-sEV treatment. Supporting correlational evidence from our metabolomics data, by way of observed changes in metabolites or predicted alterations to upstream regulators and/or canonical pathways, is included along with the corresponding biological function category as previously defined in **Figure 7**.

Given these extensive disruptions to metabolic and inflammatory processes observed in cerebellar tissue, we next sought to identify peripheral SE biomarkers that could reliably reflect localized cerebellar distress and thus offer accessible, non-invasive diagnostic criteria for sepsis-associated injury. To this end, we implemented a cross-compartment screening strategy to identify plasma metabolites that reflect the metabolic signatures of sepsis-associated cerebellar distress.

We reanalyzed core comparisons using our high-confidence, targeted-only metabolomics data to evaluate convergence with findings from the comprehensive dataset. We observed a high level of convergence between the two datasets in cerebellar tissue. Slight differences in the targeted-only dataset emphasized redox balance, mitochondrial energetics, and amino acid metabolism, while signals related to lipid signaling and nucleotide repair functions prominent in the untargeted data were reduced. This likely reflects a stronger representation of oxidative stress and energy metabolism metabolites in the targeted metabolomics panel, which were preserved even under the more restrictive inclusion criteria. In contrast, plasma results illustrated a decidedly lower consistency between datasets, likely attributable to a reduced overlap in the targeted panel and greater biological heterogeneity relative to cerebellum. Collectively, this reanalysis reinforces the validity of our core conclusions and supports the reliability of our novel comprehensive profiling strategy, while clarifying the distinct contributions of our targeted and untargeted metabolomic approaches.

This study is the first to identify distinct cross-compartment metabolites with strong promise as biomarkers of sepsis-associated CNS injury in mice while simultaneously providing human clinical evidence of those metabolite concentrations being consistently altered, supporting efforts towards earlier diagnosis. In parallel, we provide a comprehensive characterization of metabolic alterations in sepsis-associated cerebellar injury and demonstrates the therapeutic potential of MSC-sEVs in mitigating these effects. We propose that their benefits may be mediated through miRNA and protein cargo. Future work is necessary to define the specific contributions of individual MSC-sEV cargo components experimentally. Clarifying the functional roles of MSC-sEV cargo will be essential for understanding the mechanisms by which this treatment exerts its effects. Such insights will advance the development of MSC-sEV-based therapies and support the refinement of targeted interventions for sepsis-related CNS injury. In total, the results of this study identify biomarker candidates to aid in early detection and improved clinical management of sepsis and establish a foundation for mechanistic and translational advancements in sepsis-associated CNS injury by defining cerebellar metabolic disruptions caused by sepsis, demonstrating the therapeutic impact of MSC-sEV treatment in reversing these disruptions, providing correlational support for future causal investigations into MSC-sEV-based mechanisms of action.

### Limitations

We employed a well-established cecal slurry model of acute murine sepsis with strong construct validity designed to closely approximate key features of human sepsis. A key limitation of this model is the omission of clinical standard-of-care interventions, including antibiotic administration and IV fluid resuscitation. While it may impact the translatability of our findings, this omission was made intentionally to avoid confounding the therapeutic effects of MSC-derived sEVs. Future studies will incorporate standard-of-care analogs to better model real-world clinical conditions.

We also acknowledge limitations inherent to using using putative metabolite annotations based on spectral matching from LC-MS/MS fragmentation databases. To ensure transparency, we reported annotation confidence levels in the Results section and figures. We also performed parallel analyses of all core comparisons using only high-confidence targeted metabolomics data to validate our comprehensive approach findings, including those results in the supplemental materials. For detailed criteria regarding annotation confidence levels and field standards, we refer readers to the framework proposed by Schymanski et al^25^.

## ACKNOWLEDGEMENTS

The research reported in this manuscript was supported by NINDS (1R21NS135088 awarded to IK). We thank Dr. Amrita Cheema, Dr. Shivani Bansal, and Meth Jayatilake from Georgetown University Medical School Lombardi Comprehensive Cancer Center for mass spectrometry services and interpretation support. We thank Dr. John W. Ludlow from Zen-Bio Inc. for providing us with electron microscopy image and protein characterization of MSC-derived sEVs. We thank Dr. Sean Taylor and Research Scientific Computing at Seattle Children’s Research Institute for training and support with the HPC and bioinformatic analyses. We thank Christina Corbaci for graphic illustration design and support. We also thank Dr. Kritika Bhalla, Julian Naizaque, Dr. Alejandro Parga-Becerra, Dr. Emma Parkins, and Olivia Santiago from the Gallo Lab in the Norcliffe Center for Integrative Brain Research at Seattle Children’s Research Institute for their input on drafts of this manuscript.

## Declaration of generative AI and AI-assisted technologies in the writing process

During the preparation of this work, the authors used ChatGPT-4o from OpenAI for language suggestions and editing. After using this tool, the authors reviewed and edited the content as needed and takes full responsibility for the content of the publication.

