## Supplemental Materials for "Cross-species biomarkers of cerebellar injury in acute sepsis: integrated metabolomics of mouse cerebellum and plasma identify conserved human plasma biomarkers and therapeutic modulation by human MSC-sEVs"

**Supplemental RESULTS**

**Overlapping base peak chromatograms and strong quality control correlations confirmed mass spectrometry data reliability.**

Quality control was assessed through overlapping base peak chromatograms (BPC) plot of all QC samples for each mass spectrometry analysis. In general, all pooled QC samples showed strongly overlapping plots with small fluctuations in retention times and corresponding peak intensities, suggesting that the detection instruments were functioning well during sample detection and analysis (see **Supplemental Figures 1a,c – 4a,c** for overlapping BPCs from each mass spectrometry analysis). We determined that each QC was significantly correlated to one another within their specific mass spectrometry analysis using pairwise comparisons (all p-values < 0.05). Cerebellar LC-MS QCs correlations ranged from 0.92 to 0.99 for NEG QCs and 0.90 to 0.99 for POS QCs (**Supplemental Figure 1b,d**). Cerebellar LC-MS/MS QCs correlations ranged from 0.76 to 0.99 for NEG QCs and 0.64 to 0.99 for POS QCs **(Supplemental Figure 2b,d)**. Plasma LC-MS QCs correlations ranged from 0.98 to 0.99 for NEG QCs and 0.98 to 0.99 for POS QCs (**Supplemental Figure 3b,d**). Plasma LC-MS/MS QCs correlations ranged from 0.65 to 0.99 for NEG QCs and 0.02 to 0.99 for POS QCs (**Supplemental Figure 4b,d**).

**Sepsis (-) v. Control (-) comparison isolating disease state in plasma.**

We found a PLS-DA model with strong separation between Sepsis (-) and Control (-) groups (**Supplemental Figure 5a**) where Component 1 explains 48.7% of the variance between groups and Component 2 explains 17.5%. This model was optimized for five components, resulting in an average classification accuracy of 1.000, $R^2 = 0.998$, and $Q^2 = 0.941$. We then identified the top 10 metabolites according to VIP score (ranging from 1.94 – 2.93; see **Supplemental Figure 5b**). Top 10 metabolites include: ^Lipoamide, *Glycolithocholic acid, *Tetradecanoylcarnitine, *3-Hydroxyoleylcarnitine, *Oleoylcarnitine, *Palmitoyl Carnitine, *Glucosylshingosine, *N-acetylleucyl-leucyl-methioninal, ^S-Adenosylhomocysteine, and ^L-Ornithine. All metabolites except ^S-Adenosylhomocysteine were increased in septic plasma relative to controls (see **Supplemental Figures 5c-l**).

Pathway analyses resulted in a total of 863 canonical pathways predicted to be altered between disease states based on the annotated metabolites and relative concentrations measures. **Supplemental Figure 6a** displays all pathways predicted to be altered between disease states. We further explored the nature of the top 20 pathways from each comparison (illustrated in **Supplemental Figure 6b**) as determined by Pathway Impact score.

The top 10 upstream regulators identified in plasma were CPT1B, MYO1C, GNMT, SIX1, CA9, MYC, CCND1, CRBN, GATA4, BCKDHA. These regulators were predicted to be up-regulated in septic animals compared to controls, except for CPT1B, MYO1C, GNMT, and CRBN, which are predicted to be down-regulated in septic animals (**Supplemental Figure 6c**).

**Sepsis (+) v. Sepsis (-) comparison isolating treatment condition in plasma.**

We constructed a predictive model by way of PLS-DA, where Component 1 explains 51.1% of the variance between groups and Component 2 explains 7.4% (**Supplemental Figure 5m**). For a PLS-DA model with 2 components, we found an accuracy measure of 0.60, an $R^2$ value of 0.812, and a $Q^2$ value of -0.223, indicating poor model performance. Top 10 metabolites driving differences between groups were then identified (VIP scores ranging from 2.06 – 5.31; see **Supplemental Figure 5n**) and include: ^Lipoamide, ^Ethanolamine, *Proadrenomedullin (1-20), *Pancreastatin, *1-Palmitoyl-2-oleoyl-sn-glycero-3-phosphoethanolamine, ^3-(4-hydroxyphenyl)pyruvate, ^Guanosine monophosphate (GMP), ^Adenosine monophosphate (AMP), ^Normetanephrine, and *Glucagon (1-29; see **Supplemental Figure 5o-x**).

Pathway analyses resulted in a total of 742 pathways based on the annotated metabolites and relative concentrations measures. **Supplemental Figure 6d** displays all pathways predicted to be altered between treatment states. We further explored the nature of the top 20 pathways from each comparison (illustrated in **Supplemental Figure 6e**) as determined by Pathway Impact score. Upstream regulators CCND1, CD40, MYC, and SIX1 were predicted to be up-regulated in treated animals relative to non-treated, while down-regulation was predicted in CPT1B, TGFB1, GNMT, IL37, and IL1B (**Supplemental Figure 6f**).

**Supplemental FIGURES**

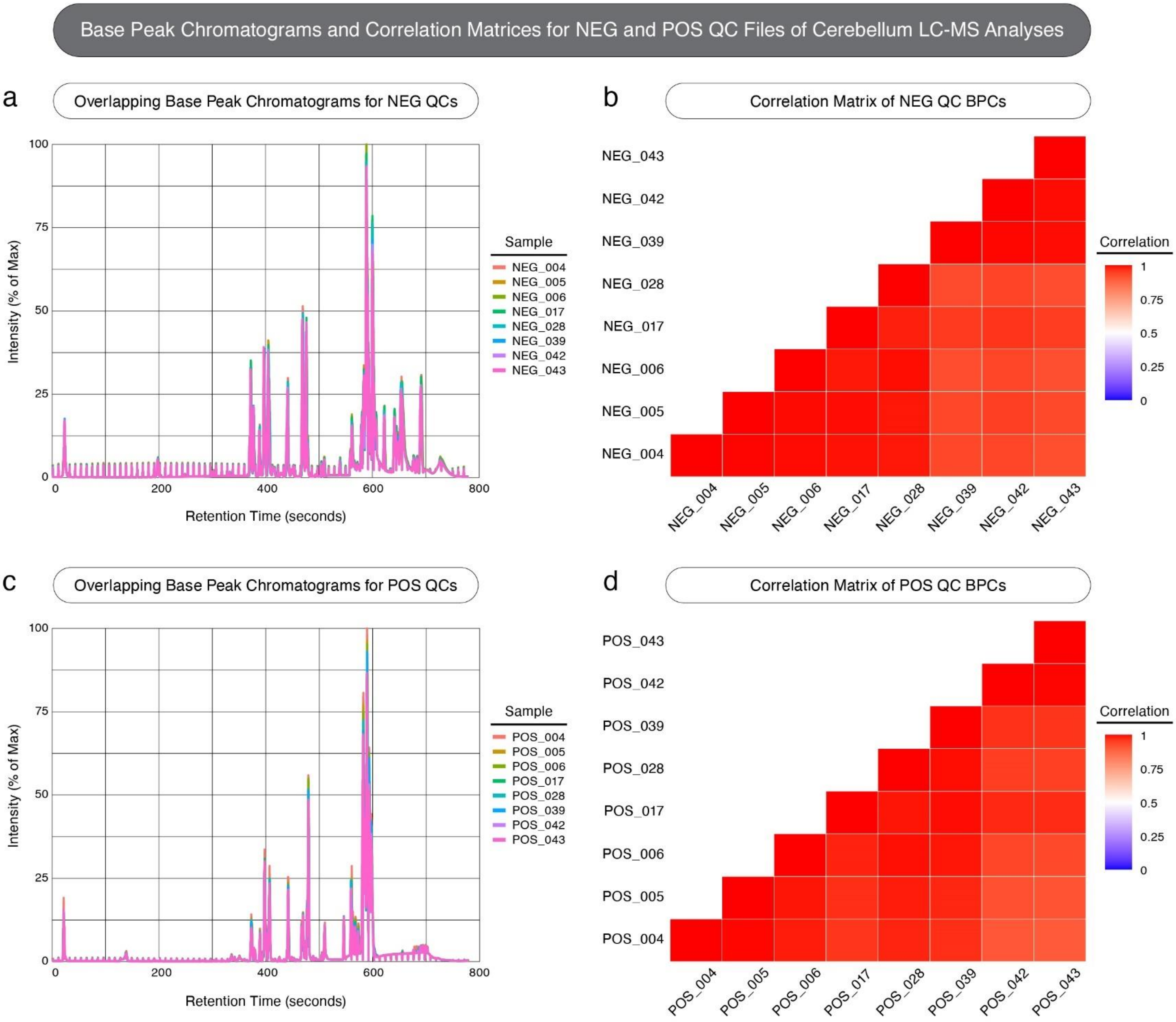

**Supplemental Figure 1**. **Base peak chromatograms (BPCs) and correlated matrices for quality control (QC) samples from cerebellar LC-MS analyses in negative and positive ion modes.** This figure serves as a quality control assessment to evaluate analytical stability and reproducibility across QC runs. (**a**) BPCs for negative ion mode (NEG) QC samples, with normalized intensity (% of maximum; y-axis) plotted against retention time (x-axis). Each trace represents a separate QC injection throughout data collection. (**b**) Pearson correlation matrix for NEG QC samples, showing pairwise correlation coefficients between QC injections. (**c**) BPCs for positive ion mode (POS) QC samples. (**d**) Pearson correlation matrix for POS QC samples. Warmer colors represent higher correlation coefficients, cooler colors represent lower correlation coefficients. Consistent retention times and strong within-ionization mode correlations highlight strong stability and reproducibility of QC measures and suggest strong data quality from mass spectrometry.

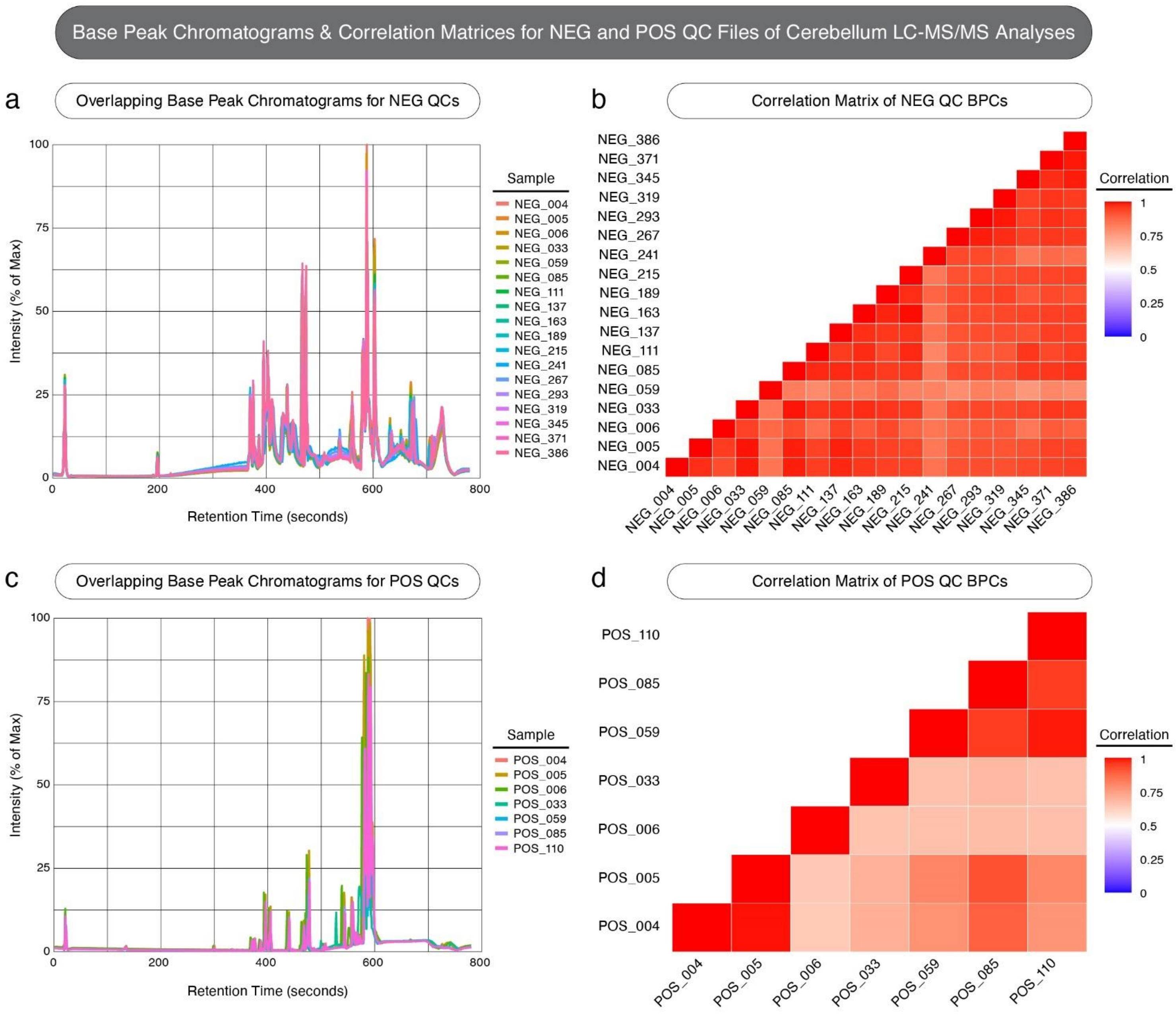

**Supplemental Figure 2. Base peak chromatograms (BPCs) and correlated matrices for quality control (QC) samples from cerebellar LC-MS/MS analyses in negative and positive ion modes.** This figure serves as a quality control assessment to evaluate analytical stability and reproducibility across QC runs. (**a**) BPCs for negative ion mode (NEG) QC samples, with normalized intensity (% of maximum; y-axis) plotted against retention time (x-axis). Each trace represents a separate QC injection throughout data collection. (**b**) Pearson correlation matrix for NEG QC samples, showing pairwise correlation coefficients between QC injections. (**c**) BPCs for positive ion mode (POS) QC samples. (**d**) Pearson correlation matrix for POS QC samples. Warmer colors represent higher correlation coefficients, cooler colors represent lower correlation coefficients. Consistent retention times and strong within-ionization mode correlations highlight strong stability and reproducibility of QC measures and suggest strong data quality from tandem mass spectrometry.

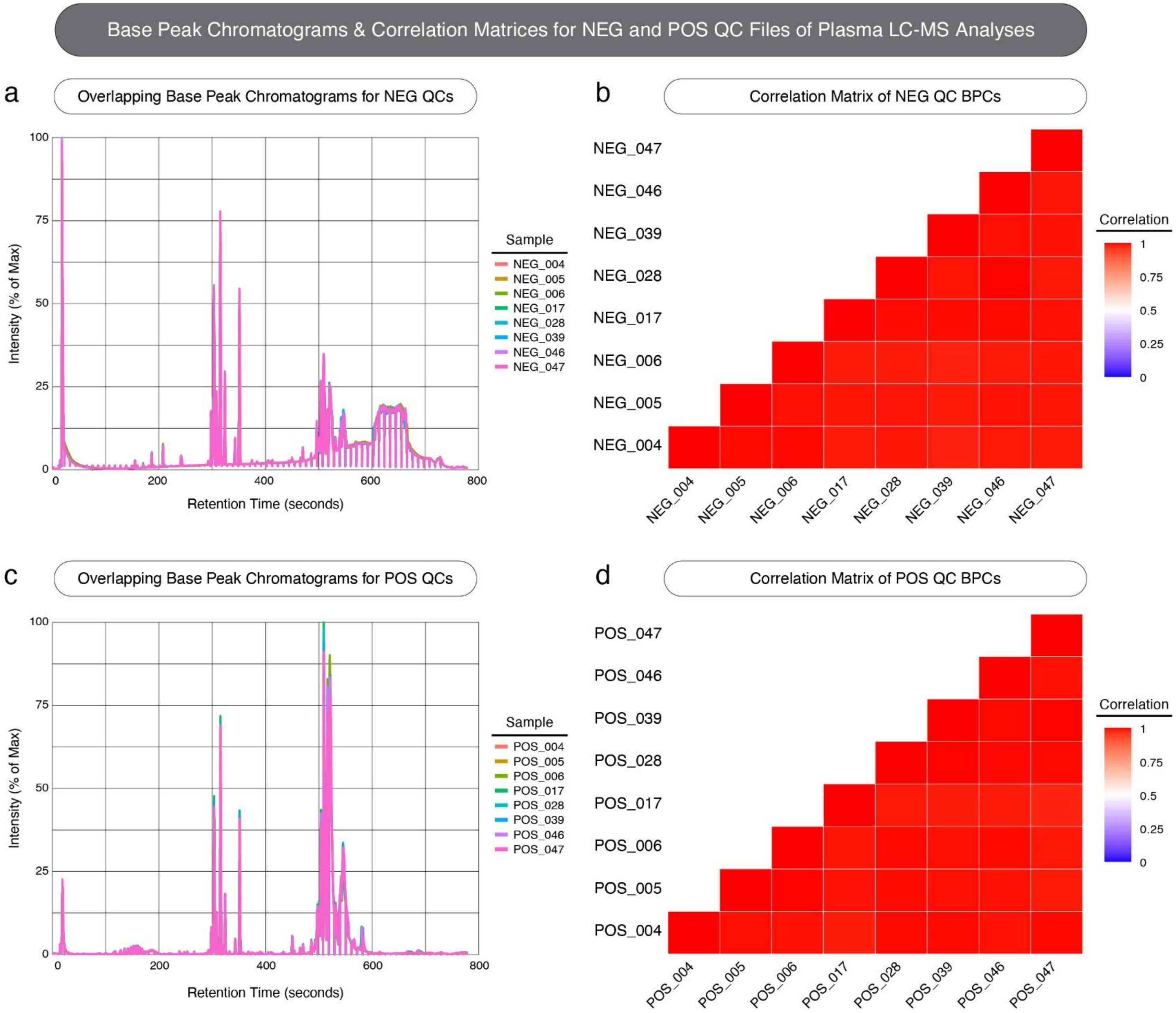

**Supplemental Figure 3**. **Base peak chromatograms (BPCs) and correlated matrices for quality control (QC) samples from plasma LC-MS analyses in negative and positive ion modes.** This figure serves as a quality control assessment to evaluate analytical stability and reproducibility across QC runs. (**a**) BPCs for negative ion mode (NEG) QC samples, with normalized intensity (% of maximum; y-axis) plotted against retention time (x-axis). Each trace represents a separate QC injection throughout data collection. (**b**) Pearson correlation matrix for NEG QC samples, showing pairwise correlation coefficients between QC injections. (**c**) BPCs for positive ion mode (POS) QC samples. (**d**) Pearson correlation matrix for POS QC samples. Warmer colors represent higher correlation coefficients, cooler colors represent lower correlation coefficients. Consistent retention times and strong within-ionization mode correlations highlight strong stability and reproducibility of QC measures and suggest strong data quality from mass spectrometry.

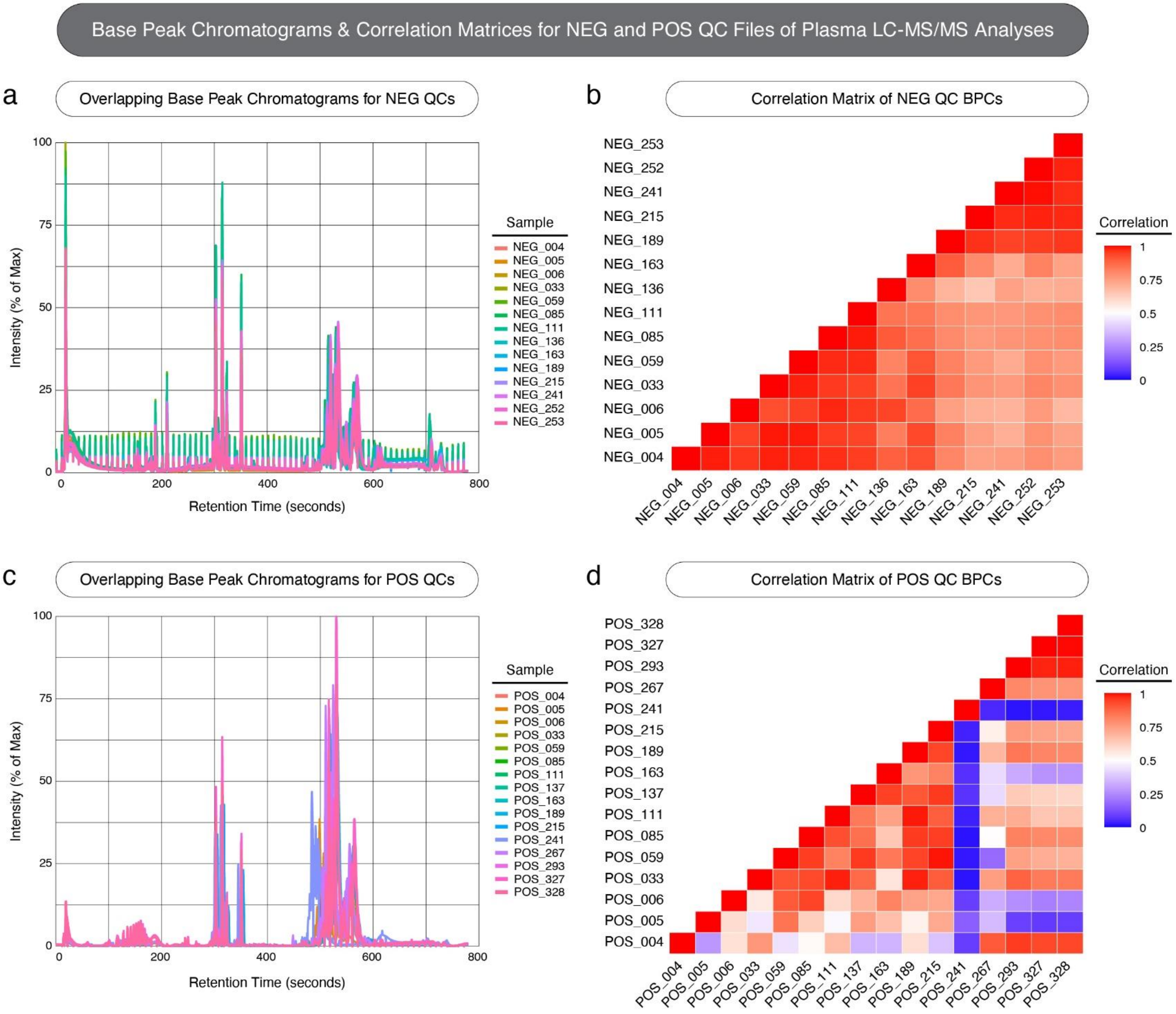

**Supplemental Figure 4. Base peak chromatograms (BPCs) and correlated matrices for quality control (QC) samples from plasma LC-MS/MS analyses in negative and positive ion modes.** This figure serves as a quality control assessment to evaluate analytical stability and reproducibility across QC runs. (**a**) BPCs for negative ion mode (NEG) QC samples, with normalized intensity (% of maximum; y-axis) plotted against retention time (x-axis). Each trace represents a separate QC injection throughout data collection. (**b**) Pearson correlation matrix for NEG QC samples, showing pairwise correlation coefficients between QC injections. (**c**) BPCs for positive ion mode (POS) QC samples. (**d**) Pearson correlation matrix for POS QC samples. Warmer colors represent higher correlation coefficients, cooler colors represent lower correlation coefficients. Consistent retention times and strong within-ionization mode correlations highlight strong stability and reproducibility of QC measures and suggest strong data quality from tandem mass spectrometry in NEG mode, but potential variability among individual POS QC injections.

| | Cerebellum | Plasma |
|---|---|---|
| Features detected in LC-MS | 4024 NEG, 5526 POS | 5187 NEG, 519 POS |
| Significant features from LC-MS | 463 | 671 |
| Features annotated from LC-MS/MS | 174 | 373 |
| Features from panel data | 324 | 324 |
| Total annotated metabolites | 498 | 697 |

**Supplemental Table 1. Summary of feature detection, quantification, and metabolite annotation in cerebellar tissue and plasma.** This table outlines the number of features detected in negative and positive ionization modes, the number of statistically significant features identified in initial LC-MS scans, and the number of metabolites successfully annotated in both cerebellar and plasma samples. Annotations are further broken down by targeted panel-based and LC-MS/MS-based spectral matching identifications. The final row indicates the total number of unique annotated metabolites per tissue.

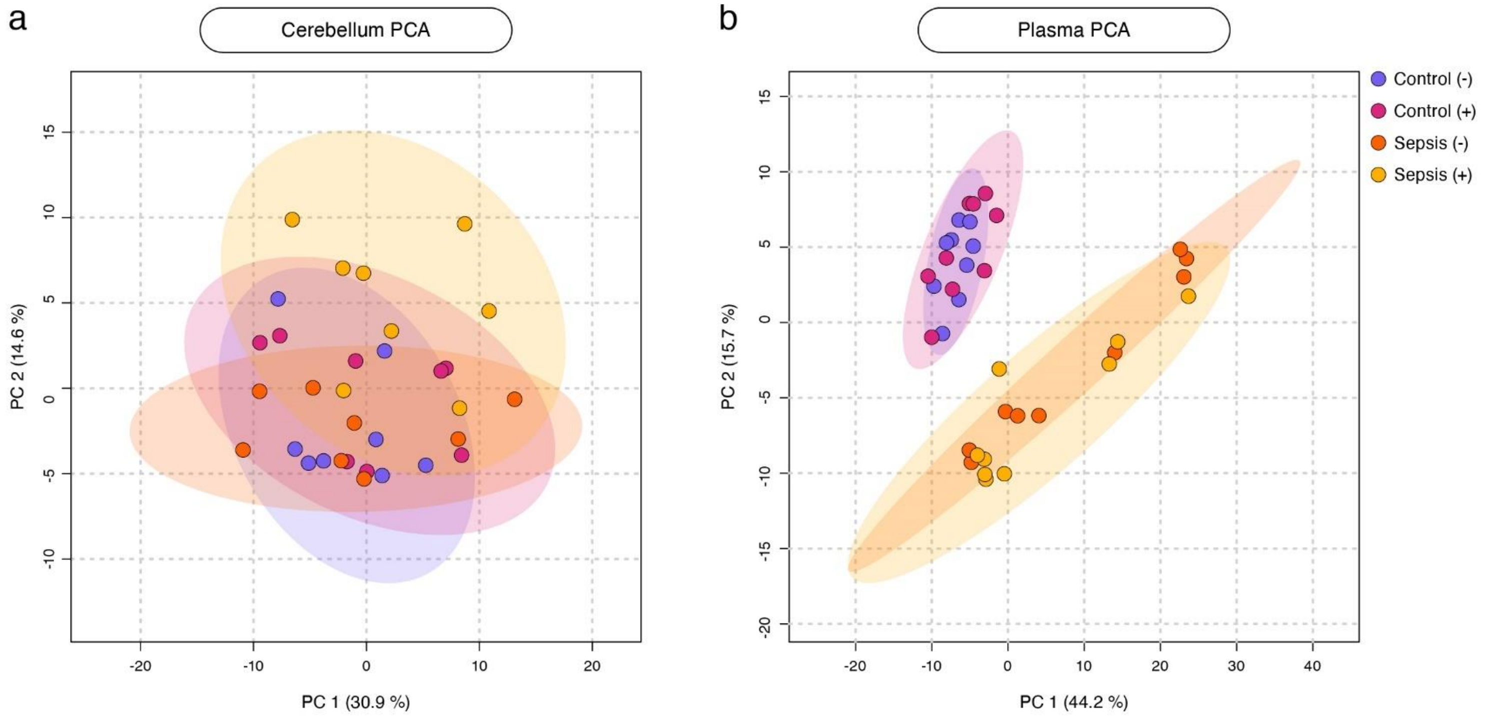

**Supplemental Figure 5. Principal component analyses (PCAs) of cerebellar tissue and plasma metabolite profiles between all experimental groups.** (**a**) PCA plot of cerebellar tissue samples showing the distribution of individual samples across the first two principal components (PC1 and PC2) as determined by percent of variance explained. (**b**) PCA plot of resulting from plasma samples. Each point represents one sample, colored corresponding to its experimental condition. Shaded ellipses represent a 95% confidence interval for each cluster. Axes represent the percentage of variance explain by PC1 (x-axis) and PC2 (y-axis).

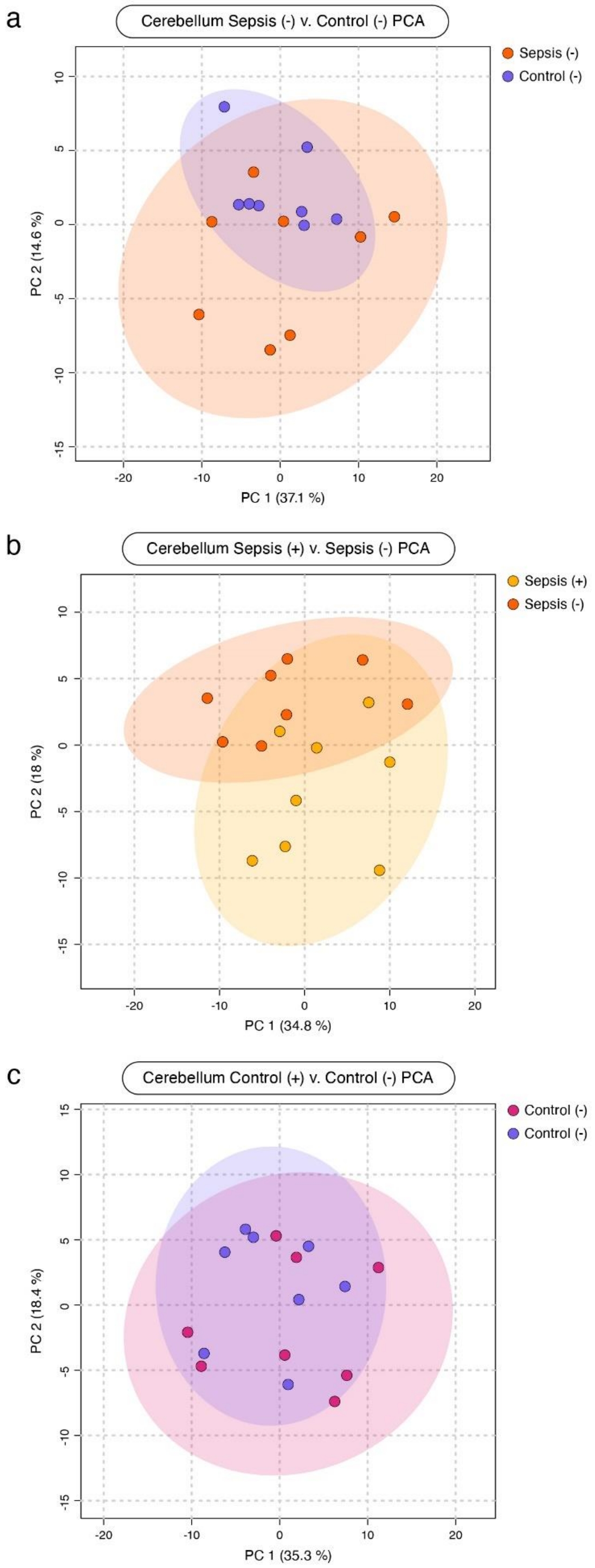

**Supplemental Figure 6. Principal component analyses (PCA) of metabolite profiles from cerebellar tissue between key pairwise comparisons of disease, treatment, and baseline MSC-sEV conditions.** (**a**) PCA plot of the disease state comparison, Sepsis (-) v. Control (-), in cerebellar tissue samples showing the distribution of individual samples across the first two principal components (PC1 and PC2) as determined by percent of variance explained. (**b**) PCA plot resulting from the treatment comparison, Sepsis (+) v. Sepsis (-). (**c**) PCA plot resulting from the baseline MSC-sEV comparison, Control (+) v. Control (-). Each point represents one sample, colored corresponding to its experimental condition. Shaded ellipses represent a 95% confidence interval for each cluster. Axes represent the percentage of variance explain by PC1 (x-axis) and PC2 (y-axis).

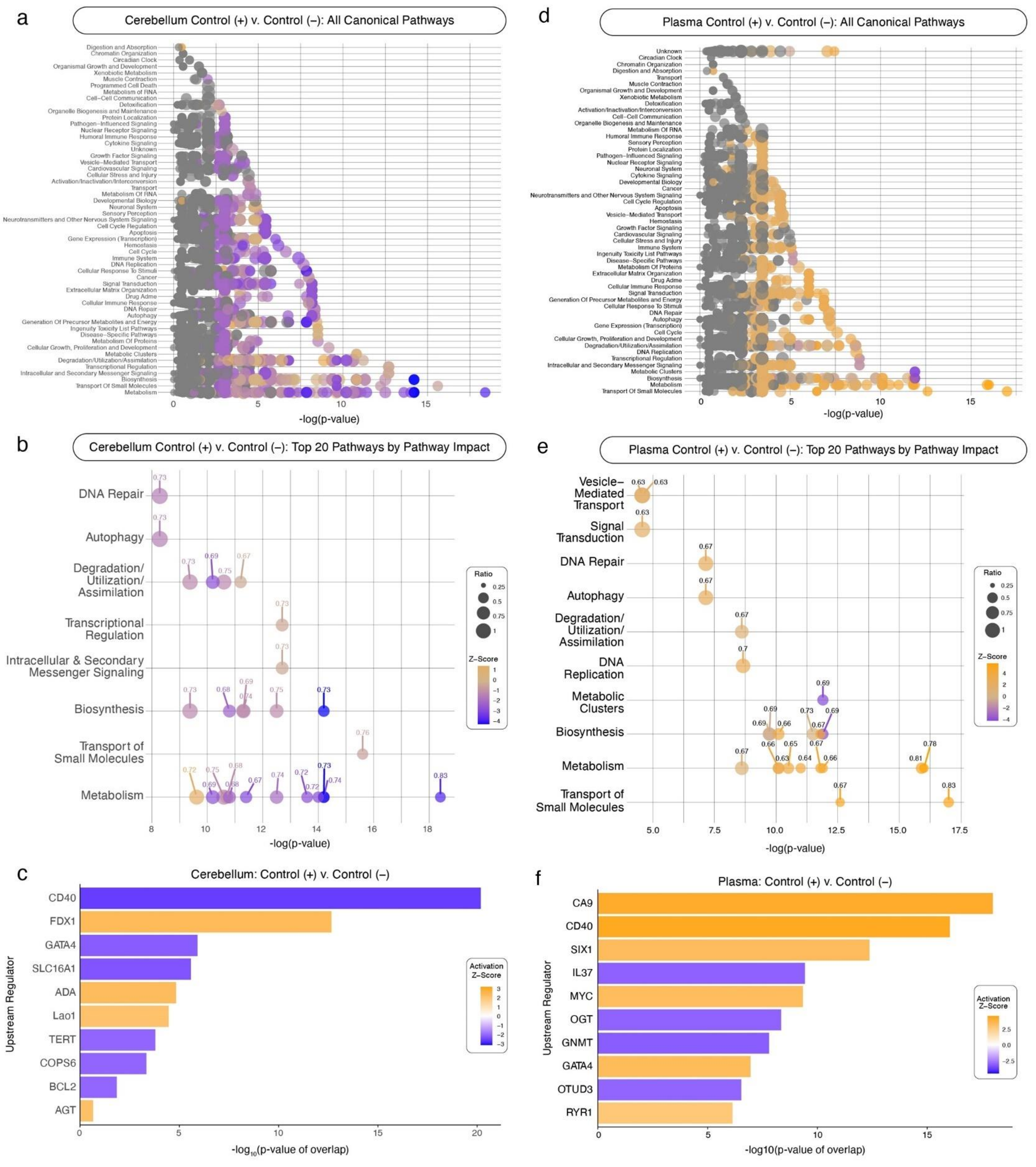

**Supplemental Figure 7**. **Canonical pathway activity and upstream regulator predictions resulting from treatment in control animals considering both cerebellum and plasma** . Panels **a-c** focus on treatment effect, Control (+) v. Control (-), seen in cerebellar tissue, while panels **d-f** focus on plasma. (**a,d**) Canonical pathway activity showing all pathways predicted to be affected as determined by metabolite intensities. Pathways are grouped by their biological function (y-axis) and plotted against their alteration significance (-log$_{10}$[p-value]; x-axis). Bubble size reflects the proportion of the measured metabolites relative to total known pathway metabolite profile, while bubble color represents the predicted activation direction and intensity (orange = predicted up-regulation, blue = predicted down-regulation). (**b,e**) Top 20 canonical pathways as ranked by Pathway Impact score. (**c,f**) Top 10 predicted upstream regulators influenced by each condition based on metabolite representation. Each regulator (y-axis) is displayed as a bar with a length representing its -log$_{10}$(p-value of overlap; x-axis) and is colored according to its predicted activation direction and intensity.

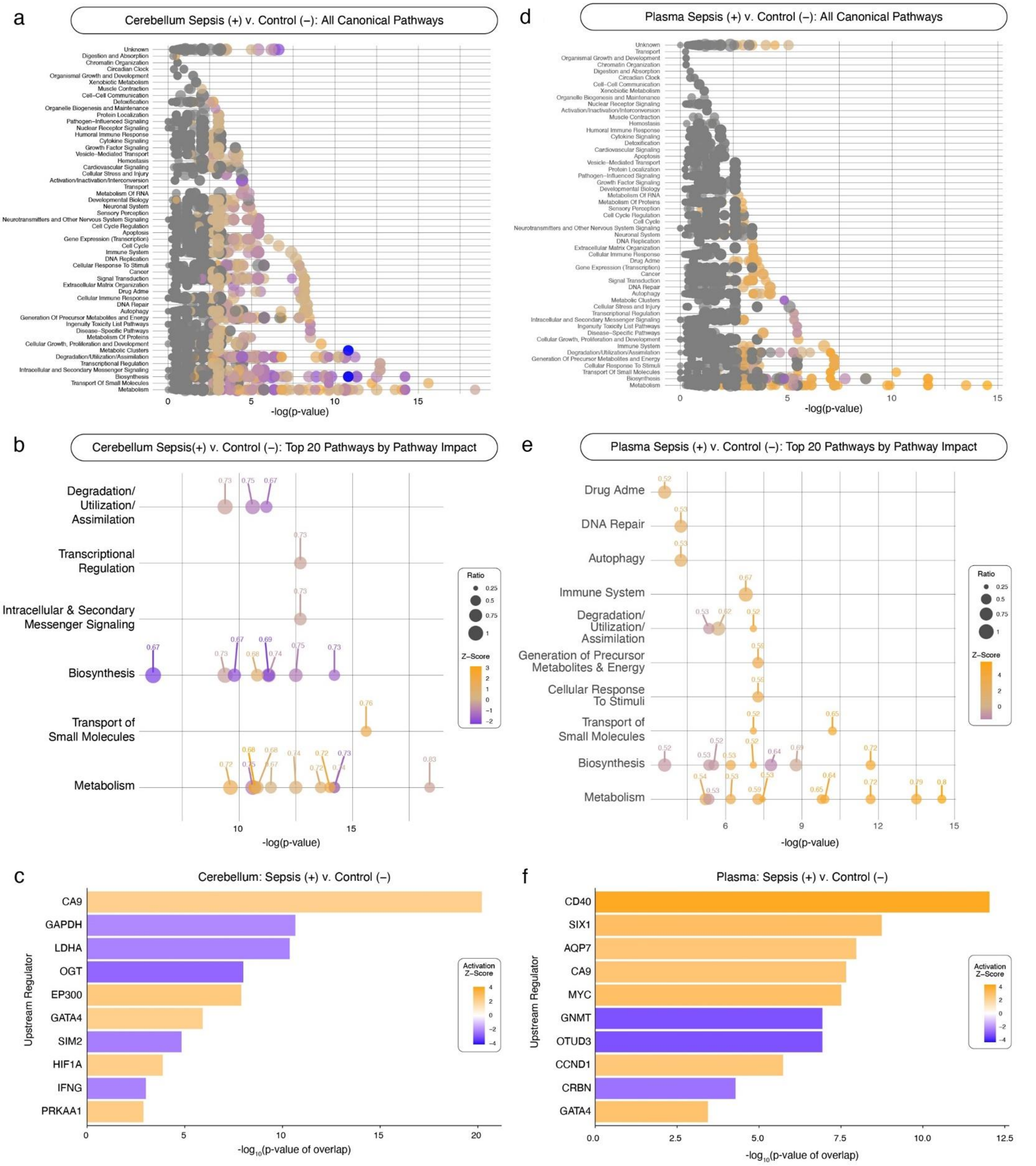

**Supplemental Figure 8**. **Canonical pathway activity and upstream regulator predictions resulting from Sepsis (+) v. Control (-) comparisons considering both cerebellum and plasma** . Panels **a-c** focus on the Sepsis (+) v. Control (-) comparison in cerebellar tissue, while panels **d-f** focus on plasma. (**a,d**) Canonical pathway activity showing all pathways predicted to be affected as determined by metabolite intensities. Pathways are grouped by their biological function (y-axis) and plotted against their alteration significance (-log$_{10}$[p-value]; x-axis). Bubble size reflects the proportion of the measured metabolites relative to total known pathway metabolite profile, while bubble color represents the predicted activation direction and intensity (orange = predicted up-regulation, blue = predicted down-regulation). (**b,e**) Top 20 canonical pathways as ranked by Pathway Impact score. (**c,f**) Top 10 predicted upstream regulators influenced by each condition based on metabolite representation. Each regulator (y-axis) is displayed as a bar with a length representing its -log$_{10}$(p-value of overlap; x-axis) and is colored according to its predicted activation direction and intensity.

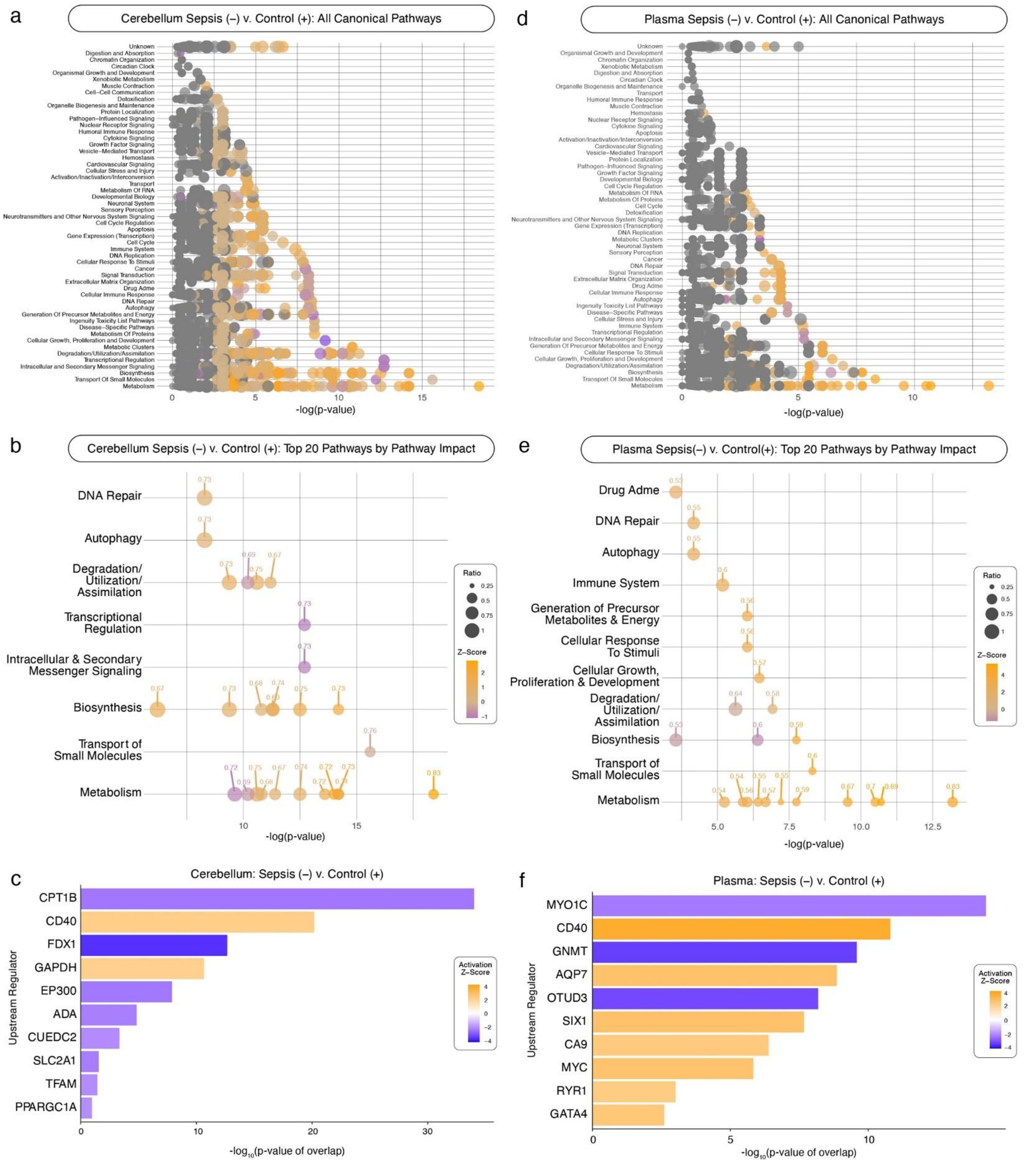

**Supplemental Figure 9**. **Canonical pathway activity and upstream regulator predictions resulting from Sepsis (-) v. Control (+) comparisons considering both cerebellum and plasma** . Panels **a-c** focus on the Sepsis (-) v. Control (+) comparison in cerebellar tissue, while panels **d-f** focus on plasma. (**a,d**) Canonical pathway activity showing all pathways predicted to be affected as determined by metabolite intensities. Pathways are grouped by their biological function (y-axis) and plotted against their alteration significance (-log$_{10}$[p-value]; x-axis). Bubble size reflects the proportion of the measured metabolites relative to total known pathway metabolite profile, while bubble color represents the predicted activation direction and intensity (orange = predicted up-regulation, blue = predicted down-regulation). (**b,e**) Top 20 canonical pathways as ranked by Pathway Impact score. (**c,f**) Top 10 predicted upstream regulators influenced by each condition based on metabolite representation. Each regulator (y-axis) is displayed as a bar with a length representing its -log$_{10}$(p-value of overlap; x-axis) and is colored according to its predicted activation direction and intensity.

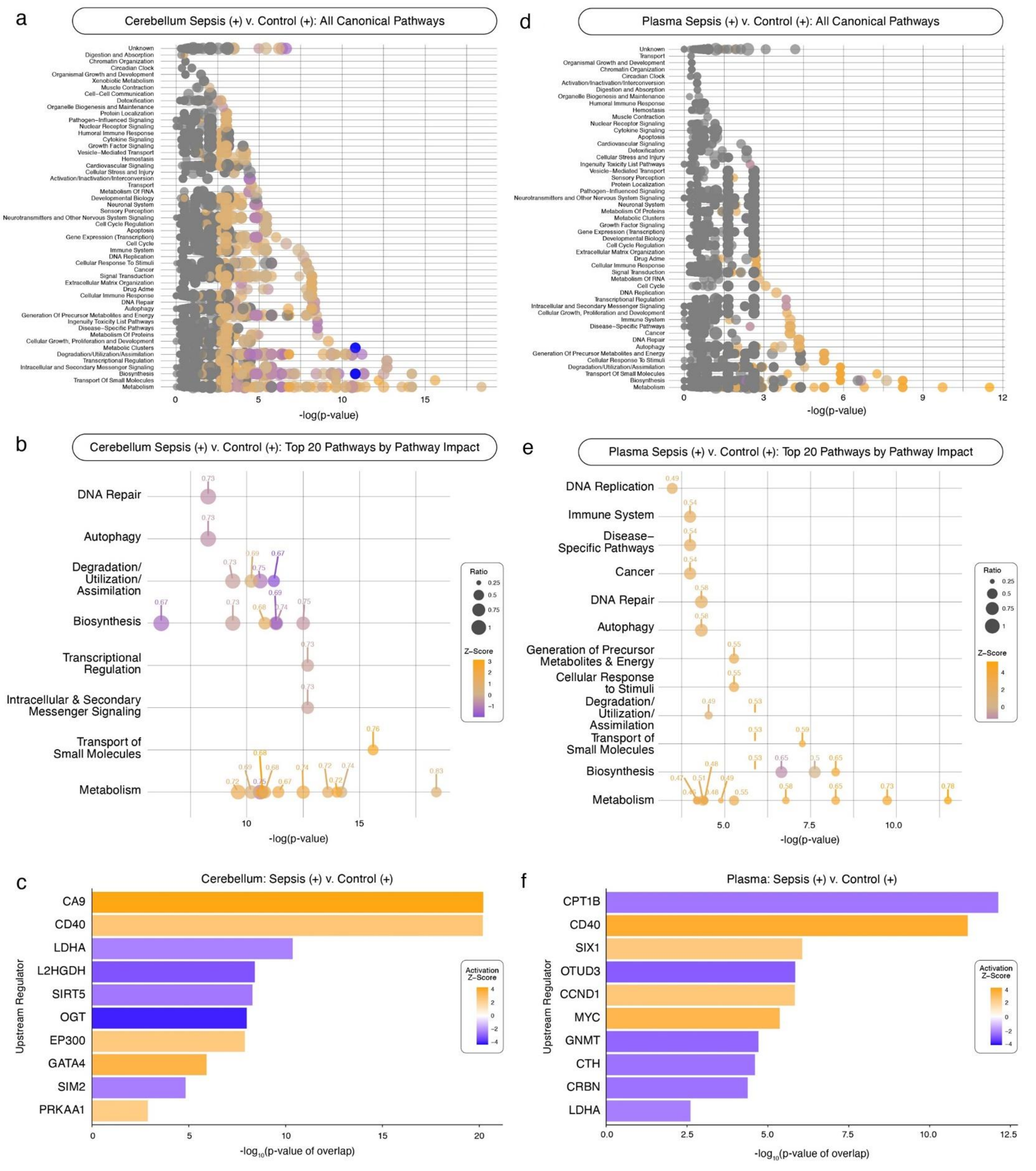

**Supplemental Figure 10**. **Canonical pathway activity and upstream regulator predictions resulting from Sepsis (+) v. Control (+) comparisons considering both cerebellum and plasma** . Panels **a-c** focus on the Sepsis (+) v. Control (+) comparison in cerebellar tissue, while panels **d-f** focus on plasma. (**a,d**) Canonical pathway activity showing all pathways predicted to be affected as determined by metabolite intensities. Pathways are grouped by their biological function (y-axis) and plotted against their alteration significance (-log$_{10}$[p-value]; x-axis). Bubble size reflects the proportion of the measured metabolites relative to total known pathway metabolite profile, while bubble color represents the predicted activation direction and intensity (orange = predicted up-regulation, blue = predicted down-regulation). (**b,e**) Top 20 canonical pathways as ranked by Pathway Impact score. (**c,f**) Top 10 predicted upstream regulators influenced by each condition based on metabolite representation. Each regulator (y-axis) is displayed as a bar with a length representing its -log$_{10}$(p-value of overlap; x-axis) and is colored according to its predicted activation direction and intensity.

| | Cerebellum | | | | | | | | | | | |
|---|---|---|---|---|---|---|---|---|---|---|---|---|
| | Control (+) v. Control (-) | | | Sepsis (+) v. Control (-) | | | Sepsis (-) v. Control (+) | | | Sepsis (+) v. Control (+) | | |
| Rank | Pathway | Biological Function Category | Pathway Impact | Pathway | Biological Function Category | Pathway Impact | Pathway | Biological Function Category | Pathway Impact | Pathway | Biological Function Category | Pathway Impact |
| 1 | Nucleotide Catabolism | Metabolism | -0.83 | Nucleotide Catabolism | Metabolism | -0.83 | Nucleotide Catabolism | Metabolism | +0.83 | Nucleotide Catabolism | Metabolism | +0.83 |
| 2 | Transport of bile salts & organic acids, metal ions & amine compounds | Transport of small molecules | -0.76 | Transport of bile salts & organic acids, metal ions & amine compounds | Transport of small molecules | +0.76 | Transport of bile salts & organic acids, metal ions & amine compounds | Transport of small molecules | -0.76 | Transport of bile salts & organic acids, metal ions & amine compounds | Transport of small molecules | +0.76 |
| 3 | Superpathway of Citrulline Metabolism | Biosynthesis | -0.75 | Superpathway of Citrulline Metabolism | Biosynthesis | -0.75 | Superpathway of Citrulline Metabolism | Biosynthesis | +0.75 | Superpathway of Citrulline Metabolism | Biosynthesis | -0.75 |
| 4 | Urea Cycle | Metabolism, Degradation/Utilization/Assimilation | -0.75 | Urea Cycle | Metabolism, Degradation/Utilization/Assimilation | -0.75 | Urea Cycle | Metabolism, Degradation/Utilization/Assimilation | +0.75 | Urea Cycle | Metabolism, Degradation/Utilization/Assimilation | -0.75 |
| 5 | Citric acid cycle (TCA cycle) | Metabolism | -0.74 | Citric acid cycle (TCA cycle) | Metabolism | +0.74 | Citric acid cycle (TCA cycle) | Metabolism | +0.74 | Citric acid cycle (TCA cycle) | Metabolism | +0.74 |
| 6 | Arginine Biosynthesis IV | Biosynthesis | -0.74 | Arginine Biosynthesis IV | Biosynthesis | -0.74 | Arginine Biosynthesis IV | Biosynthesis | +0.74 | Arginine Biosynthesis IV | Biosynthesis | -0.74 |
| 7 | Interconversion of nucleotide di- & triphosphates | Metabolism | -0.73 | Interconversion of nucleotide di- & triphosphates | Metabolism | +0.73 | Interconversion of nucleotide di- & triphosphates | Metabolism | +0.73 | Interconversion of nucleotide di- & triphosphates | Metabolism | +0.74 |
| 8 | Nucleotide salvage | Metabolism, Biosynthesis | -0.73 | Nucleotide salvage | Metabolism, Biosynthesis | -0.73 | Nucleotide salvage | Metabolism, Biosynthesis | +0.73 | Citrulline-Nitric Oxide Cycle | Biosynthesis, Degradation/Utilization/Assimilation | -0.73 |
| 9 | Citrulline-Nitric Oxide Cycle | Biosynthesis, Degradation/Utilization/Assimilation | -0.73 | Citrulline-Nitric Oxide Cycle | Biosynthesis, Degradation/Utilization/Assimilation | -0.73 | Citrulline-Nitric Oxide Cycle | Biosynthesis, Degradation/Utilization/Assimilation | +0.73 | Sirtuin Signaling Pathway | Intracellular and Secondary Messenger Signaling, Transcriptional Regulation | -0.73 |
| 10 | Mismatch Repair | Autophagy, DNA Repair | -0.73 | Sirtuin Signaling Pathway | Intracellular and Secondary Messenger Signaling, Transcriptional Regulation | -0.72 | Mismatch Repair | Autophagy, DNA Repair | +0.73 | Mismatch Repair | Autophagy, DNA Repair | -0.73 |
| 11 | Sirtuin Signaling Pathway | Intracellular & Secondary Messenger Signaling, Transcriptional Regulation | -0.72 | Sulfur amino acid metabolism | Metabolism | +0.72 | Sirtuin Signaling Pathway | Intracellular & Secondary Messenger Signaling, Transcriptional Regulation | -0.72 | Sulfur amino acid metabolism | Metabolism | +0.72 |
| 12 | Sulfur amino acid metabolism | Metabolism | -0.72 | Metabolism of water-soluble vitamins & cofactors | Metabolism | +0.72 | Sulfur amino acid metabolism | Metabolism | +0.72 | Metabolism of water-soluble vitamins & cofactors | Metabolism | +0.72 |
| 13 | Metabolism of water-soluble vitamins & cofactors | Metabolism | -0.72 | Choline catabolism | Metabolism | +0.71 | Metabolism of water-soluble vitamins & cofactors | Metabolism | +0.72 | Choline catabolism | Metabolism | +0.72 |
| 14 | Choline catabolism | Metabolism | +0.71 | Purine Nucleotides De Novo Biosynthesis II | Biosynthesis | -0.69 | Choline catabolism | Metabolism | -0.71 | Urea Cycle | Metabolism, Degradation/Utilization/Assimilation | +0.69 |
| 15 | Purine Nucleotides De Novo Biosynthesis II | Biosynthesis | -0.69 | Glutamate and glutamine metabolism | Metabolism | +0.68 | Purine Nucleotides De Novo Biosynthesis II | Biosynthesis | +0.69 | Purine Nucleotides De Novo Biosynthesis II | Biosynthesis | -0.69 |
| 16 | Glutamate & glutamine metabolism | Metabolism | -0.68 | Nucleotide biosynthesis | Metabolism | +0.68 | Nucleotide biosynthesis | Metabolism | +0.68 | Glutamate & glutamine metabolism | Metabolism | +0.68 |
| 17 | Nucleotide biosynthesis | Metabolism | -0.68 | Pyruvate metabolism | Metabolism | +0.67 | Pyruvate metabolism | Metabolism | +0.67 | Nucleotide biosynthesis | Metabolism | +0.68 |
| 18 | Pyruvate metabolism | Metabolism | -0.67 | Superpathway of Methionine Degradation | Degradation/Utilization/Assimilation | -0.67 | Superpathway of Methionine Degradation | Degradation/Utilization/Assimilation | +0.67 | Pyruvate metabolism | Metabolism | +0.67 |
| 19 | Superpathway of Methionine Degradation | Degradation/Utilization/Assimilation | -0.67 | Flavin Biosynthesis IV (Mammalian) | Biosynthesis | -0.67 | Flavin Biosynthesis IV (Mammalian) | Biosynthesis | +0.67 | Superpathway of Methionine Degradation | Degradation/Utilization/Assimilation | -0.67 |
| 20 | Flavin Biosynthesis IV (Mammalian) | Biosynthesis | -0.67 | Gluconeogenesis I | Biosynthesis | -0.67 | Cytosolic sensors of pathogen-associated DNA | Immune System | +0.67 | Flavin Biosynthesis IV (Mammalian) | Biosynthesis | -0.67 |

**Supplemental Table 2. Top 20 canonical pathways impacted in cerebellar tissue across four auxiliary group comparisons.** This tables lists the top 20 canonical pathways as ranked by Pathway Impact score in cerebellar tissue for the following four group comparisons: Control (+) v. Control (-), Sepsis (+) v. Control (-), Sepsis (-) v. Control (+), and Sepsis (+) v. Control (+). For each pathway, the table includes the associated biological function category and calculated Pathway Impact score. Direction of regulation is denoted by sign on impact score: positive (+) for predicted up-regulation, negative (-) for predicted down-regulation.

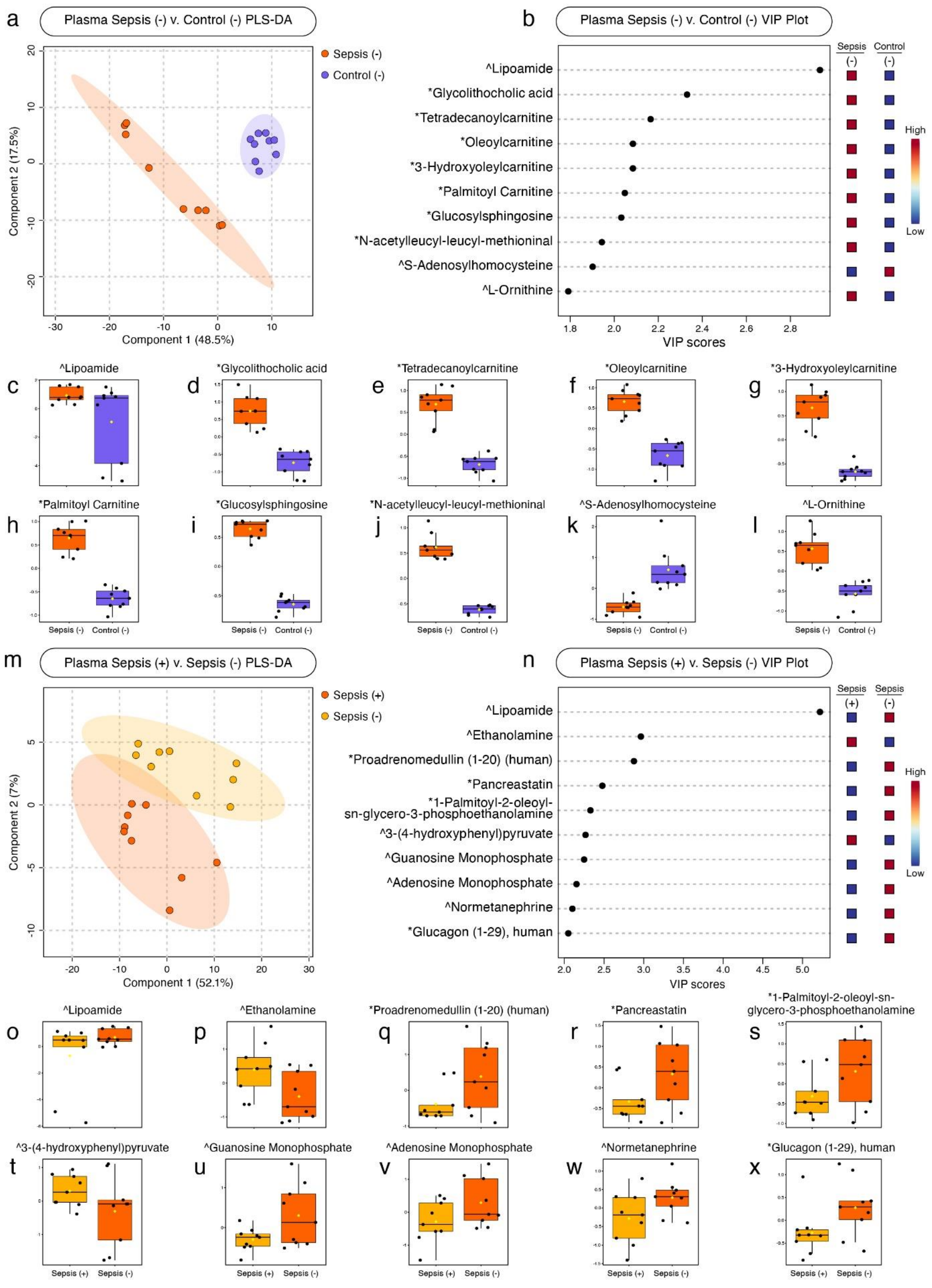

**Supplemental Figure 11. Discriminant metabolite profiles from plasma samples in both disease state and treatment conditions.** Panels **a-l** show results from the disease state comparison, Sepsis (-) v. Control (-), and panel **m-x** display results from the treatment comparsion, Sepsis (+) v. Sepsis (-). (**a, m**) PLS-DA plots displaying separation of experimental groups along the first two model components. (**b, n**) Top 10 metabolites contributing to separation by group assignment, as determined by VIP score. (**c-l, o-x**) Box plots of the top 10 metabolites in disease state and treatment comparisons, respectively, displaying the average relative concentrations of each metabolite by group. Caret symbols(^*)* denote a Level 1-2 confidence identification from core facility panel confirmed by reference standard; Asterisks (*) denote a putatively annotated feature from LC-MS/MS fragmentation matches using the NIST MS Search software, earning Level 3+ identification confidence[25].

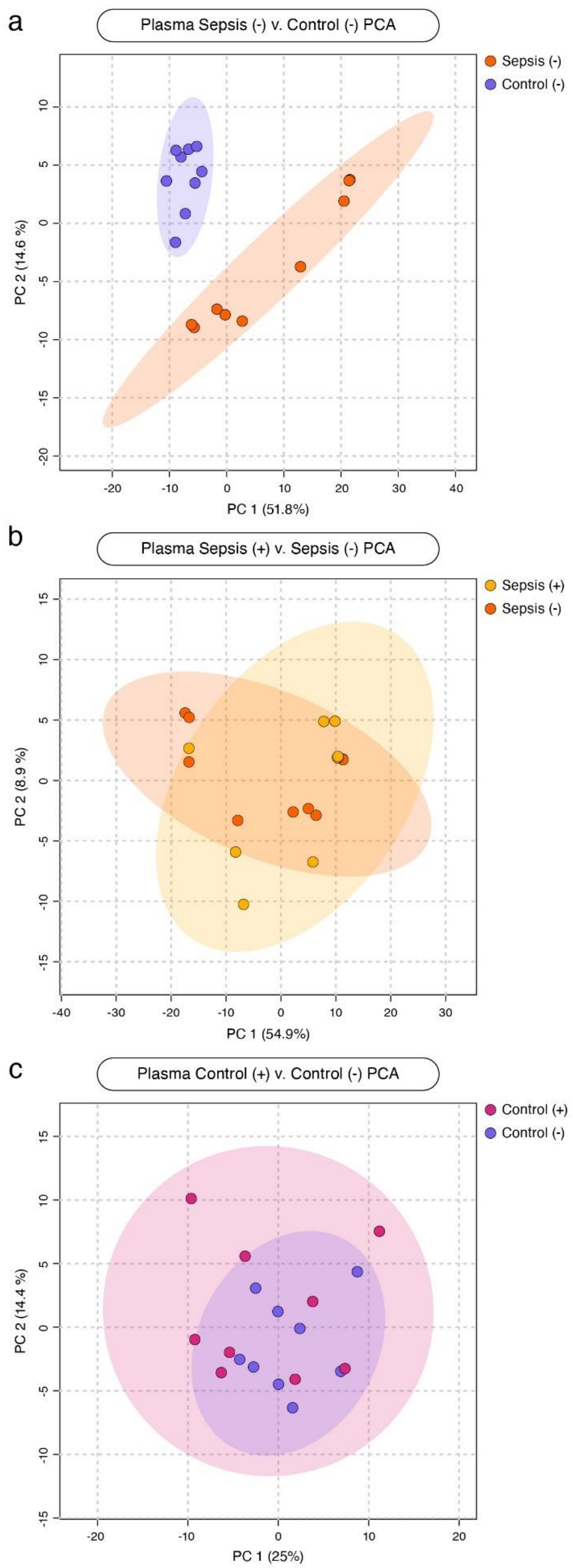

**Supplemental Figure 12. Principal component analyses (PCA) of metabolite profiles from plasma between key pairwise comparisons of disease, treatment, and baseline MSC-sEV conditions.** (**a**) PCA plot of the disease state comparison, Sepsis (-) v. Control (-), in cerebellar tissue samples showing the distribution of individual samples across the first two principal components (PC1 and PC2) as determined by percent of variance explained. (**b**) PCA plot resulting from the treatment comparison, Sepsis (+) v. Sepsis (-). (**c**) PCA plot resulting from the baseline MSC-sEV comparison, Control (+) v. Control (-). Each point represents one sample, colored corresponding to its experimental condition. Shaded ellipses represent a 95% confidence interval for each cluster. Axes represent the percentage of variance explain by PC1 (x-axis) and PC2 (y-axis).

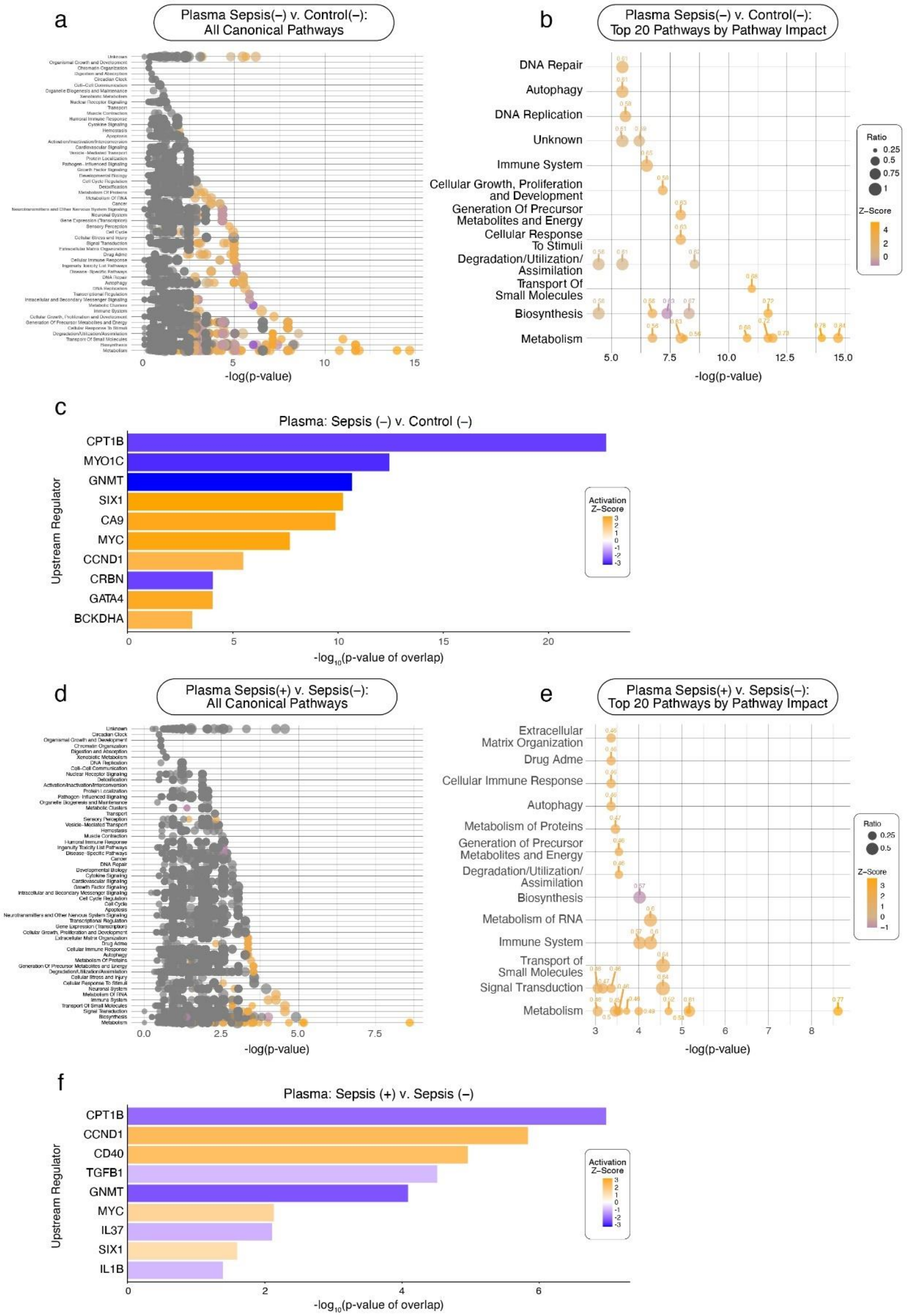

**Supplemental Figure 13**. **Canonical pathway activity and upstream regulator predictions in plasma across disease and treatment comparisons**. Panels **a-c** focus on the disease state comparison, Sepsis (-) v. Control (-), while panels **d-f** focus on the treatment comparison, Sepsis (+) v. Sepsis (-). (**a,d**) Canonical pathway activity showing all pathways predicted to be affected as determined by cerebellar metabolite intensities. Pathways are grouped by their biological function (y-axis) and plotted against their alteration significance (-$\log_{10}$[p-value]; x-axis). Bubble size reflects the proportion of the measured metabolites relative to total known pathway metabolite profile, while bubble color represents the predicted activation direction and intensity (orange = predicted up-regulation, blue = predicted down-regulation). (**b,e**) Top 20 canonical pathways as ranked by Pathway Impact score. (**c,f**) Top 10 predicted upstream regulators influenced by each condition based on metabolite representation. Each regulator (y-axis) is displayed as a bar with a length representing its -$\log_{10}$(p-value of overlap; x-axis) and is colored according to its predicted activation direction and intensity.

| | Plasma | | | | | |
|---|---|---|---|---|---|---|
| | Sepsis (-) v. Control (-) | | | Sepsis (+) v. Sepsis (-) | | |
| Rank | Pathway Name | Biological Function Category | Impact Score | Pathway Name | Biological Function Category | Impact Score |
| 1 | Interconversion of nucleotide di- and triphosphates | Metabolism | +0.84 | Metabolism of water-soluble vitamins and cofactors | Metabolism | +0.77 |
| 2 | Nucleotide catabolism | Metabolism | +0.78 | Vasopressin regulates renal water homeostasis via Aquaporins | Transport of small molecules | +0.64 |
| 3 | Sulfur amino acid metabolism | Metabolism | +0.73 | GPER1 signaling | Signal Transduction | +0.64 |
| 4 | Nucleotide salvage | Metabolism | +0.72 | Metabolism of polyamines | Metabolism | +0.61 |
| 5 | Metabolism of water-soluble vitamins and cofactors | Metabolism | +0.68 | Rap1 signalling | Immune System | +0.60 |
| 6 | Transport of bile salts and organic acids, metal ions and amine compounds | Transport of small molecules | +0.68 | rRNA modification in the nucleus and cytosol | Metabolism of RNA | +0.60 |
| 7 | Pyrimidine Deoxyribonucleotides De Novo Biosynthesis I | Biosynthesis | -0.67 | Cytosolic sensors of pathogen-associated DNA | Immune System | +0.57 |
| 8 | Cytosolic sensors of pathogen-associated DNA | Immune System | +0.65 | Biotin-carboxyl Carrier Protein Assembly | Biosynthesis | -0.57 |
| 9 | Pentose phosphate pathway | Generation of Precursor Metabolites and Energy | +0.63 | Nucleotide catabolism | Metabolism | +0.54 |
| 10 | Salvage Pathways of Pyrimidine Ribonucleotides | Biosynthesis | -0.63 | Sulfur amino acid metabolism | Metabolism | +0.52 |
| 11 | Superpathway of Methionine Degradation | Degradation/Utilization/Assimilation | +0.62 | Creatine metabolism | Metabolism | +0.50 |
| 12 | Mismatch Repair | DNA Repair | +0.61 | Metabolism of cofactors | Metabolism | +0.49 |
| 13 | Ascorbate Recycling (Cytosolic) | Degradation/Utilization/Assimilation | +0.61 | Gamma carboxylation, hypusinylation, hydroxylation, and arylsulfatase activation | Metabolism of Proteins | +0.47 |
| 14 | Methionine Degradation I (to Homocysteine) | Degradation/Utilization/Assimilation | +0.61 | G alpha (z) signalling events | Signal Transduction | +0.47 |
| 15 | Cysteine Biosynthesis III (mammalia) | Biosynthesis | +0.59 | Branched-chain amino acid catabolism | Metabolism | +0.46 |
| 16 | Folate Signaling Pathway | Cellular Growth, Proliferation and Development | +0.58 | Extra-nuclear estrogen signaling | Signal Transduction | +0.46 |
| 17 | Synthesis of DNA | DNA Replication | +0.58 | Interconversion of nucleotide di- and triphosphates | Metabolism | +0.46 |
| 18 | Glycine Degradation (Creatine Biosynthesis) | Biosynthesis | +0.56 | Ketone body metabolism | Metabolism | +0.46 |
| 19 | Phase II - Conjugation of compounds | Metabolism | +0.56 | RAS processing | Signal Transduction | +0.46 |
| 20 | Nucleotide biosynthesis | Metabolism | +0.56 | Peroxisomal lipid metabolism | Metabolism | +0.45 |

**Supplemental Table 3. Top 20 canonical pathways impacted in plasma samples from primary disease state and treatment comparisons.** This table lists the top 20 canonical pathways ranked by Pathway Impact score as determined by plasma sample results of those two primary comparisons, Sepsis (-) v. Control (-) for disease state and Sepsis (+) v. Sepsis (-) for treatment. Each pathway is listed with its corresponding biological function category and Pathway Impact score. Direction of regulation is denoted by sign on impact score: positive (+) for predicted up-regulation, negative (-) for predicted down-regulation.

| | Plasma | | | | | | | | | | | |
|---|---|---|---|---|---|---|---|---|---|---|---|---|
| | Control (+) v. Control (-) | | | Sepsis (+) v. Control (-) | | | Sepsis (-) v. Control (+) | | | Sepsis (+) v. Control (+) | | |
| Rank | Pathway | Biological Function Category | Pathway Impact | Pathway | Biological Function Category | Pathway Impact | Pathway | Biological Function Category | Pathway Impact | Pathway | Biological Function Category | Pathway Impact |
| 1 | Transport of bile salts and organic acids, metal ions and amine compounds | Transport Of Small Molecules | +0.83 | Nucleotide catabolism | Metabolism | +0.80 | Interconversion of nucleotide di- and triphosphates | Metabolism | +0.83 | Nucleotide catabolism | Metabolism | +0.78 |
| 2 | Interconversion of nucleotide di- and triphosphates | Metabolism | +0.81 | Interconversion of nucleotide di- and triphosphates | Metabolism | +0.79 | Metabolism of water-soluble vitamins and cofactors | Metabolism | +0.70 | Interconversion of nucleotide di- and triphosphates | Metabolism | +0.73 |
| 3 | Nucleotide catabolism | Metabolism | +0.78 | Nucleotide salvage | Metabolism | +0.72 | Nucleotide catabolism | Metabolism | +0.69 | Nucleotide salvage | Metabolism | +0.65 |
| 4 | Superpathway of Citrulline Metabolism | Biosynthesis | +0.73 | Pyrimidine Deoxyribonucleotides De Novo Biosynthesis I | Biosynthesis | -0.69 | Sulfur amino acid metabolism | Metabolism | +0.67 | Salvage Pathways of Pyrimidine Ribonucleotides | Biosynthesis | -0.65 |
| 5 | Synthesis of DNA | DNA Replication | +0.70 | Cytosolic sensors of pathogen-associated DNA | Immune System | +0.67 | Ascorbate Recycling (Cytosolic) | Degradation/ Utilization/ Assimilation | -0.64 | Transport of bile salts and organic acids, metal ions and amine compounds | Transport Of Small Molecules | +0.59 |
| 6 | tRNA Charging | Biosynthesis | -0.70 | Sulfur amino acid metabolism | Metabolism | +0.65 | Transport of bile salts and organic acids, metal ions and amine compounds | Transport Of Small Molecules | +0.60 | Metabolism of water-soluble vitamins and cofactors | Metabolism | +0.58 |
| 7 | Arginine Biosynthesis IV | Biosynthesis | +0.69 | Transport of bile salts and organic acids, metal ions and amine compounds | Transport Of Small Molecules | +0.65 | Salvage Pathways of Pyrimidine Ribonucleotides | Biosynthesis | -0.60 | Mismatch Repair | Autophagy | +0.58 |
| 8 | Citrulline Biosynthesis | Biosynthesis | -0.69 | Salvage Pathways of Pyrimidine Ribonucleotides | Biosynthesis | -0.64 | Cytosolic sensors of pathogen-associated DNA | Immune System | +0.60 | Pentose phosphate pathway | Cellular Response To Stimuli | +0.55 |
| 9 | Transport of inorganic cations/anions and amino acids/oligopeptides | Transport Of Small Molecules | +0.67 | Metabolism of water-soluble vitamins and cofactors | Metabolism | +0.64 | Nucleotide salvage | Metabolism | +0.59 | Cytosolic sensors of pathogen-associated DNA | Immune System | +0.54 |
| 10 | Mismatch Repair | Autophagy | +0.67 | Ascorbate Recycling (Cytosolic) | Degradation/ Utilization/ Assimilation | +0.62 | Superpathway of Methionine Degradation | Degradation/Utilization/Assimilation | +0.58 | Polyamine Regulation in Colon Cancer | Cancer | +0.54 |
| 11 | Urea Cycle | Metabolism | +0.67 | Pentose phosphate pathway | Cellular Response To Stimuli | +0.59 | Tryptophan catabolism | Metabolism | +0.57 | Transport of vitamins, nucleosides, and related molecules | Transport Of Small Molecules | +0.53 |
| 12 | Nucleotide salvage | Metabolism | +0.67 | Choline catabolism | Metabolism | +0.54 | Folate Signaling Pathway | Cellular Growth, Proliferation & Development | +0.57 | Metabolism of polyamines | Metabolism | +0.51 |
| 13 | Metabolism of water-soluble vitamins and cofactors | Metabolism | +0.66 | Mismatch Repair | Autophagy | +0.53 | Pentose phosphate pathway | Cellular Response To Stimuli | +0.56 | Pyrimidine Deoxyribonucleotides De Novo Biosynthesis I | Biosynthesis | 0.50 |
| 14 | Nucleotide biosynthesis | Metabolism | +0.66 | Phase II - Conjugation of compounds | Metabolism | +0.53 | Phase II - Conjugation of compounds | Metabolism | +0.55 | Sulfur amino acid metabolism | Metabolism | +0.49 |
| 15 | Glucose metabolism | Metabolism | +0.65 | Nucleotide biosynthesis | Biosynthesis | +0.53 | Metabolism of cofactors | Metabolism | +0.55 | Synthesis of DNA | DNA Replication | +0.49 |
| 16 | Sulfur amino acid metabolism | Metabolism | +0.64 | Uridine-5'-phosphate Biosynthesis | Biosynthesis | -0.53 | Mismatch Repair | Autophagy | +0.55 | Superpathway of Methionine Degradation | Degradation/ Utilization/ Assimilation | +0.49 |
| 17 | Pyruvate metabolism | Metabolism | +0.63 | Atorvastatin ADME | Drug Adme | +0.52 | Metabolism of polyamines | Metabolism | +0.54 | Integration of energy metabolism | Metabolism | +0.48 |
| 18 | Gap junction trafficking and regulation | Vesicle-Mediated Transport | +0.63 | Methionine Salvage II (Mammalian) | Biosynthesis | -0.52 | Glyoxylate metabolism and glycine degradation | Metabolism | +0.54 | Glucose metabolism | Metabolism | +0.48 |
| 19 | Signaling by Type 1 Insulin-like Growth Factor 1 Receptor (IGF1R) | Signal Transduction | +0.63 | Transport of vitamins, nucleosides, and related molecules | Biosynthesis | +0.52 | Atorvastatin ADME | Drug Adme | +0.53 | Tryptophan catabolism | Metabolism | +0.47 |
| 20 | RAB GEFs exchange GTP for GDP on RABs | Vesicle-Mediated Transport | +0.63 | Gluconeogenesis I | Biosynthesis | -0.52 | Methionine Salvage II (Mammalian) | Biosynthesis | -0.53 | Metabolism of cofactors | Metabolism | +0.46 |

**Supplemental Table 4. Top 20 canonical pathways impacted in plasma across four auxiliary group comparisons.** This tables lists the top 20 canonical pathways as ranked by Pathway Impact score in cerebellar tissue for the following four group comparisons: Control (+) v. Control (-), Sepsis (+) v. Control (-), Sepsis (-) v. Control (+), and Sepsis (+) v. Control (+). For each pathway, the table includes the associated biological function category and calculated Pathway Impact score. Direction of regulation is denoted by sign on impact score: positive (+) for predicted up-regulation, negative (-) for predicted down-regulation.

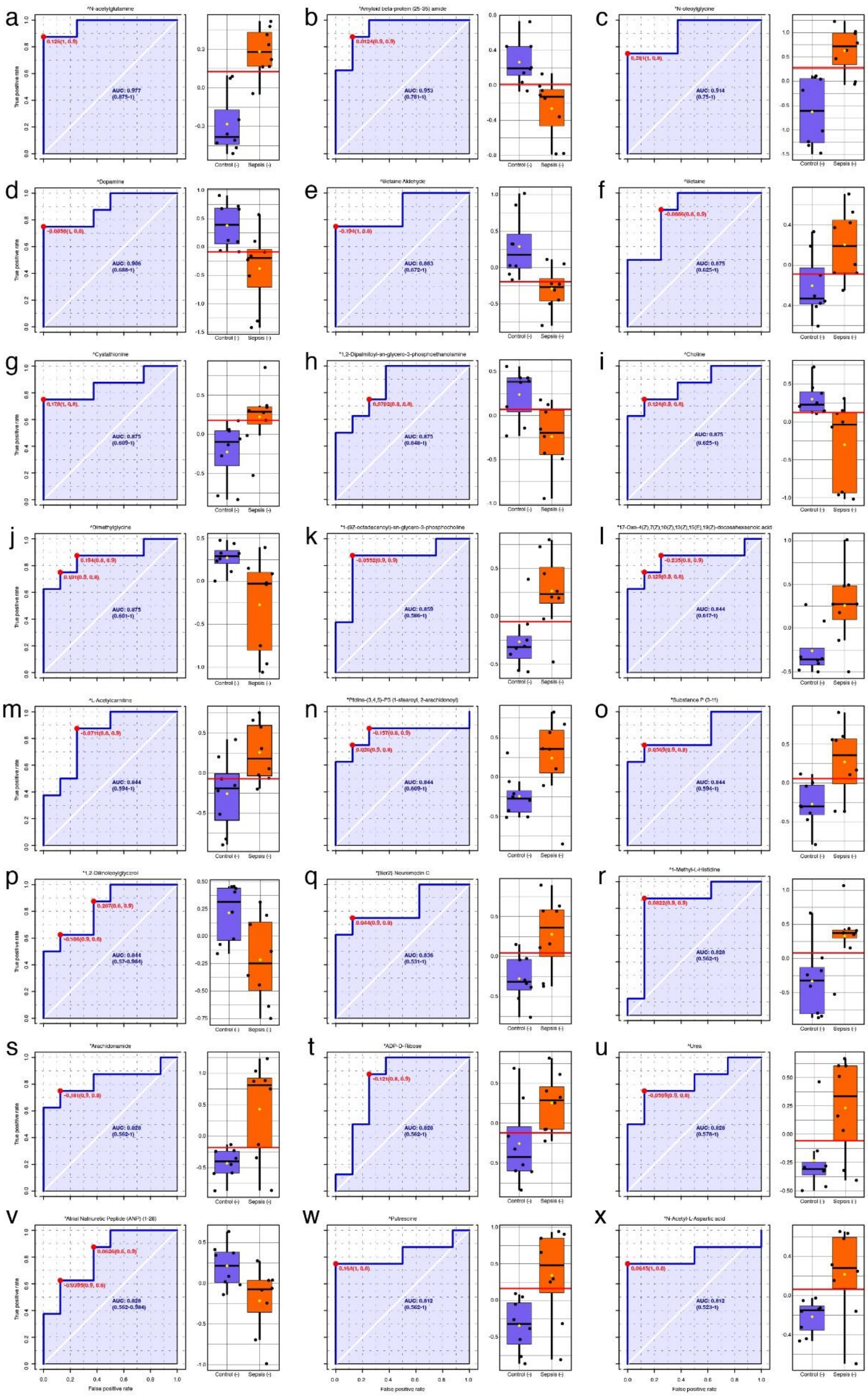

**Supplemental Figure 14. Top biomarker candidates identified in cerebellar tissue with strong diagnostic criteria.** This figure displays 24 metabolite features with strong diagnostic potential (AUC ≥ 0.8) for identifying Sepsis (-) animals from Control (-) animals. Panels **a-x** display receiver operating characteristic (ROC) curves for each overlapping metabolite candidate, with AUC values shown to reflect classification performance. Box plots are displayed to the right of each metabolite's ROC curve, illustrating the relative average concentrations of each metabolite by condition. Where known, the best diagnostic threshold is displayed by a red horizontal line on the box plots. Caret symbols (^) denote a Level 1-2 confidence identification from core facility panel confirmed by reference standard; Asterisks (*) denote a putatively annotated feature from LC-MS/MS fragmentation matches using the NIST MS Search software, earning Level 3+ identification confidence[25].

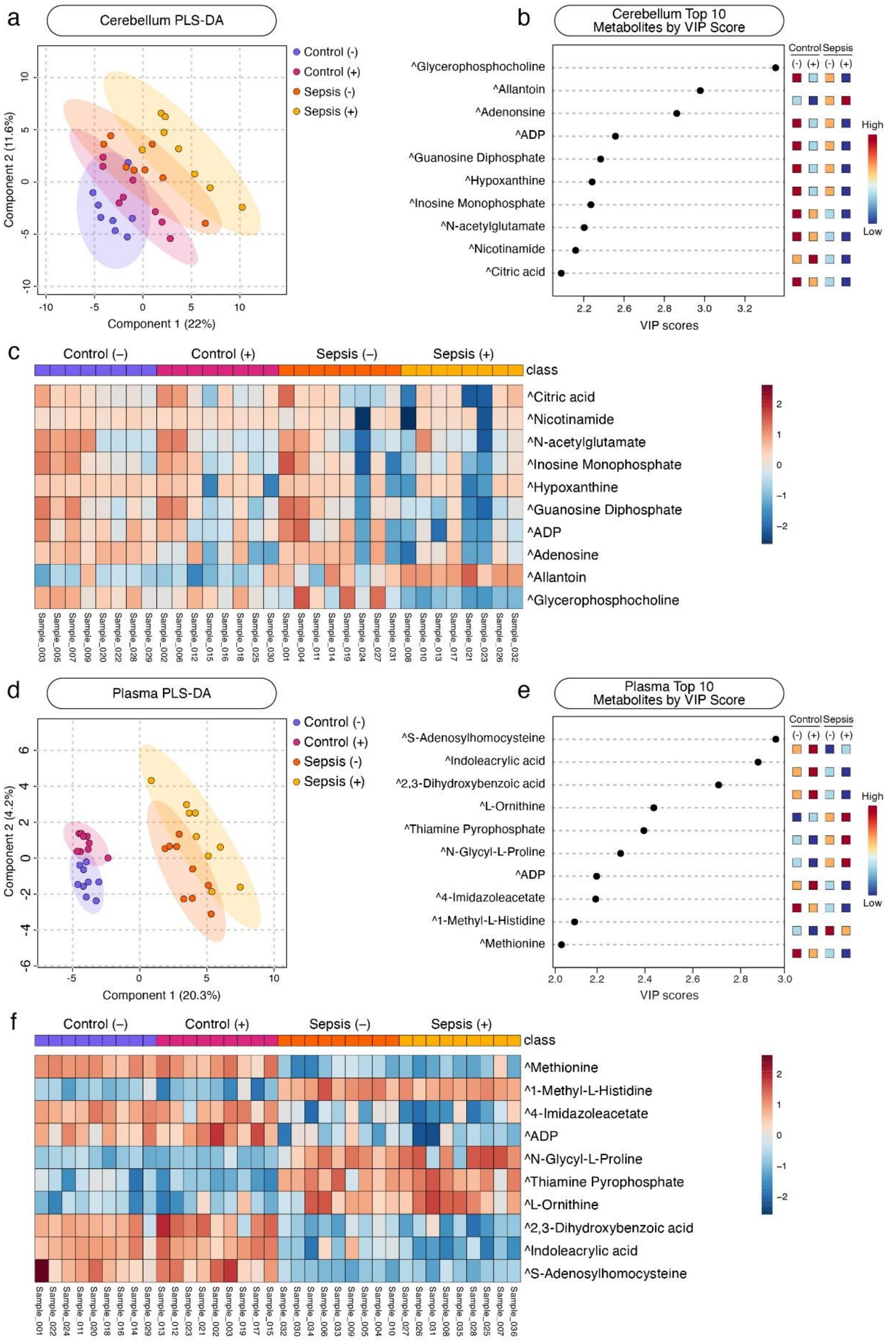

**Supplemental Figure 15. Targeted metabolomics-based multivariate analysis identifies discriminatory targeted metabolite profiles in cerebellar tissue and plasma.** PLS-DA plots of cerebellum (**a**) and plasma (**d**) samples where the horizontal and vertical axes represent Components 1 and 2, respectively. These components capture directions in the data that maximize covariance between metabolite profiles and group labels where Component 1 explains the strongest class-related pattern and Component 2 the next most discriminative, orthogonal pattern. Ellipses represent the 95% confidence interval. b) Variable Importance in Projection (VIP) score graphs for cerebellum (**b**) and plasma (**e**) samples display the top 10 metabolites contributing to group separation along Component 1, with VIP scores reflecting each metabolite's contribution to the model's power to discriminate samples by group. Heatmaps of the top 10 metabolites by VIP score from cerebellum (**c**) and plasma (**f**) PLS-DAs showing relative peak intensities across each sample and grouped by experimental condition. Caret symbols (^) denote a Level 1-2 confidence identification from core facility panel confirmed by reference standard[25].

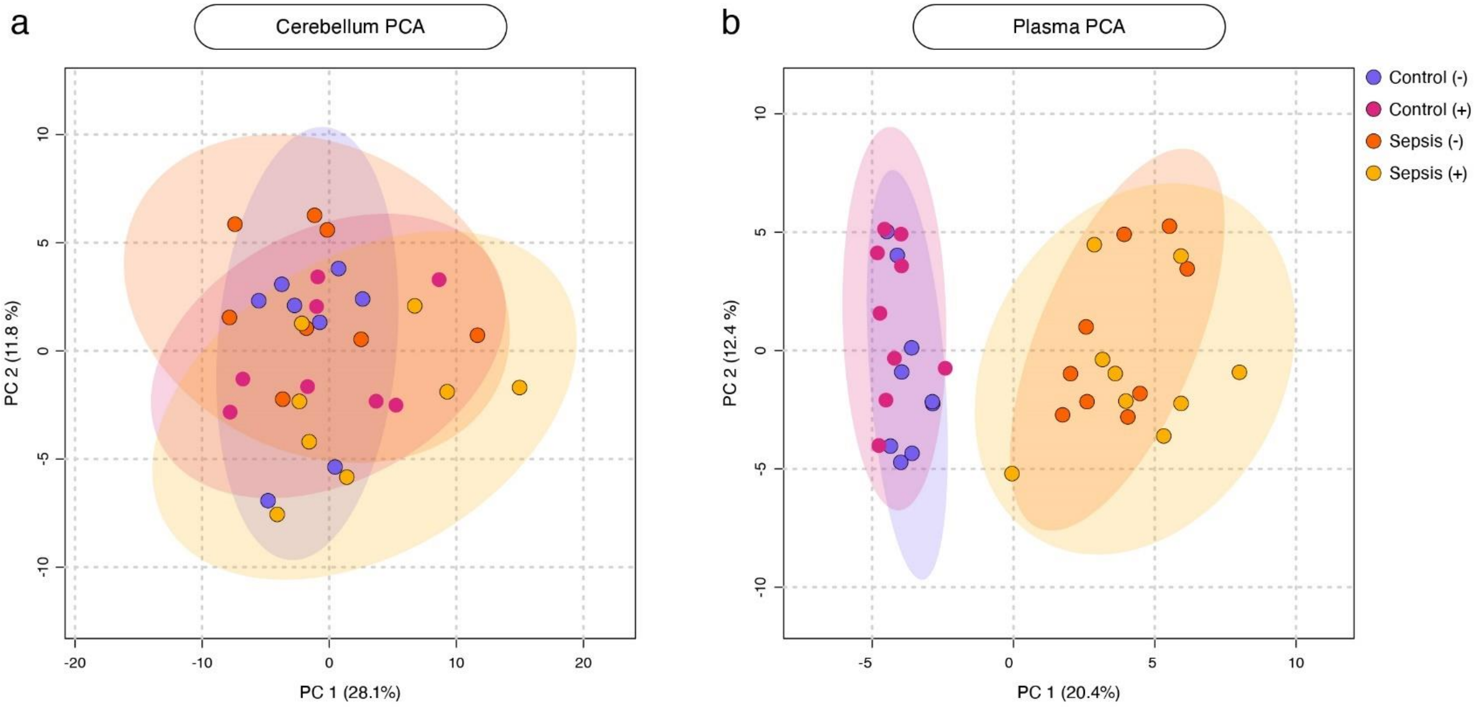

**Supplemental Figure 16. Principal component analyses (PCAs) of cerebellar tissue and plasma targeted-only metabolite profiles between all experimental groups.** (**a**) PCA plot of cerebellar tissue samples showing the distribution of individual samples across the first two principal components (PC1 and PC2) as determined by percent of variance explained. (**b**) PCA plot of resulting from plasma samples. Each point represents one sample, colored corresponding to its experimental condition. Shaded ellipses represent a 95% confidence interval for each cluster. Axes represent the percentage of variance explain by PC1 (x-axis) and PC2 (y-axis).

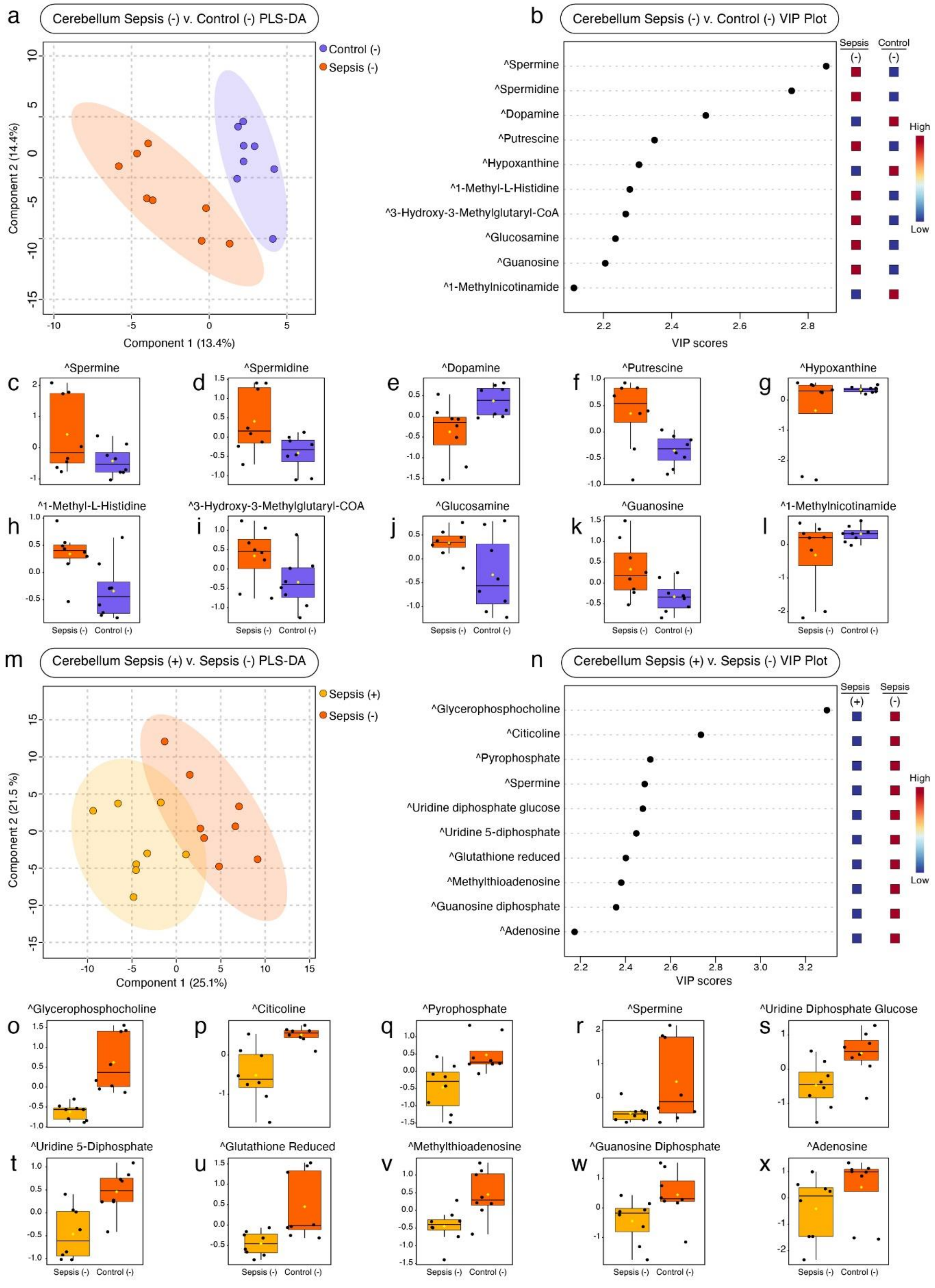

**Supplemental Figure 17. Targeted metabolomics reveal most influential metabolites driving disease state and treatment-dependent group differentiation in cerebellum.** This figure presents PLS-DA-derived discriminant features from cerebellar samples, restricted to two pairwise group comparisons. Panels **a-l** illustrate the comparison between untreated septic animals and healthy control, Sepsis (-) v. Control (-), isolating the metabolic profiles associated with disease state. Panels **m-x** display data from the Sepsis (+) v. Sepsis (-) comparison, capturing effects related to the effect of MSC-sEV treatment under septic conditions. (**a, m**) PLS-DA model illustration showing sample clustering along the first two components with 95% confidence ellipses for each group. (**b, n**) VIP score plots showing the top 10 metabolites contributing to Component 1 and driving separation by experimental condition. (**c-l, o-x**) Box plots for each of the top 10 metabolites by VIP score in their respective comparisons, depicting average relative peak intensity by group. Caret symbols (^) denote a Level 1-2 confidence identification from core facility panel confirmed by reference standard[25].

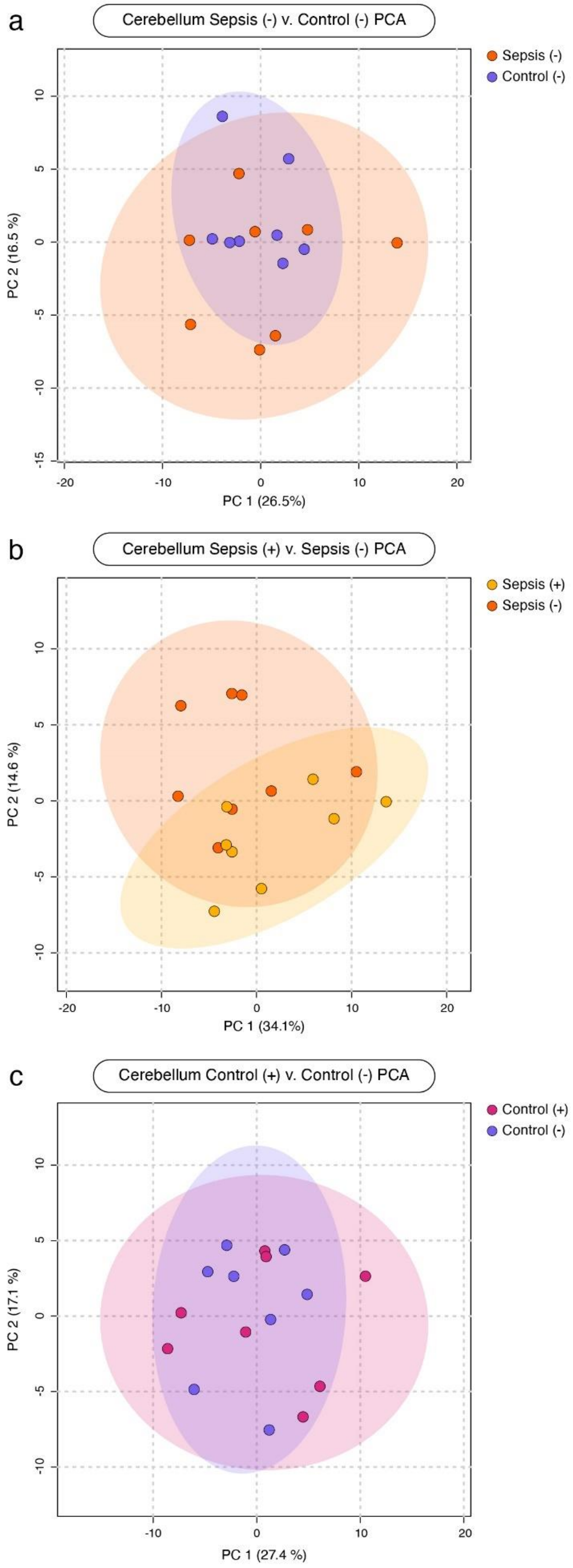

**Supplemental Figure 18. Principal component analyses (PCA) of targeted-only metabolite profiles in cerebellar tissue between key pairwise comparisons of disease, treatment, and baseline MSC-sEV conditions.** (**a**) PCA plot of the disease state comparison, Sepsis (-) v. Control (-), in cerebellar tissue samples showing the distribution of individual samples across the first two principal components (PC1 and PC2) as determined by percent of variance explained. (**b**) PCA plot resulting from the treatment comparison, Sepsis (+) v. Sepsis (-). (**c**) PCA plot resulting from the baseline MSC-sEV comparison, Control (+) v. Control (-). Each point represents one sample, colored corresponding to its experimental condition. Shaded ellipses represent a 95% confidence interval for each cluster. Axes represent the percentage of variance explain by PC1 (x-axis) and PC2 (y-axis).

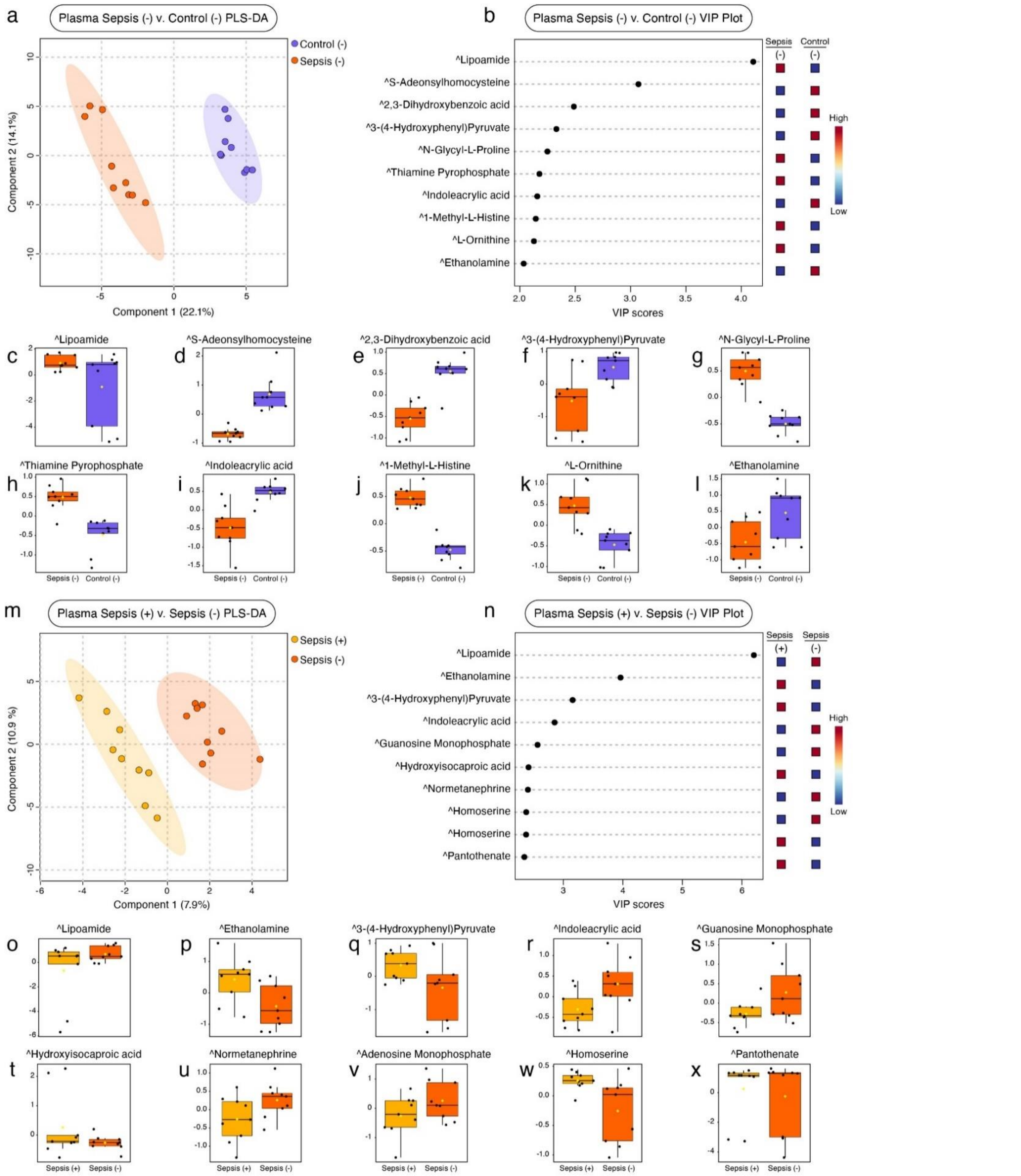

**Supplemental Figure 19. Targeted metabolomics-based discriminant metabolite profiles from plasma samples in both disease state and treatment conditions.** Panels **a-l** show results from the disease state comparison, Sepsis (-) v. Control (-), and panel **m-x** display results from the treatment comparsion, Sepsis (+) v. Sepsis (-). (**a, m**) PLS-DA plots displaying separation of experimental groups along the first two model components. (**b, n**) Top 10 metabolites contributing to separation by group assignment, as determined by VIP score. (**c-l, o-x**) Box plots of the top 10 metabolites in disease state and treatment comparisons, respectively, displaying the average relative concentrations of each metabolite by group. Caret symbols (^) denote a Level 1-2 confidence identification from core facility panel confirmed by reference standard[25].

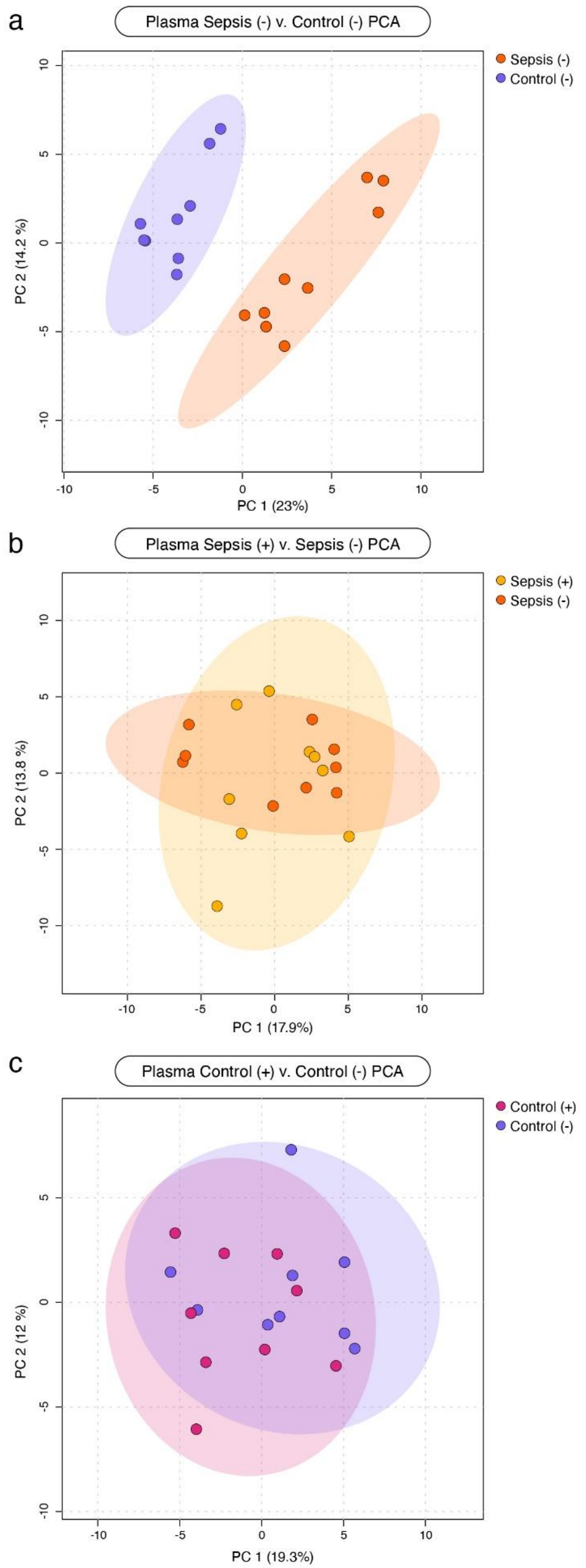

**Supplemental Figure 20. Principal component analyses (PCA) of targeted-only metabolite profiles from plasma between key pairwise comparisons of disease, treatment, and baseline MSC-sEV conditions.** (**a**) PCA plot of the disease state comparison, Sepsis (-) v. Control (-), in cerebellar tissue samples showing the distribution of individual samples across the first two principal components (PC1 and PC2) as determined by percent of variance explained. (**b**) PCA plot resulting from the treatment comparison, Sepsis (+) v. Sepsis (-). (**c**) PCA plot resulting from the baseline MSC-sEV comparison, Control (+) v. Control (-). Each point represents one sample, colored corresponding to its experimental condition. Shaded ellipses represent a 95% confidence interval for each cluster. Axes represent the percentage of variance explain by PC1 (x-axis) and PC2 (y-axis).

| | Targeted Metabolomics Only | | | | | |
|---|---|---|---|---|---|---|
| | Sepsis (-) v. Control (-) | | | Sepsis (+) v. Sepsis (-) | | |
| Rank | Pathway Name | Biological Function Category | Impact Score | Pathway Name | Biological Function Category | Impact Score |
| 1 | Transport of bile salts and organic acids, metal ions and amine compounds | Transport of small molecules | -0.84 | Transport of bile salts and organic acids, metal ions and amine compounds | Transport of small molecules | +0.84 |
| 2 | Superpathway of Citrulline Metabolism | Biosynthesis | +0.80 | Superpathway of Citrulline Metabolism | Biosynthesis | -0.80 |
| 3 | Citric acid cycle (TCA cycle) | Metabolism | -0.79 | Citric acid cycle (TCA cycle) | Metabolism | +0.79 |
| 4 | Metabolism of water-soluble vitamins and cofactors | Metabolism | +0.79 | Metabolism of water-soluble vitamins and cofactors | Metabolism | +0.79 |
| 5 | Urea Cycle | Metabolism | +0.78 | Urea Cycle | Metabolism | -0.78 |
| 6 | Arginine Biosynthesis IV | Biosynthesis | +0.78 | Arginine Biosynthesis IV | Biosynthesis | -0.78 |
| 7 | Sirtuin Signaling Pathway | Intracellular and Secondary Messenger Signaling, Transcriptional Regulation | -0.78 | Sirtuin Signaling Pathway | Intracellular and Secondary Messenger Signaling, Transcriptional Regulation | -0.78 |
| 8 | Nucleotide catabolism | Metabolism | +0.77 | Nucleotide catabolism | Metabolism | +0.77 |
| 9 | Nucleotide salvage | Metabolism | -0.76 | Nucleotide salvage | Metabolism | -0.76 |
| 10 | Citrulline-Nitric Oxide Cycle | Biosynthesis | +0.76 | Citrulline-Nitric Oxide Cycle | Biosynthesis | -0.76 |
| 11 | Choline catabolism | Metabolism | -0.75 | Choline catabolism | Metabolism | +0.75 |
| 12 | Sulfur amino acid metabolism | Metabolism | -0.75 | Sulfur amino acid metabolism | Metabolism | +0.75 |
| 13 | Pyruvate metabolism | Metabolism | -0.72 | Warburg Effect Signaling Pathway | Cancer | +0.72 |
| 14 | Superpathway of Methionine Degradation | Degradation/Utilization/Assimilation | +0.72 | Pyruvate metabolism | Metabolism | +0.72 |
| 15 | Nucleotide biosynthesis | Metabolism | -0.72 | Superpathway of Methionine Degradation | Degradation/Utilization/Assimilation | -0.72 |
| 16 | Transport of inorganic cations/anions and amino acids/oligopeptides | Transport of small molecules | -0.71 | Nucleotide biosynthesis | Metabolism | +0.72 |
| 17 | Glucose metabolism | Metabolism | -0.71 | Transport of inorganic cations/anions and amino acids/oligopeptides | Transport of small molecules | +0.71 |
| 18 | Glyoxylate metabolism and glycine degradation | Metabolism | -0.71 | Glucose metabolism | Metabolism | +0.71 |
| 19 | Gluconeogenesis I | Biosynthesis | +0.70 | Glyoxylate metabolism and glycine degradation | Metabolism | +0.71 |
| 20 | Cytosolic sensors of pathogen-associated DNA | Immune System | +0.69 | Gluconeogenesis I | Biosynthesis | -0.70 |

**Supplemental Table 5. Targeted metabolomics-based top 20 canonical pathways altered in cerebellar tissue by disease and treatment conditions**. Listed in this table are the top 20 canonical pathways altered from between disease state, Sepsis (-) v. Control (-), and treatment, Sepsis (+) v. Sepsis (-), comparisons as determined by Pathway Impact score based only on targeted metabolomics results. This table identifies the individual pathway names, biological function category to which each pathway belongs, and the Pathway Impact score for each pathway. Direction of regulation is denoted by sign on impact score: positive (+) for predicted up-regulation, negative (-) for predicted down-regulation.

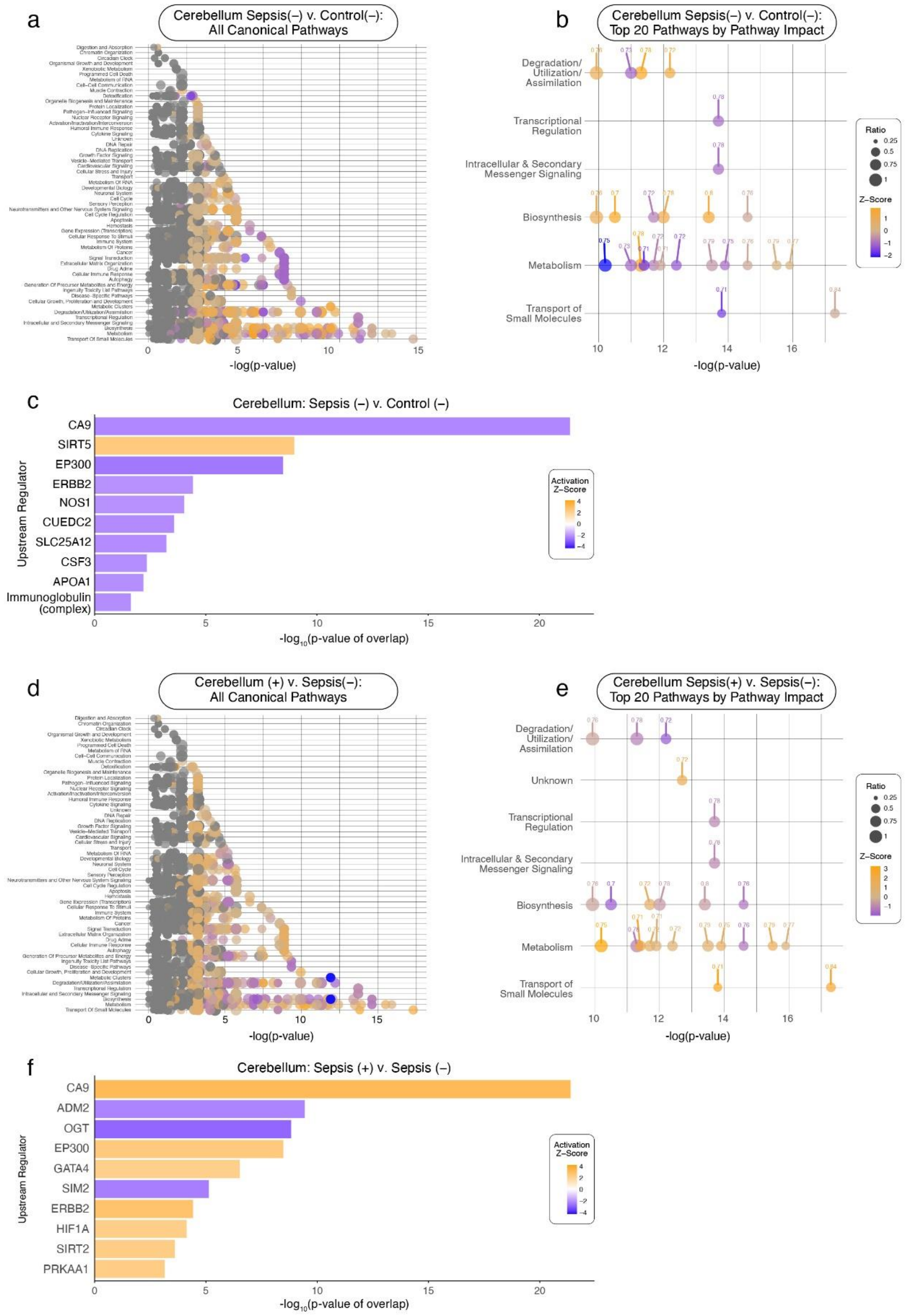

**Supplemental Figure 21. Targeted metabolomics-based canonical pathway activity and upstream regulator predictions in cerebellar tissue across disease and treatment comparisons**. Panels **a-c** focus on the disease state comparison, Sepsis (-) v. Control (-), while panels **d-f** focus on the treatment comparison, Sepsis (+) v. Sepsis (-). (**a,d**) Canonical pathway activity showing all pathways predicted to be affected as determined by cerebellar targeted metabolite intensities. Pathways are grouped by their biological function (y-axis) and plotted against their alteration significance (-$\log_{10}$[p-value]; x-axis). Bubble size reflects the proportion of the measured metabolites relative to total known pathway metabolite profile, while bubble color represents the predicted activation direction and intensity (orange = predicted up-regulation, blue = predicted down-regulation). (**b,e**) Top 20 canonical pathways as ranked by Pathway Impact score. (**c,f**) Top 10 predicted upstream regulators influenced by each condition based on metabolite representation. Each regulator (y-axis) is displayed as a bar with a length representing its -$\log_{10}$(p-value of overlap; x-axis) and is colored according to its predicted activation direction and intensity.

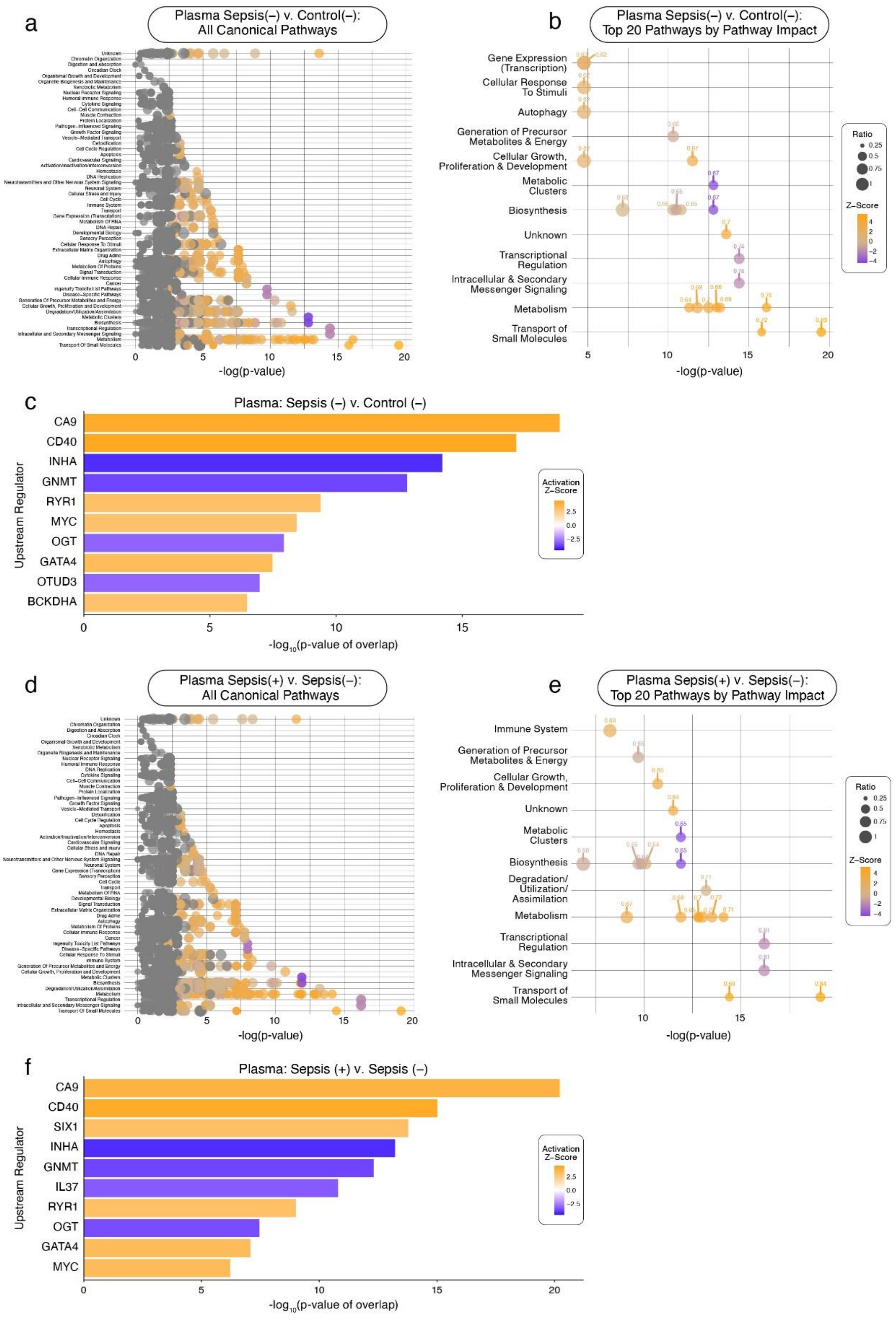

**Supplemental Figure 22. Targeted metabolomics-based canonical pathway activity and upstream regulator predictions in plasma across disease and treatment comparisons**. Panels **a-c** focus on the disease state comparison, Sepsis (-) v. Control (-), while panels **d-f** focus on the treatment comparison, Sepsis (+) v. Sepsis (-). (**a,d**) Canonical pathway activity showing all pathways predicted to be affected as determined by plasma targeted metabolite intensities. Pathways are grouped by their biological function (y-axis) and plotted against their alteration significance (-$\log_{10}$[p-value]; x-axis). Bubble size reflects the proportion of the measured metabolites relative to total known pathway metabolite profile, while bubble color represents the predicted activation direction and intensity (orange = predicted up-regulation, blue = predicted down-regulation). (**b,e**) Top 20 canonical pathways as ranked by Pathway Impact score. (**c,f**) Top 10 predicted upstream regulators influenced by each condition based on metabolite representation. Each regulator (y-axis) is displayed as a bar with a length representing its -$\log_{10}$(p-value of overlap; x-axis) and is colored according to its predicted activation direction and intensity.

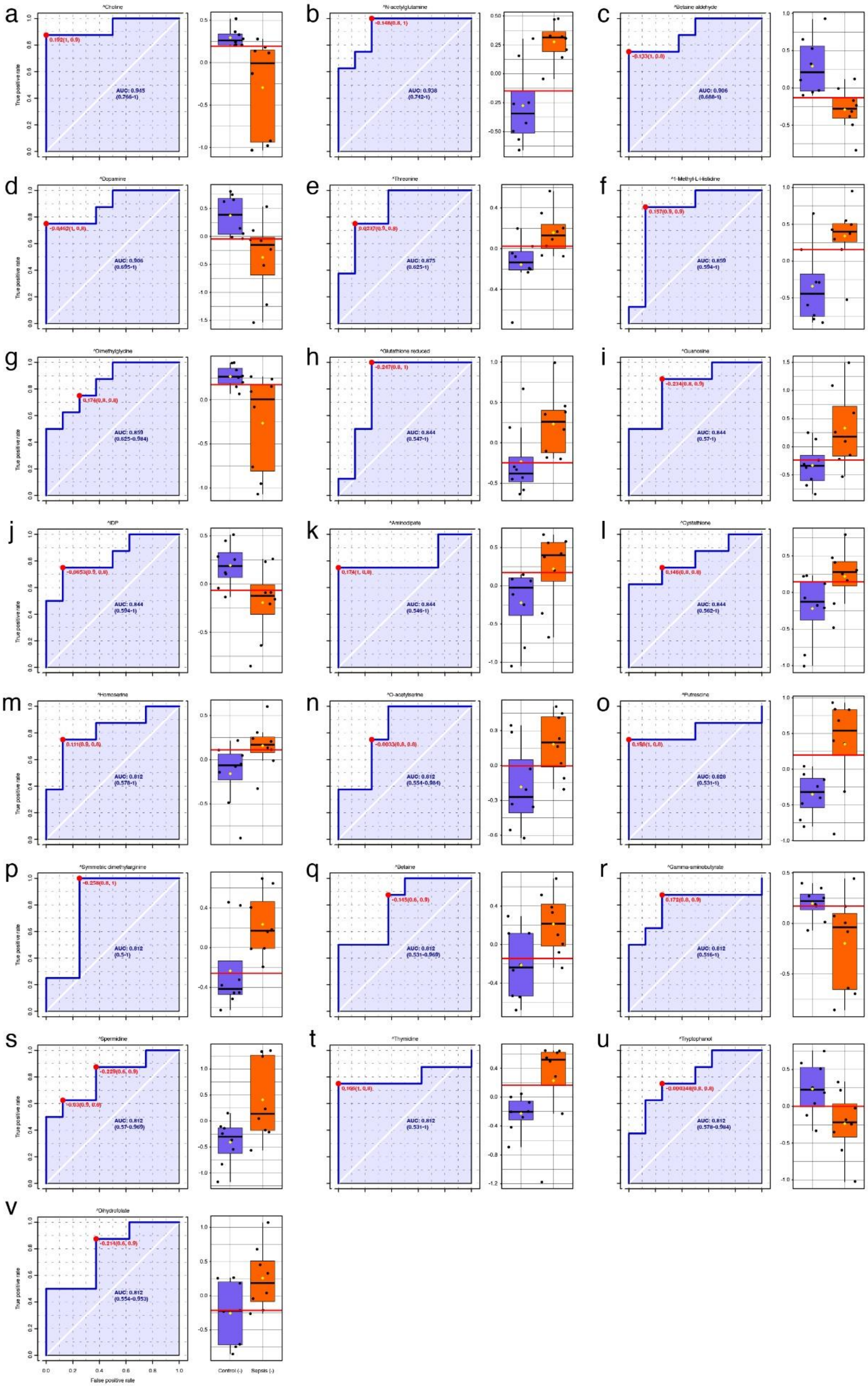

**Supplemental Figure 23. Targeted metabolomics-based top biomarker candidates identified in cerebellar tissue with strong diagnostic criteria.** This figure displays 22 targeted metabolite features with strong diagnostic potential (AUC $\geq$ 0.8) for identifying Sepsis (-) animals from Control (-) animals. Panels **a-x** display receiver operating characteristic (ROC) curves for each overlapping metabolite candidate, with AUC values shown to reflect classification performance. Box plots are displayed to the right of each metabolite's ROC curve, illustrating the relative average concentrations of each metabolite by condition. Where known, the best diagnostic threshold is displayed by a red horizontal line on the box plots. Caret symbols (^) denote a Level 1-2 confidence identification from core facility panel confirmed by reference standard[25].

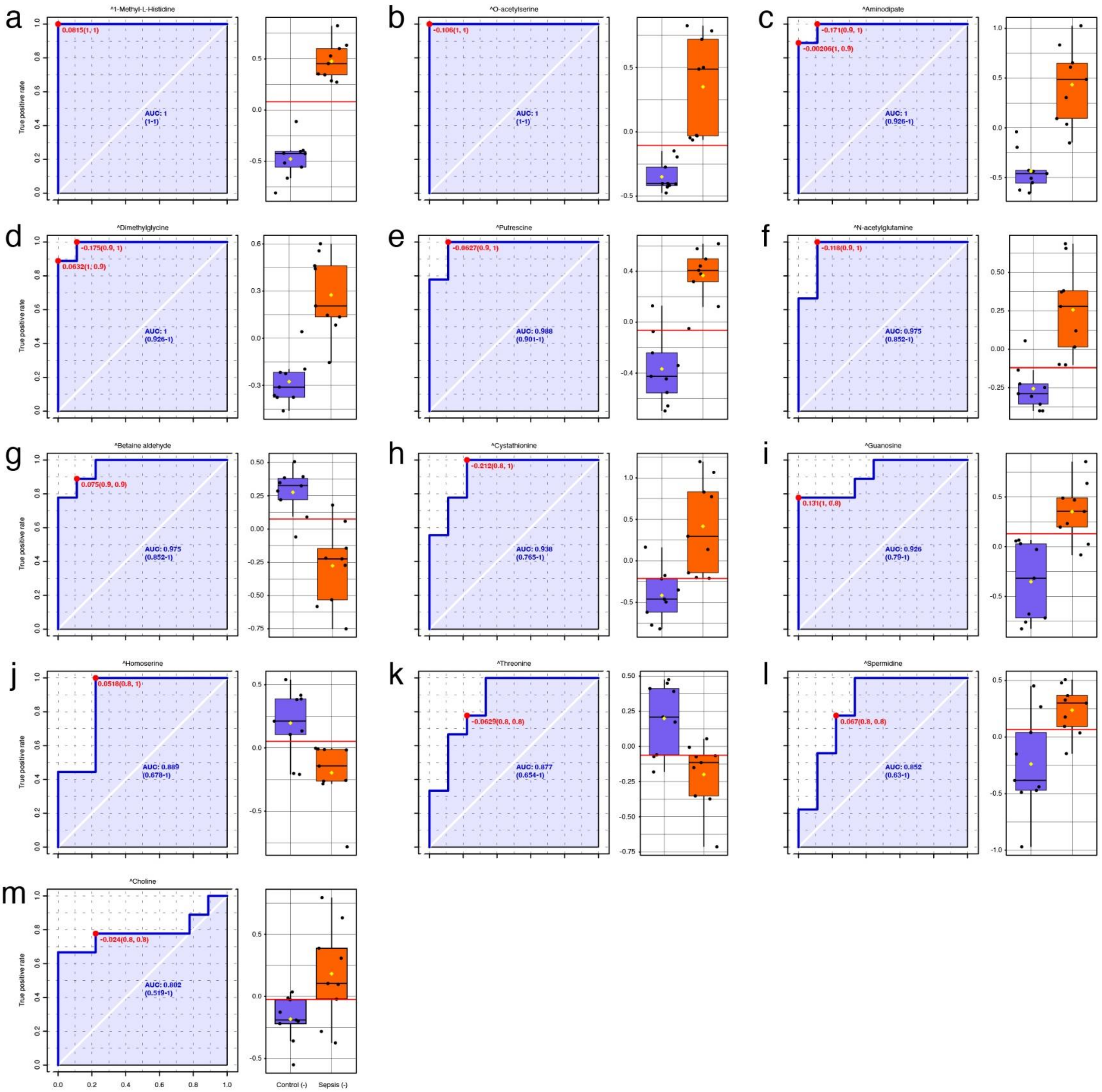

**Supplemental Figure 24. Targeted metabolomics-based overlapping biomarker candidates with high diagnostic performance in both cerebellum and plasma.** This figure displays 13 targeted metabolite features with strong diagnostic potential (AUC $\geq 0.8$) that were altered in both cerebellar and plasma samples considering the disease state comparison, Sepsis (-) v. Control (-). Panels **a-i** display receiver operating characteristic (ROC) curves for each overlapping metabolite candidate, with AUC values shown to reflect classification performance. Box plots are displayed to the right of each metabolite's ROC curve, illustrating the relative average concentrations of each metabolite by condition. Where known, the best diagnostic threshold is displayed by a black horizontal line on the box plots. Caret symbols (^) denote a Level 1-2 confidence identification from core facility panel confirmed by reference standard[25].
